# Mechanism of NanR transcriptional activation of sialic acid metabolism in *Streptococcus pneumoniae*

**DOI:** 10.64898/2025.12.10.693482

**Authors:** David M. Wood, Ashleigh S. Johns, Zachary D. Tillett, Heather L. Shearer, Santosh Panjikar, Yee-Foong Mok, Michael D.W. Griffin, Timothy M. Allison, Rachel A. North, Paul E. Pace, Borries Demeler, Mark B. Hampton, Renwick C.J. Dobson, Christopher R. Horne

## Abstract

In *Streptococcus pneumoniae*, the RpiR transcriptional regulator NanR (*Sp*NanR) senses sialic acid in the environment and upregulates transcription of the *nan* and *sia*A operons to increase uptake and metabolism of sialic acid. The molecular basis of this activation is unknown. Here, we demonstrate that *Sp*NanR binds *N*-acetylmannosamine-6-phosphate, a metabolite of sialic acid catabolism. *Sp*NanR exists in a dimer-tetramer equilibrium, and *N*-acetylmannosamine-6-phosphate binding strongly stabilizes the tetramer. Crystal structures and site-specific substitutions demonstrate that *N*-acetylmannosamine-6-phosphate bridges and stabilizes the *Sp*NanR tetramer. *Sp*NanR binds its DNA recognition sequence with nanomolar affinity. Notably, the effector *N*-acetylmannosamine-6-phosphate does not affect the affinity of *Sp*NanR for DNA. The DNA binding domains are not structurally coupled to the sugar isomerase domains, explaining why *N*-acetylmannosamine-6-phosphate binding does not affect DNA binding. Structural analysis reveals that sequence specificity arises through distortion of B-DNA and an unusual π-stack formed by two arginine residues in the minor groove, while affinity is driven by backbone contacts. We propose a mechanism by which *S. pneumoniae* regulates sialic acid metabolism, consistent with our biophysical experiments and *in vivo* regulatory behavior. These findings define a unique activation mechanism for an RpiR regulator and provide new insights into carbohydrate-responsive gene regulation in pneumococci.

## Introduction

Transcriptional gene regulation enables a cell to tailor its gene expression in response to a molecular signal. For example, when a cell detects changes in the environmental nutrient levels, genes encoding the appropriate machinery are induced, while genes that are not required are repressed. In this way, bacteria optimize growth and improve their fitness within a new environment by optimizing growth. However, the molecular mechanisms that govern such decisions are often unknown. Here, we define how a Ribose phosphate isomerase Regulator (RpiR) protein controls the expression of genes required for the uptake and utilization of an alternative carbohydrate in *Streptococcus pneumoniae*.

The ability to metabolize alternative carbohydrates, such as sialic acids, is a known virulence factor for bacterial colonization and pathogenesis (1–4). Sialic acids are amino monosaccharides commonly found conjugated to the surface of mammalian cells (5), where they serve as markers of ‘self’ for the human immune system (6,7). Some bacteria, including *S. pneumoniae*, cleave, import and metabolize sialic acids derived from mammalian glycoconjugates [(8,9), reviewed (10–12)]. Disrupting the genes responsible for sialic acid import (4,13,14) or catabolism (15–17) decreases microbial pathogenesis. Bacteria import sialic acids for several reasons: they provides precursors (mannosamine and glucosamine) for cell wall synthesis (18); they can be catabolized *via* glycolysis for chemical energy and serve as a source of carbon and nitrogen precursors (19); and, in some species, repurposed sialic acids are displayed on their own cell surface to evade the immune system through molecular mimicry (7,20,21).

The ability to metabolize sialic acids by *S. pneumoniae* is conferred by the *nan* gene cluster (22,23). While it varies slightly amongst pneumococcal strains, the *nan* gene cluster typically comprises two *nan* operons and a *sia*A operon (**Figure 1A**). *Nan* operon-I encodes catabolic enzymes that convert sialic acid to *N-*acetylglucosamine-6-phosphate through a series of steps (24–28) (**Figure 1B**), while *nan* operon-II encodes processing enzymes and an ABC transporter. The *sia*A operon encodes a sialidase responsible for cleaving extracellular, host-derived sialic acids conjugated on mammalian cell surfaces (23).

**Figure 1.**
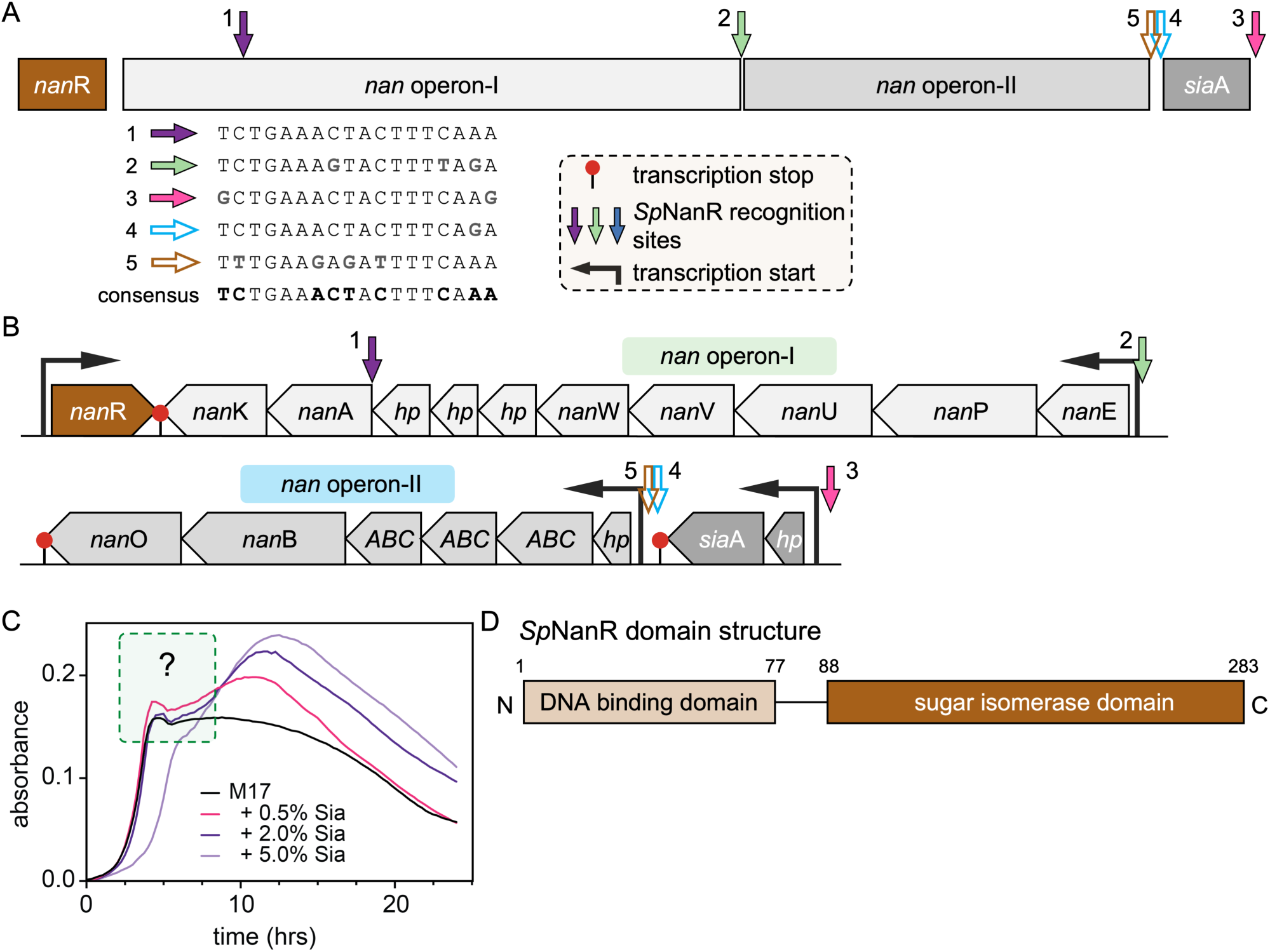
Organization of the *nan* gene cluster and *Sp*NanR recognition sequences. (**A**) The *nan* gene cluster. Three transcripts are defined by boxes and the location of the three *Sp*NanR recognition sequences reported are annotated (solid arrows). We also identified two additional sites (open arrows). The *Sp*NanR recognition sequences vary, but the consensus sequence is ^5’^TCTGAAACTACTTTCAAA^3’^. (**B**) The *nan* cluster transcripts show the location of each gene, including *nan*R (*Sp*NanR, brown). Black arrows are located at transcription start sites and red lollipops denote transcription stop sites. Colored arrows show the location of the *Sp*NanR recognition sequences annotated from strain D39, NC_008533.2 (as in A). Nan operon-I (in the direction of transcription): *nan*E *(N*-acetylmannosamine-6-phosphate 2-epimerase), *nan*P (PTS, IIBC components), *nan*U (ABC sugar-binding protein), *nan*V (ABC transporter permease), *nan*W (ABC carbohydrate transporter), *hp* (hypothetical proteins), *nan*A (*N*-acetylneuraminate lyase), *nan*K (*N*-acetylneuraminate kinase). Nan operon-II: *hp* (hypothetical protein), *ABC* (ABC substrate-binding protein, transporter permease, carbohydrate transporter), *nan*B (neuramidase), *nan*O (oxidoreductase). *sia*A operon: *hp* (hypothetical protein) and *sia*A (sialidase, sometimes annotated as *nan*A). (**C**) Growth of *S. pneumoniae* on M17 (no lactose) supplemented with PBS (control, black) or increasing quantities of sialic acid (%w/v). Between hours five and eight and only when sialic acid is included, there is lag phase and a secondary logarithmic growth phase. The delayed log phase observed in the 5% sialic acid concentration (light purple) could be due to toxicity. (**D**) Domain structure of *Sp*NanR, as predicted by InterPro (29).

The proteins *Sp*NanR and the catabolite control protein A (CcpA) regulate transcription of the genes that allow for sialic acid utilization in *S. pneumoniae* (22,23,30). *Sp*NanR is predicted to recognize an 18 base pair near palindromic DNA sequence, which has been identified at three sites within the *nan* gene cluster (**Figure 1A-B**, solid arrows) (22). In addition to these, we identified two further putative recognition sites (open arrows) with a consensus sequence of ^5’^TCTGAAACTACTTTCAAA^3’^ across these sites (**Figure 1A**). In the presence of sialic acid but absence of glucose, expression of *nan* operon-I and the *sia*A operon is significantly increased; this effect is abolished upon deletion of *Sp*NanR (22) indicating that *Sp*NanR functions as a transcriptional activator. However, in the presence of glucose and sialic acid, transcription of these operons is not induced. Repression is attributed to the binding of the CcpA protein to the −35 box, which blocks RNA polymerase. CcpA recognizes a ‘catabolite repression element’ (consensus is a pseudo palindrome ^5’^tTGAAAGtGtTTaCAa^3’^ (30) that is found within the promoter regions of *nan* operon-I and the *sia*A operon. In this way, *S. pneumoniae* exhibits classical diauxic growth in media containing both glucose and sialic acid (**Figure 1C**). The resulting growth curve is biphasic, with an initial growth phase followed by a lag phase before growth resumes (31,32). Increasing the concentration of sialic acid enhances the magnitude of the second growth phase, suggesting that glucose is preferentially consumed first, followed by sialic second (**Figure 1C**). Here, we examine the molecular mechanism underlying this phenotype.

*Sp*NanR is an RpiR transcriptional regulator comprising two domains (**Figure 1D**). The N-terminal DNA-binding domain is a well-conserved helix-turn-helix motif, defined by a tri-helical structure where the third α-helix is often referred to as the ‘recognition helix’ and interacts directly with the DNA major groove (33). The C-terminal domain is a sugar isomerase domain, a common feature in regulators of sugar-phosphate metabolism (34). *N*-Acetylmannosamine or *N*-acetylmannosamine-6-phosphate, which are both sialic acid derivatives, are thought to modulate transcription of the *nan* gene cluster, presumably through binding the *Sp*NanR sugar isomerase domain (22,23).

RpiR transcription regulators control a variety of processes within bacteria (35–39), but are understudied at the mechanistic and structural level. Five independent domain structures have been determined experimentally: two for the N-terminal DNA-binding domain (PDB ID: 3IWF, 203F; both unpublished) and three for the sugar isomerase domain only (PDB ID: 3SHO, unpublished; 3CVJ, unpublished; 7EN6 (40)]. In addition, there are two full-length RpiR structures: NanR protein from *Vibrio vulnificus* (*Vv*NanR) in complex with *N*-acetylmannosamine-6-phosphate [PDB ID: 4IVN (41)] and the recently reported *Pseudomonas aeruginosa* RccR regulator of carbon metabolism [PDB ID: 8JU9 (42,43)]. To date, no crystal structures of full-length RpiR bound to DNA or in the apo form (*i.e.*, without effector) have been reported, representing a key knowledge gap in understanding the molecular mechanism of RpiR transcriptional regulators.

Here, we report on the molecular mechanism of sialic acid regulation in *S. pneumoniae*. We identify the effector that binds *Sp*NanR to regulate expression of the *nan* and *sia*A operons. *N-*Acetylmannosamine-6-phosphate binding modulates the oligomeric state of *Sp*NanR but does not affect the affinity of *Sp*NanR for DNA. We report the first full-length structures of the RpiR family transcriptional regulator *Sp*NanR, including the unbound (apo) *Sp*NanR crystal structure, *Sp*NanR in complex with *N*-acetylmannosamine-6-phosphate, and *Sp*NanR in complex with its cognate DNA recognition sequence. Together, these results allow us to propose a molecular mechanism for the interplay between sialic acid, operon structure and the transcriptional regulator *Sp*NanR, which affords a competitive advantage to the bacterium *S. pneumoniae* during host colonization.

## Results

### The sialic acid pathway intermediate N-acetylmannosamine-6-phosphate binds SpNanR

*In vivo* studies demonstrate that mannose and sialic acid promote the expression of the *S. pneumoniae nan* operon (22,23) and that these sugars, or an intermediate metabolite within the sialic acid catabolic pathway, are the likely effector that binds to and modulates the function of *S. pneumoniae* NanR (*Sp*NanR). Recombinant, full-length *Sp*NanR (**Supplementary Table 1**) was expressed in *E. coli* and purified (**Supplementary Figure 1A**). To identify the *Sp*NanR effector we screened sialic acid pathway intermediates and precursors for binding using differential scanning fluorimetry (**Figure 2A**, **Supplementary Table 2**, **Supplementary Figure 2**). Only *N*-acetylmannosamine-6-phosphate (manNAc-6-P in Figure 2A) increases the melting temperature of *Sp*NanR (50.5 °C) compared to *Sp*NanR alone (46.4 °C), suggesting that binding is specific. Notably, *N*-acetylglucosamine-6-phosphate (glcNAc-6-P), which differs only by the stereochemistry of the acetyl group at the C2 position, and *N*-acetylmannosamine (manNAc), which is missing the phosphate, do not alter the melting temperature. The dissociation constant (*K*_D_) for *N*-acetylmannosamine-6-phosphate is 20.4 ±6.0 μM (*n* = 3), measured using isothermal titration calorimetry (**Figure 2B**, **Supplementary Figure 3**). The binding coefficient (*N*) is 1.19 ±0.04, suggesting that one *N*-acetylmannosamine-6-phosphate molecule binds each *Sp*NanR monomer. Together, these data support that *N*-acetylmannosamine-6-phosphate is the effector that modulates *Sp*NanR.

**Figure 2.**
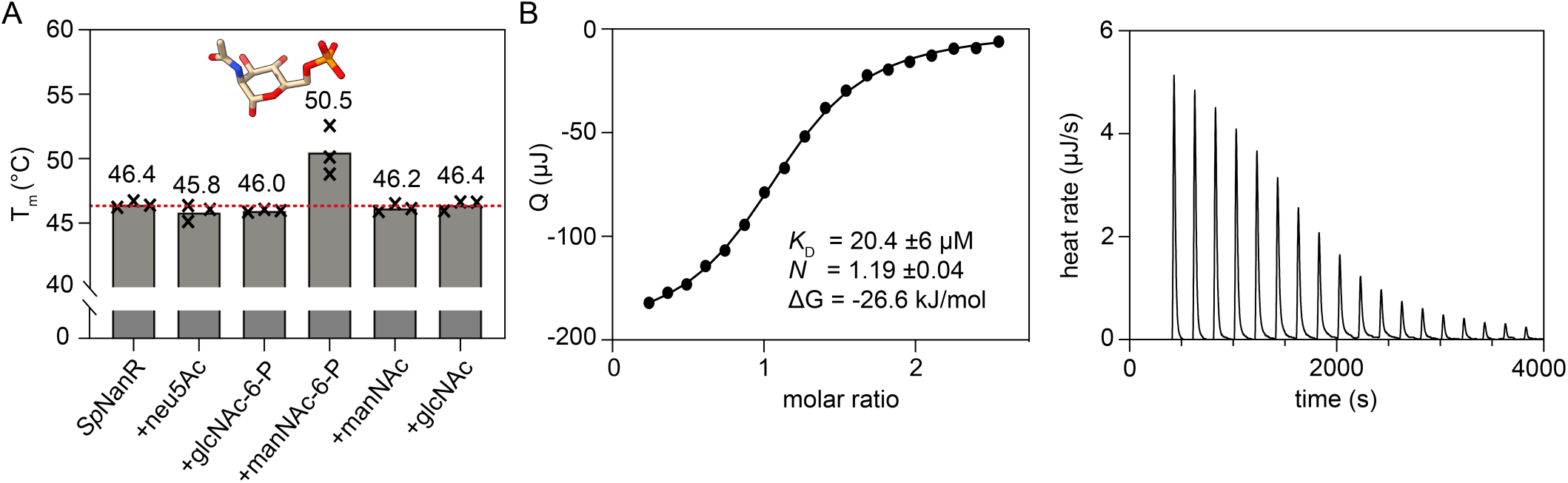
*N*-Acetylmannosamine-6-phosphate binds to *Sp*NanR. (**A**) The mean melting temperature (T_m_, °C) for *Sp*NanR in the presence of sialic acid derivatives. Measurements are in triplicate (black crosses). *N*-acetylmannosamine-6-phosphate (manNAc-6-P, displayed as sticks) shows a significant increase in stability, +4.1 °C, relative to *Sp*NanR alone (red dash). (**B**) Representative isothermal titration calorimetry data demonstrating binding of *N*-acetylmannosamine-6-phosphate (left panel). The thermodynamic parameters are an average of three independent experiments (**Supplementary Figure 3**). The heat rate (μJ/s) with corrected baseline for injections of *N*-acetylmannosamine-6-phosphate is shown (right panel).

### N-Acetylmannosamine-6-phosphate modulates the oligomeric state of SpNanR

Recombinant *Sp*NanR elutes as a single but highly asymmetric peak by size exclusion chromatography (**Supplementary Figure 1A**), indicating it may form multiple oligomeric species. To assess the oligomeric state of *Sp*NanR in solution, we undertook analytical ultracentrifugation sedimentation velocity studies across a broad range of *Sp*NanR concentrations with or without *N*-acetylmannosamine-6-phosphate. We then analyzed the data using 2-dimensional spectrum analysis (2DSA) (44) and Monte Carlo analysis (2DSA-MC) (45), followed by van Holde-Weischet analysis (46), according to methods described in (47).

In the absence of *N*-acetylmannosamine-6-phosphate, the van Holde-Weischet analysis (**Figure 3A**, open circles) shows an increasing S value as a function of protein concentration (from ∼4.4 S to ∼5.8 S) and a shape that is diagnostic of a mass action driven self-association (48). Differential sedimentation coefficient distributions based on 2DSA-MC across a broader concentration range (4.3–227.7 μM) shows a smaller peak ∼4.4 S at low concentrations that shifts to a larger peak at ∼5.8 S as the protein concentration increases (**Supplementary Figure 4A** and **Supplementary Table 3**). The species at ∼4.4 S evident at the lowest concentration (4.3 μM, **Supplementary Figure 4A**, dark blue trace) has a mass of 64.1 kDa determined from the 2DSA-MC distribution with a *f/f*_o_ = 1.31, which is consistent with an *Sp*NanR dimer (calculated mass from the sequence is 65.2 kDa). Native mass spectrometry corroborates the dimeric *Sp*NanR species and that the larger species evident at high protein concentrations is a tetramer (**Supplementary Figure 6A**).

**Figure 3.**
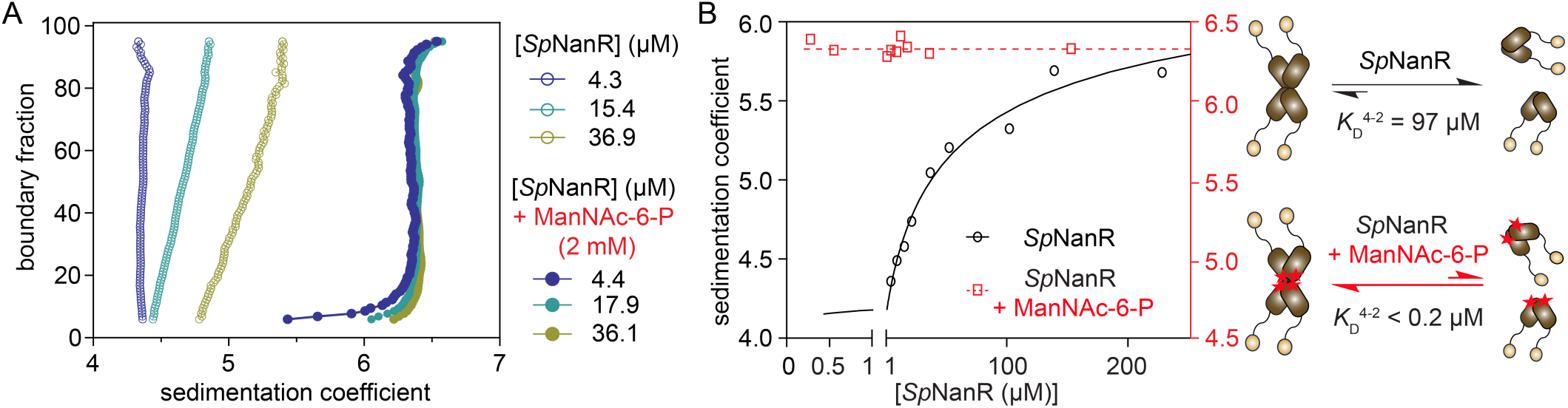
The oligomeric state of *Sp*NanR is modulated by *N*-acetylmannosamine-6-phosphate. (**A**) *Sp*NanR in the absence of *N*-acetyl mannosamine-6-phosphate (open circles) self-associates between two species determined to a dimer and tetramer. However, across the same concentration range in the presence of *N*-acetyl mannosamine-6-phosphate (filled circles), only the tetramer is evident at ∼6.3 S. The van Holde-Weischet plots (46) are generated from analytical ultracentrifugation sedimentation velocity data across a range of *Sp*NanR concentrations (4.3–36.9 μM) with or without *N*-acetyl mannosamine-6-phosphate (2 mM). (**B**) In the absence of *N*-acetylmannosamine-6-phosphate (black, left axis), the tetramer-dimer dissociation constant (*K*_D_^4-2^) is 97 μM (68% confidence interval = 41–234 μM). In the presence of *N*-acetyl mannosamine-6-phosphate (2 mM, red, right axis), even at the lowest concentration tested (0.27 μM), *Sp*NanR is still tetrameric with a *K*_D_^4-2^ <0.2 μM, demonstrating that *N*-acetylmannosamine-6-phosphate significantly stabilizes the tetrameric form. The weight-averaged sedimentation coefficients reported in the 2DSA-MC analysis (**Supplementary Figures 4A** and **5A**) are plotted as a function of *Sp*NanR monomer concentration and fitted to a dimer-tetramer self-association model using Sedphat (49).

In the presence of *N*-acetylmannosamine-6-phosphate (2 mM), however, the van Holde-Weischet analysis shows a single species at ∼6.3 S even at the low *Sp*NanR concentration of 4.4 μM (**Figure 3A**, filled circles) that is also evident in the 2DSA distribution (**Supplementary Figure 5A** and **Supplementary Table 3**). This species at ∼6.3 S ([*Sp*NanR] = 36.1 μM, **Supplementary Figure 5A**, pink trace) has a mass of 129.3 kDa (*f/f*_o_ = 1.42), which is consistent with a tetramer of *Sp*NanR (calculated mass is 130.4 kDa).

The dissociation constants (*K*_D_^4-2^) were estimated by plotting the weight-averaged sedimentation coefficients against protein concentration and fitting this to a dimer-tetramer self-association model (50) (**Figure 3B**). In the absence of *N*-acetylmannosamine-6-phosphate, the *K*_D_^4-2^ is 97 μM. In contrast, in the presence of *N*-acetylmannosamine-6-phosphate (2 mM) the *K*_D_^4-2^ could not be determined as the tetrameric species dominates even at 0.27 μM, but does provide an upper boundary for the *K*_D_^4-2^ of *Sp*NanR + *N*-acetylmannosamine-6-phosphate as < 0.2 μM, demonstrating that *N*-acetylmannosamine-6-phosphate binding results in at least a 2,000-fold increase in the affinity for the tetramer (illustrated in **Figure 3B**, right panel).

Together, these studies demonstrate that *Sp*NanR self-associates between dimeric and tetrameric species, and that the effector *N*-acetylmannosamine-6-phosphate stabilizes the tetrameric state.

### SpNanR binds the nanE recognition sequence with nanomolar affinity and this is unaffected by N-acetylmannosamine-6-phosphate

The *S. pneumoniae* D39 genome contains five 18 base pair near-palindromic *Sp*NanR recognition sequences that differ slightly across the *nan* cluster (**Figure 1A**). We determined the binding affinity of *Sp*NanR to three recognition sequences located [1] upstream of *nan* operon-I (site 2 in **Figure 1A,B**), [2] upstream of the *nan*A gene (site 1), and [3] the cryptic site downstream of the *sia*A operon (site 4) using microscale thermophoresis. The recognition sequence in the promoter region of the *sia*A operon is highly similar to that of *nan* operon-I and was therefore not studied separately. Additionally, the putative site in the promoter region of *nan* operon-II was not analyzed because it is nonfunctional (22). Despite their subtle differences in sequence, *Sp*NanR has a similar affinity for all three (483–752 nM, **Supplementary Figure 7**). We chose to focus on the 18 base pair *nan*E recognition sequence (^5’^TCTGAAAGTACTTTTAGA^3’^) for subsequent studies because *nan* operon-I has been demonstrated to be activated by *Sp*NanR *in vivo* (22).

To determine the dissociation constant for *Sp*NanR and the *nan*E recognition sequence and define the effect of *N*-acetylmannosamine-6-phosphate, we titrated *Sp*NanR (0.019–5 μM) against FAM-labeled DNA (50 nM) with or without *N*-acetylmannosamine-6-phosphate and monitored the change in sedimentation coefficient using fluorescence detection analytical ultracentrifugation (**Figure 4A**). The fraction of DNA bound to *Sp*NanR was determined, and in both cases the data best fitted a simple binding model with a Hill coefficient (**Figure 4B**). The dissociation constant (*K*_D_) for *Sp*NanR binding to the *nan*E recognition sequence is 476 ± 75 nM, which is consistent with the microscale thermophoresis experiment (**Supplementary Figure 7**). Surprisingly, a similar dissociation constant was observed in the presence of *N*-acetylmannosamine-6-phosphate (2 mM) (549 ± 84 nM), indicating that effector binding does not significantly alter *Sp*NanR’s DNA binding affinity for the *nan*E recognition sequence and simply induces *Sp*NanR tetramer formation. We corroborated these results using multi-wavelength sedimentation velocity experiments (**Figure 4C** and **Supplementary Figure 10**), which allowed us to monitor the DNA and *Sp*NanR sedimentation signals independently without the need for fluorescent dye. These data demonstrate that *Sp*NanR binds the *nan*E recognition sequence as both a dimer and tetramer, and that the dissociation constant for this interaction is similar with or without *N*-acetylmannosamine-6-phosphate, since the fraction bound is unchanged with the addition of the effector.

**Figure 4.**
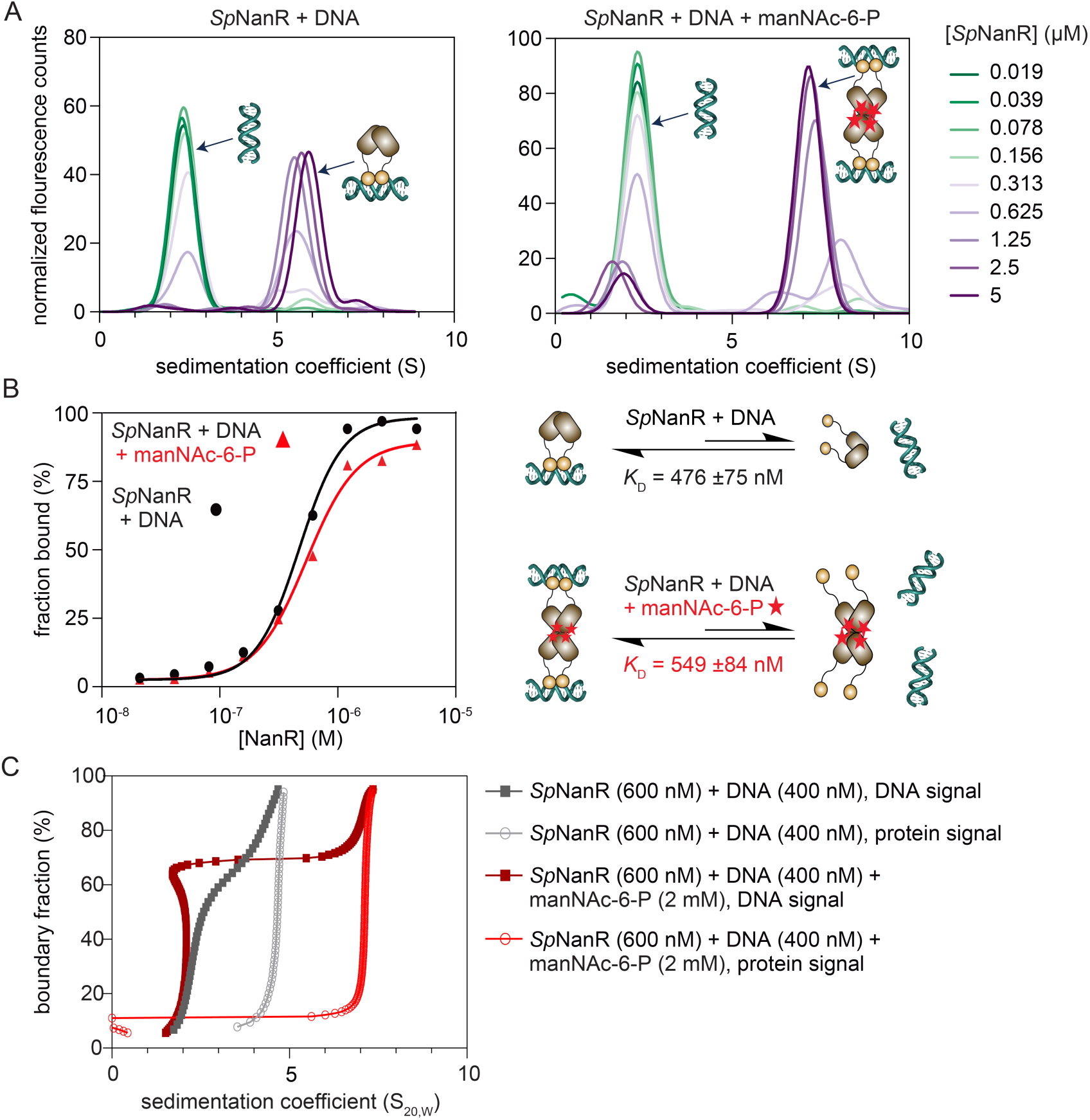
Sedimentation velocity analysis of *Sp*NanR binding to cognate DNA sequence. (**A**) Sedimentation distributions of fluorescently labeled DNA (*nan*E recognition sequence ^5’^TCTGAAAGTACTTTTAGA^3’^, 50 nM) following titration of *Sp*NanR (0.019–5 μM). The sedimentation data were evaluated by the two-dimensional spectrum analysis (2DSA) (44) followed by genetic algorithm analysis (51). 2DSA distributions are shown for *Sp*NanR + DNA (left) and *Sp*NanR + DNA + *N*-acetylmannosamine-6-phosphate (2 mM, right). Fits to the data are in **Supplementary Figures 8** & **9**, fit statistics are in **Supplementary Table 5**. (**B**) The fraction of DNA bound to protein from A (*Sp*NanR + DNA, black line; *Sp*NanR + DNA + *N*-acetylmannosamine-6-phosphate; manNAc-6P, red line) was calculated from the data in A using UltraScan. These data were fit to a simple binding model with a Hill coefficient, plotted as a function of total *Sp*NanR concentration. The Hill coefficient (*n*) is 2.3 (95% CI = 1.75–3.11) in the absence *N*-acetylmannosamine-6-phosphate and 2.0 (95% CI = 1.50–2.68) in the presence of *N*-acetylmannosamine-6-phosphate. A schematic of the binding interactions and measured affinities for *Sp*NanR (brown) and the *nan*E recognition sequence (green), without or with *N*-acetylmannosamine-6-phosphate (red stars) is shown. (**C**) Integral distribution plots of deconvoluted multi-wavelength analytical ultracentrifugation data for *Sp*NanR (600 nM) with unlabeled *nan*E recognition sequence DNA (400 nM) in the absence (gray) or presence (red) of *N*-acetylmannosamine-6-phosphate (2 mM). The presence of *N*-Acetylmannosamine-6-phosphate does not alter the amount of *Sp*NanR bound to DNA. **Supplementary Figure 10** shows a concentration series, demonstrating that the *Sp*NanR^mon^:DNA stoichiometry approaches 2:1 in the absence of *N*-acetylmannosamine-6-phosphate and 4:1 in the presence of *N*-acetylmannosamine-6-phosphate.

The Hill coefficient is ∼2 in both cases, suggesting there are at least two equilibria with different dissociation constants operating in this experiment. We considered two possibilities: that there is non-specific binding between the charged patches on the protein (presumably the DNA binding domain) and the negatively charged DNA; or a DNA molecule binds the DNA binding domains from two different *Sp*NanR dimers (or tetramers) and since the DNA sequence is not a perfect palindrome the affinities may be different, especially if the two dimers/tetramers sterically hinder binding to the second site. In any event, the data demonstrates that the effector *N*-acetylmannosamine-6-phosphate does not influence the binding affinity for the *nan*E recognition sequence for *Sp*NanR.

These data also allow us to estimate the stoichiometry of binding. As the *Sp*NanR concentration increases, the free DNA signal at 2.1 S decreases and a new peak at ∼5.7 S (**Figure 4A**, left) or 7.2 S (right) appears, which corresponds to the *Sp*NanR-DNA complex. In the absence of *N*-acetylmannosamine-6-phosphate, the peak at ∼5.7 S corresponds to a mass of 85.3 kDa, consistent with a *Sp*NanR dimer with one DNA molecule bound (76.4 kDa). In the presence of *N*-acetylmannosamine-6-phosphate, the peak at ∼7.2 S corresponds to a mass of 144.9 kDa, consistent with a *Sp*NanR tetramer bound to either one DNA oligomer (141.8 kDa) or two DNA oligomers (153.2 kDa). These stoichiometries were corroborated by native mass spectrometry, which identifies *Sp*NanR dimer with one DNA molecule and tetramer species bound to either one or two DNA molecules (**Supplementary Figure 6B**).

To summarize, *Sp*NanR binds each of the three *Sp*NanR recognition sequences across the nan operons with similar affinity and requires the 18 base pair near-palindromic sequence for optimal binding. *Sp*NanR binds DNA with nanomolar affinity, though this is not altered by *N*-acetylmannosamine-6-phosphate binding.

### Structural basis of effector-binding and oligomerization

We solved crystal structures of *Sp*NanR with and without *N*-acetylmannosamine-6-phosphate to define how the effector stabilizes the tetrameric form. The structure of *Sp*NanR with *N*-acetylmannosamine-6-phosphate was initially determined using a single wavelength anomalous diffraction strategy (**Table 1**). This was used as a template to solve subsequent structures by molecular replacement, including two further structures of *Sp*NanR with *N*-acetylmannosamine-6-phosphate (2.01 Å and 2.95 Å), apo-*Sp*NanR (2.55 Å), and *Sp*NanR complexed with DNA (next section).

**Table 1.** Data collection and refinement statistics for *Sp*NanR crystal structure. * This is a merged dataset described in **Supplementary Table 8**.

|  | *SpNanR* + manNAc-6-P + triiodide * | *SpNanR* + manNAc-6-P | *SpNanR* + manNAc-6-P | *SpNanR* | *SpNanR* + DNA |
| --- | --- | --- | --- | --- | --- |
| **Data Collection** |  |  |  |  |  |
| Wavelength (Å) | 1.458644 | 0.953649 | 0.953732 | 0.953739 | 0.953739 |
| Space group | *P*2<sub>1</sub> | *I*4 | *C*222 | *P*2<sub>1</sub>2<sub>1</sub>2<sub>1</sub> | *P*22<sub>1</sub>2<sub>1</sub> |
| Unit cell parameters [a, b, c (Å)] | 147.3, 84.4, 148.0<br>(β = 90.2°) | 147.6, 147.6, 82.3 | 116.9, 223.7, 52.7 | 53.2, 109.2, 202.4 | 57.9, 159.4, 171.8 |
| Resolution range (Å) | 49.3–2.65 (2.71–2.65) | 42.2–2.01 (2.08–2.01) | 47.7–2.95 (3.06–2.95) | 45.9–2.55 (2.64–2.55) | 46.0–2.97 (3.13–2.97) |
| No. of unique reflections | 207,452 (14,687) | 58,719 (5,722) | 133,994 (13,126) | 39,440 (3,859) | 30,952 (4,821) |
| Multiplicity (outer shell) | 21.0 (3.5) | 13.5 (13.8) | 8.9 (8.9) | 10.1 (10.1) | 4.9 (5.1) |
| Completeness (%) (outer shell) | 99.6 (95.3) | 99.8 (95.7) | 99.9 (99.9) | 97.6 (98.0) | 92.0 (99.8) |
| *I*/σ(*I*) (outer shell) | 13.0 (1.0) | 16.8 (1.6) | 7.4 (0.9) | 9.16 (0.77) | 7.9 (0.8) |
| CC<sub>(1/2)</sub> (outer shell) | 0.999 (0.498) | 0.999 (0.777) | 0.994 (0.355) | 0.998 (0.61) | 0.998 (0.250) |
| CC<sub>anom</sub> | 31(0) | - | - | - | - |
| *R*<sub>merge</sub> (outer shell) | 0.17 (1.04) | 0.080 (1.187) | 0.2461 (2.364) | 0.188 (2.469) | 0.120 (1.765) |
| **Refinement Statistics** |  |  |  |  |  |
| No. of reflections used | - | 58,704 | 15,052 | 38,521 | 30,262 |
| No. of protein atoms | - | 5,991 | 3,712 | 7,863 | 10,185 |
| No. of water molecules | - | 184 | 0 | 21 | 4 |
| No. of ligand atoms | - | 100 | 51 | 64 | 55 |
| *R*<sub>work</sub> (%) | - | 21.0 | 25.0 | 26.2 | 23.0 |
| *R*<sub>free</sub> (%) | - | 27.1 | 30.1 | 31.2 | 26.7 |
| Reflections used for *R*<sub>free</sub> | - | 2,832 (306) | 1,767 (100) | 1,898 (194) | 1,522 (154) |
| Ramachandran (favored, allowed, disallowed) | - | 98.2, 1.8, 0.0 | 96.29, 3.71, 0.0 | 96.6, 3.3, 0.1 | 95.1, 4.8, 0.1 |
| RMS bond lengths (Å) | - | 0.015 | 0.006 | 0.006 | 0.004 |
| RMS angles (°) | - | 1.490 | 0.87 | 0.83 | 0.64 |
| Overall B-factor (protein, waters) | - | 60.85 | 72.91 | 74.3 | 86.11 |
| Clashscore (PHENIX) | - | 9.34 | 13.0 | 18.9 | 10.1 |
| PDB id | - | 8TX9 | 8V4V | 8V5F | 9DQN |

The *Sp*NanR monomer comprises two domains connected by a linker region of 11 residues (**Figure 5A**). The N-terminal DNA-binding domain comprises helices α1–α5 and adopts the well-characterized helix-turn-helix motif, which is common throughout prokaryotic and eukaryotic DNA binding proteins (52). The C-terminal sugar isomerase domain comprises an α/β-fold, with five parallel β-strands forming the core flanked by five of eight α-helices creating an αβα sandwich structure. Alignment of ligand-free monomers, however, highlights the alternative conformational arrangements of the DNA-binding domains relative to the sugar isomerase domains, likely due to flexibility in the linker region (**Supplementary Figure 11A**). This linker flexibility is further evidenced by the observation that only the backbone of the linker could be confidently modeled in the apo structure (**Supplementary Figure 11B**). The five DNA-binding domains across the two structures align well (r.m.s.d. = 0.44–0.75 Å, **Supplementary Figure 11C**) and the C-terminal domains also show strong structural agreement across all structures (r.m.s.d. = 0.20–0.35 Å, **Supplementary Figure 11D**). Further, the sugar isomerase domain, including the placement of the ligand binding pocket, is conserved across the four RpiR transcriptional regulator structures reported (**Supplementary Figure 11E**). We note that this domain aligns well with the sugar isomerase domain of glucosamine-6-phosphate-synthetase (GlmS, PDB ID: 1MOR, r.m.s.d. = 2.6 Å, **Supplementary Figure 11F**), which is known to isomerize fructose-6-phosphate to glucosamine-6-phosphate (53) and likely is an evolutionarily related domain.

**Figure 5.**
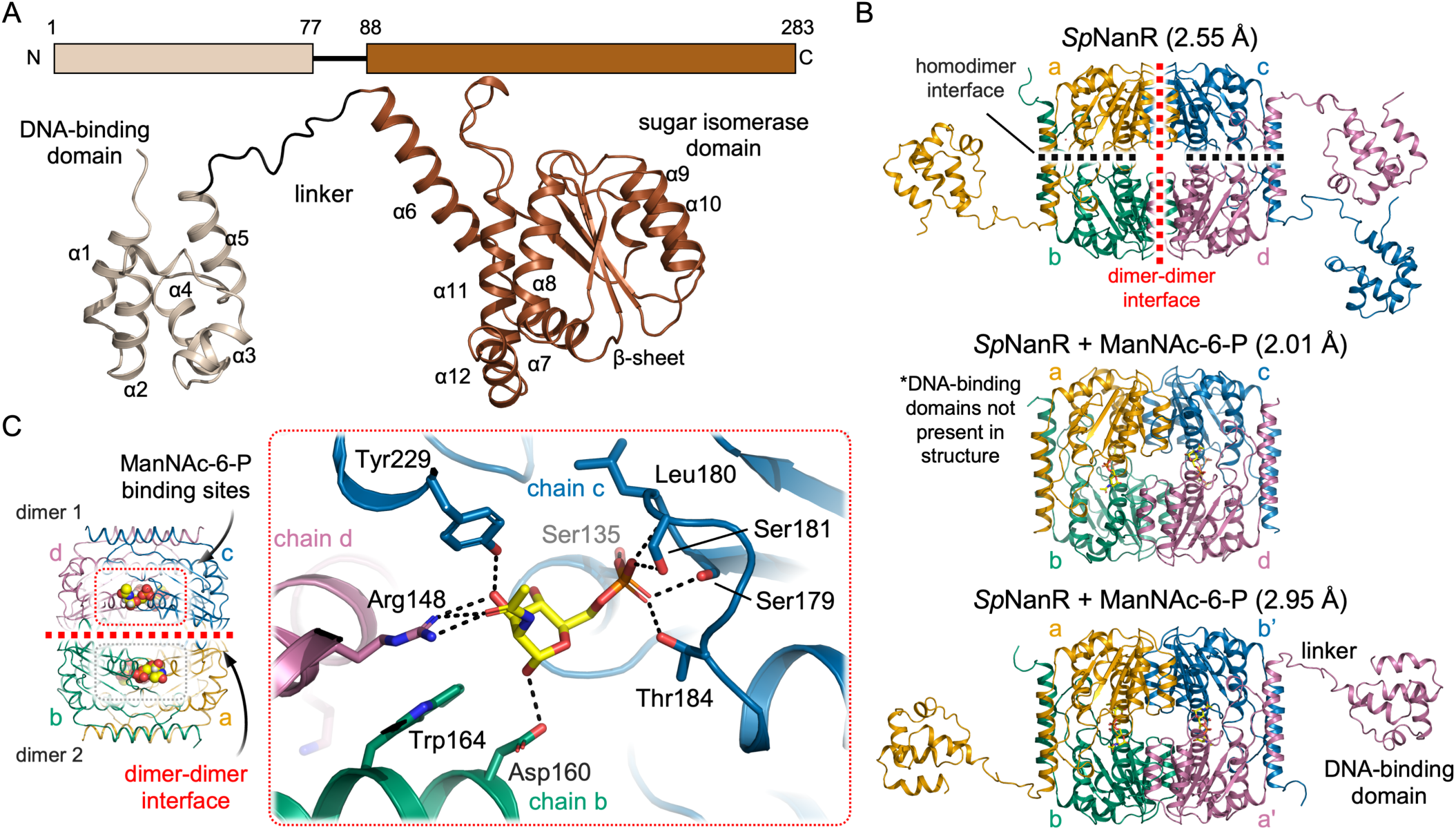
Structure of *Sp*NanR and the *N*-acetylmannosamine-6-phosphate binding site. (**A**) The *Sp*NanR monomer is made up of two domains; an N-terminal DNA-binding domain (light brown) and a C-terminal sugar isomerase domain (dark brown) fused by a flexible 11 amino acid linker region (black). (**B**) Crystal structures of *Sp*NanR assemble into a tetramer with and without *N*-acetylmannosamine-6-phosphate. Each monomer is coloured and annotated a-d. Two interfaces are formed: a dimer interface (black dotted lines) and a dimer-dimer interface (red dotted lines), calculated *via* PDBePISA. (**C**) *N*-Acetylmannosamine-6-phosphate binds close to and bridges across the dimer-dimer interface (dotted lines).

The *Sp*NanR tetramer comprises a ‘dimer-of-dimers’ formed through two extensive interfaces involving the sugar isomerase domains: a dimer interface between monomers a/b, or c/d (**Figure 5B**, black dashed line, buried surface area = ∼855 Å^2^); and a dimer-dimer interface between monomers a/c, or b/d, which forms the tetramer (red dashed line, buried surface area = ∼835 Å^2^). Neither the DNA binding domains nor the linker regions participate in oligomerization. Instead, the flexible linker regions (residues 77–88) position the DNA binding domains distal to the central sugar isomerase domain that forms the tetramer (**Figure 5B**, **Supplementary Figure 11B**), providing a rationale for why *N*-acetylmannosamine-6-phosphate does not affect the affinity between *Sp*NanR and DNA. If the DNA binding and sugar isomerase domains are independent, *i.e.*, not coupled, there is no way to transmit a structural signal induced by effector binding between the two domains. In contrast, the linker region is ordered in homologous RpiR regulators, allowing the DNA binding and sugar isomerase domains to interact (**Supplementary Figure 12**). This provides a mechanism for transmitting effector-induced structural changes between the domains and explains the corresponding changes in DNA-binding affinity. For example, in the *P. aeruginosa* RccR regulator (*Pa*RccR, PDB ID: 8JU9) the 16 residue linker forms a short α-helix that positions the DNA binding domains adjacent to the sugar isomerase domain (42,43).

We then verified whether this tetrameric configuration found in the crystal structure matches that in solution using small-angle X-ray scattering experiments (statistics in **Supplementary Table 9**). In-line size exclusion chromatography demonstrates that *Sp*NanR is polydisperse, with both tetrameric and dimeric populations (**Supplementary Figure 13A**). In contrast, *Sp*NanR in the presence of *N*-acetylmannosamine-6-phosphate (2 mM) is monodisperse and tetrameric (**Supplementary Figure 13B**). The maximum particle dimension (*D*_max_) of *Sp*NanR, with or without *N*-acetylmannosamine-6-phosphate was ∼165 Å from the *P*(*r*) analysis (**Supplementary Figure 13C**), which is consistent with the *D*_max_ measured from the crystal structures (∼160 Å). The Guinier analysis produced a linear plot for both datasets supporting that each sample is free from aggregation or inter-particle interference (**Supplementary Figure 13C**). Next, we tested whether the extended conformation seen in the crystal structures, or an alternative compact conformation where the DNA binding domains interact directly with isomerase domains—as observed in homologues—can be detected in solution. Using CRYSOL, the theoretical scattering profile of the extended conformation was best fit to the experimental SAXS data with *N*-acetylmannosamine-6-phosphate (*χ*^2^ value of 0.38; **Supplementary Figure 13D**), supporting that our structures are match the in solution structure and not an artifact of the crystal lattice. This extended conformation is further evidenced in the Kratky analysis, suggesting *Sp*NanR has two domains connected *via* a flexible linker, with or without *N-*acetylmannosamine-6-phosphate, which we observe in our electron density maps (**Supplementary Figure 11B**) and the bimodal charge state distributions by native mass spectrometry (**Supplementary Figure 6A**) (54,55).

Four *N*-acetylmannosamine-6-phosphate molecules bind in identical pockets near the dimer-dimer interface (**Figure 5C**, **Supplementary Table 3**). The tetrameric sugar isomerase domains from all three structures overlay closely (Cα r.m.s.d. = 0.30–0.46) and the dimer-dimer interface remains largely unchanged between the unbound and *N*-acetylmannosamine-6-phosphate bound structures (**Supplementary Figure 13E**), indicating that N-acetylmannosamine-6-phosphate does not stabilize the tetramer by rearranging this interface. Instead, each *N*-acetylmannosamine-6-phosphate molecule bridges three monomers across the tetrameric interface, thereby stabilizing the tetrameric state (**Figure 5C**). The phosphate group coordinates Ser135, Ser179, the Leu180 backbone, Ser181, and Thr184 from one monomer, while the sugar moiety interacts with Arg148 and Tyr229 from the second monomer within the dimer and with Asp160 from the third monomer across the dimer-dimer interface (**Figure 5C** inset and **Supplementary Figure 14A, Supplementary Table 6**). Arg148 coordinates the C2 *N*-acetyl and C3 hydroxyl groups, explaining why *Sp*NanR does not bind *N*-acetyl *gluc*osamine-6-phosphate (**Figure 2A**), whose *N*-acetyl group adopts a different stereochemical orientation Notably, Arg148 also participates in a cation-π-anion stacking interaction with Trp164 and Asp160 across the dimer-dimer interface.

We prepared recombinant *Sp*NanR variants containing site specific alanine substitutions to evaluate the contribution of binding site residues to effector binding and oligomerization. These include a triple mutant *Sp*NanR^S179A/S181A/T184A^, which probes phosphate coordination, and *Sp*NanR^R148A^ and *Sp*NanR^D160A^, which probes binding of the *N*-acetyl and sugar hydroxyl groups and the role of the cation-π-anion stacking interaction across the dimer-dimer interface (**Figure 5C**). First, we evaluated each variant’s thermal stability by differential scanning fluorimetry. All *Sp*NanR variants exhibited melting temperatures that were comparable with *Sp*NanR^WT^ (ranging from 49 to 51°C; (**Supplementary Figure 14B**), suggesting they fold correctly. Upon incubation with *N*-acetylmannosamine-6-phosphate, *Sp*NanR^WT^ showed an increase in thermal stability (from 50.8 °C to 63.0 °C), consistent with effector binding. *Sp*NanR^D160A^, which bridges the dimer-dimer interface, exhibited a comparable increase, indicating that this residue is less important for effector binding. Conversely, *Sp*NanR^S179A/S181A/T184A^ and *Sp*NanR^R148A^ showed only marginal changes in thermal stability in the presence of *N*-acetylmannosamine-6-phosphate, suggesting that effector binding is abolished.

We next tested whether these *Sp*NanR variants affected oligomeric state using sedimentation velocity experiments. As expected, *N*-acetylmannosamine-6-phosphate does not stabilize the tetrameric state of *Sp*NanR^S179A/S181A/T184A^, consistent with thermal stability data indicating that the effector does not bind this variant (**Supplementary Figure 14C**). Interestingly, *Sp*NanR^D160A^ exists exclusively as a dimer in the absence of *N*-acetylmannosamine-6-phosphate, whereas *Sp*NanR^WT^ shows both dimeric and tetrameric species in solution. This supports that *Sp*NanR^D160A^ destabilizes the tetrameric form (**Supplementary Figure 14D**). However, in the presence of *N*-acetylmannosamine-6-phosphate, this variant behaves similarly to *Sp*NanR^WT^, consistent with the observation that it still binds *N*-acetylmannosamine-6-phosphate and that this sufficiently stabilizes the tetrameric form. In contrast, *Sp*NanR^R148A^ remains exclusively dimeric at the same concentration as *Sp*NanR^WT^, even in the presence of *N*-acetylmannosamine-6-phosphate (**Supplementary Figure 14E**). This suggests that the cation-π-anion stacking interaction across the dimer-dimer interface is critical for tetramer formation and subsequent stabilization by *N*-acetylmannosamine-6-phosphate.

To summarize, crystal structures of *Sp*NanR with and without *N*-acetylmannosamine-6-phosphate reveal how *Sp*NanR oligomerizes, show how the effector bridges the dimer-dimer interface to stabilize the tetramer, and explain why effector binding does not affect DNA binding.

### SpNanR recognizes the palindrome through an ‘indirect readout’ mechanism

We determined the structure *Sp*NanR in complex with the *nan*E recognition sequence (^5’^TCT<u>G</u>AAAGTACTTT<u>T</u>AGA^3’^, underline representing a break in the palindrome) to define how *Sp*NanR engages a specific DNA sequence. Crystals of the complex diffract to 2.96 Å and the final structure has good global statistics (*R*_free_ = 26.7%) and geometry (**Table 1**). All amino acids (except three residues at the N- and C-terminal) and all nucleotides could be built into the electron density. This structure is without *N-*acetylmannosamine-6-phosphate, since its addition failed to yield crystals that diffracted despite significant effort. The asymmetric unit contains two *Sp*NanR dimers, and two *nan*E recognition sequence DNA molecules (**Figure 6A**). The two dimers are loosely connected but offset in what appears to be crystal contacts at the same dimer–dimer interface used to form the tetramer.

**Figure 6.**
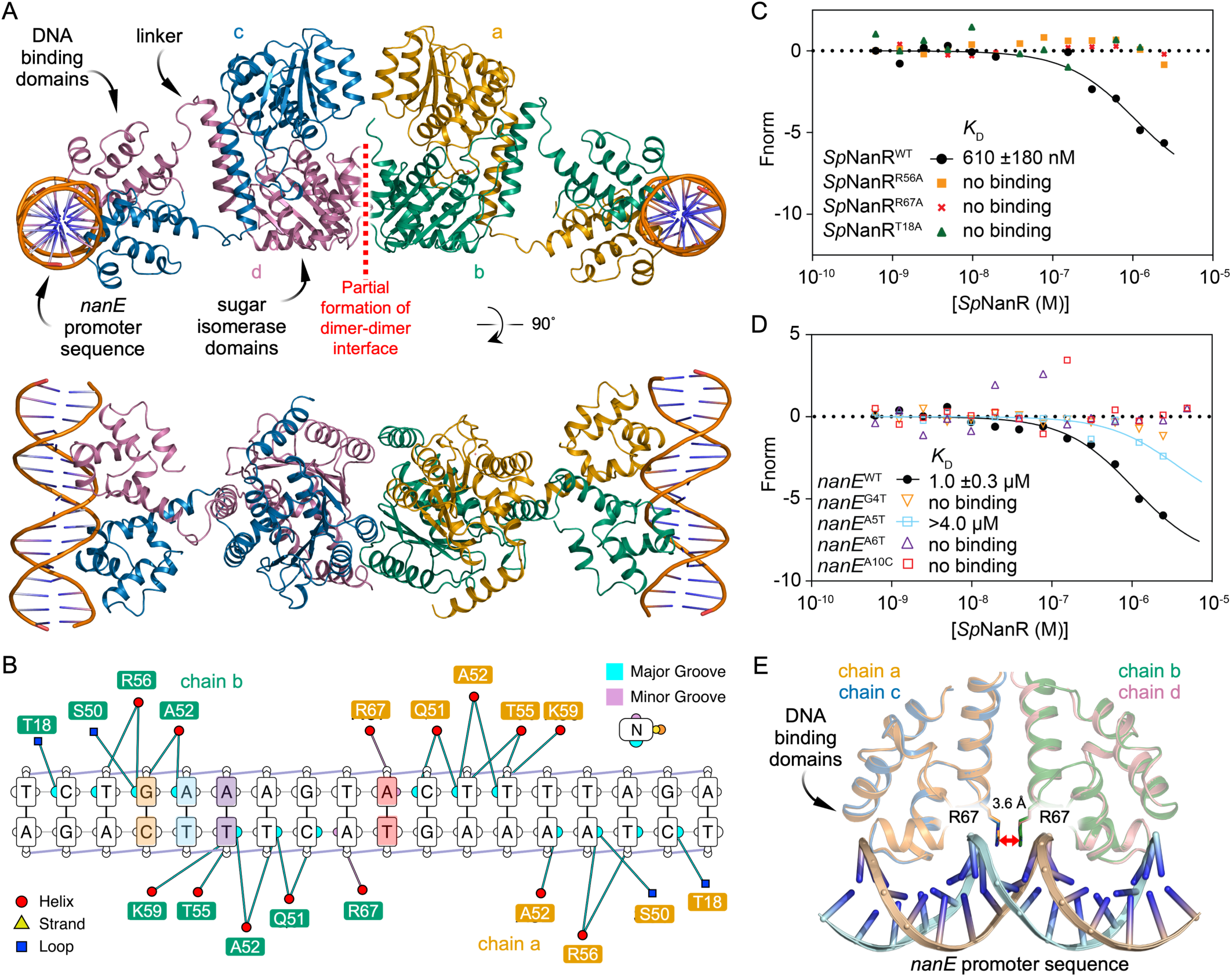
*Sp*NanR in complex with the *nan*E recognition sequence. (**A**) The protein-DNA complex crystallized with two *Sp*NanR dimers in the asymmetric unit, each bound to an 18 base pair dsDNA *nan*E recognition sequence (^5’^TCTGAAAGTACTTTTAGA^3’^). Each monomer is individually colored. **B**, *nan*E recognition sequence and *Sp*NanR amino acid interactions are depicted. Nucleotides that are mutated are colored (G4, A7, A6, and A10). Their binding affinity is tested in D. (**C**) Microscale thermophoresis results when titrating *Sp*NanR and variants (10–5,000 nM) against the native FAM-labeled *nan*E recognition sequence and mutated variants (20 nM). (**D**) Microscale thermophoresis results when titrating *Sp*NanR (10–5,000 nM) against native and mutated FAM-labeled *nan*E recognition sequences (20 nM). (**E**) View of the *Sp*NanR DNA-binding domains from the *Sp*NanR dimer binding to the 18 base pair *nan*E recognition sequence (sense strand = cyan, anti-sense = tan), highlighting the unique binding contributions from each R67 (orange label) residue in the two *Sp*NanR monomers.

In each dimer, the DNA binding domains engage one DNA molecule and are set away from the sugar isomerase domains by the disordered linker regions. Aligning the two dimers in the asymmetric unit shows that the dimeric sugar isomerase domains align well (r.m.s.d. = 0.47 Å over 361 Cα), however the DNA-binding domains and *nan*E recognition sequence DNA components are then offset (**Supplementary Figure 15A**). This suggests that although the DNA-binding domains interact with the *nan*E recognition sequence in the same manner, they are conformationally flexible relative the core sugar isomerase domains. An overlay of the DNA binding domains demonstrates that each dimer engages the DNA sequence in the same way (**Supplementary Figure 15B**). The *nan*E recognition sequence is in a B-DNA conformation (right-handed double helix with base pairs perpendicular to the helix axis) and when bound to *Sp*NanR remains straight rather than being bent or kinked.

Multiple experiments suggest that, in solution, *Sp*NanR bound to DNA and *N-*acetylmannosamine-6-phosphate adopts an extended and flexible conformation. Firstly, we collected small-angle X-ray scattering data with in-line size exclusion chromatography for the *Sp*NanR:*nan*E recognition sequence:*N-*acetylmannosamine-6-phosphate complex (**Supplementary Figure 16A**, **Supplementary Table 9**). The *D*_max_ from these data is ∼178 Å, which is consistent with the longest distance from the crystal structure (∼175 Å) and the Kratky plot has two distinct peaks consistent with multiple, independent domains (**Supplementary Figure 16B-C**). Using CRYSOL, we generated theoretical scattering profiles based on the crystal structure and additional models that include one or two DNA molecules. When compared against the small-angle X-ray scattering data, the model of the tetramer with two DNA molecules bound and extended away from the isomerase domains best fit the data (**Supplementary Figure 16D**). Next, native mass spectrometry data reports the presence of extended/flexible species when in the presence of DNA and *N-*acetylmannosamine-6-phosphate (**Supplementary Figure 6B**). Finally, attempts to determine the structure of the *Sp*NanR:*nan*E recognition sequence:*N*-acetylmannosamine-6-phosphate complex using single particle cryo-electron microscopy were unsuccessful due to particle heterogeneity (data not shown). The central sugar isomerase domains could be resolved, but the DNA-binding domains with DNA could not be resolved. Taken together, these data point to a flexible domain structure where the DNA binding domains are decoupled from the isomerase domains, as seen on the crystal structure.

*Sp*NanR engages the *nan*E recognition sequence in a way that explains the sequence specificity. Each monomer within the *Sp*NanR dimer interacts symmetrically with the major groove of the DNA through a conserved helix-turn-helix (HTH) motif. A survey of the interactions between *Sp*NanR and the the *nan*E recognition sequence is shown in **Figure 6B** and these are consistent across the two dimers in the asymmetric unit. As the *nan*E recognition sequence is an imperfect palindrome, the interactions with each DNA binding domain are similar but not identical. The recognition helix of *Sp*NanR (residues 50–61) nestles into the major DNA groove above the A—T track. Recognition and binding to specific nucleotide sequences often involves the formation of hydrogen bonds between amino acid sidechains and specific bases in the major groove—a process known as the ‘direct readout’ mechanism (56). Surprisingly, the recognition helix makes only one, long distance hydrogen bond with a base in the major groove and instead largely interacts with the sugar phosphate backbone (**Figure 6B**, **Supplementary Figure 15C**). Arg56 weakly interacts with the N7 nitrogen of the base at position 4 (guanidine) or position 15 (adenine) at ∼3.2 Å (average over four monomers in the asymmetric unit); these positions break the palindrome for the *nan*E recognition sequence. It also interacts with the phosphate backbone (distance ∼3.0 Å). Substituting Arg56 with alanine (*Sp*NanR^R56A^) results in no detectable binding to the *nan*E recognition sequence compared to *Sp*NanR^WT^ (**Figure 6C**, orange boxes).

Modeling suggests that mutating guanidine at position 4 or adenine at position 15 to the pyrimidines thymine or cytosine introduces a steric clash, partly explaining the sequence specificity at this position. Indeed, mutagenesis of the guanine at position 4 to thymine (*nan*E^G4T^) abolishes binding, as demonstrated by microscale thermophoresis (**Figure 6D**, orange open triangle). These results together suggest that although the interaction between Arg56 and the base is weak, it is essential for binding the *nan*E recognition sequence. Thr55 interacts exclusively with the backbone phosphate at position 12 of the sequence (∼2.7 Å), while Lys59 interacts with the backbone phosphate at position 13 (∼2.8 Å), and Ser50 interacts with the backbone phosphate at position 4 (∼2.6 Å). Ala52 points into the major groove, but the methyl sidechain does not make any direct connections with the bases. While water-mediated interactions are possible, none were observed in the major groove at 2.95 Å resolution, although waters interacting with DNA can be identified elsewhere. Outside of the recognition helix, the Thr18 sidechain and Leu20 mainchain amide interact with the phosphate backbone of the major groove. Substitution of Thr18 with alanine, *Sp*NanR^T18A^, abolishes binding (**Figure 6C**, green triangle). The mainchain amides of Tyr66 and Arg67 form hydrogen bonds with the phosphate of the nucleotide at position 12 (∼2.6 Å, and ∼3.5 Å, respectively).

A distinctive feature is the insertion of dual Arg67 sidechains from each *Sp*NanR monomer into the minor groove (**Figure 6E**). The guanidinium groups π-stack ∼3.6 Å apart (averaged over the two dimers) and each form a weak hydrogen bond with the N3 nitrogen of adenine at position 10 (∼3.5 Å). Both sidechains also interact with the backbone phosphates at position 10 (∼3.8 Å) and 11 (∼3.4 Å). Substitution of Arg67 with alanine abolishes binding (*Sp*NanR^R67A^, **Figure 6C**, red cross), evidence this interaction is essential for engaging the *nan*E recognition sequence. Arginine residues are commonly found to interact with the minor groove (57). A survey of helix-turn-helix DNA binding domains where an arginine interacts with the minor groove (57) reveals the π-stacked arginine conformation observed here as unique, expanding the repertoire of interactions that recognize the minor groove of B-DNA.

Aligning the *nan*E recognition sequence with an idealized B-DNA molecule with the same sequence shows it is distorted (r.m.s.d. = 1.4 Å aligning all 18 base pairs, **Supplementary Movie 1**), suggesting that recognition and specificity may be due to an ‘indirect readout’ mechanism (58). This is especially relevant because each monomer of *Sp*NanR makes only two long-range interactions with the base pairs: Arg56 (∼3.2 Å) and Arg67 (∼3.5 Å).

The minor groove width is significantly deformed upon *Sp*NanR binding (**Figure 7A**)—the width is compressed where engaging the recognition helix (8.6 Å, measured from the phosphate +4 bases of the opposing strand) and widened at nucleotides 9 and 10 where the two Arg67 insert (10.7 Å, **Figure 7B**). The minor groove width for an idealized B-DNA molecule is 11.7 Å. The base pair stacking is most altered at this position such that the TA dinucleotide (at positions 9 and 10) is significantly misaligned (helical twist is ∼50°, ideal is ∼36°, **Figure 7C**) and instead better stacked with the C and G bases on either side (helical twist is ∼26°). Arginine residues are enriched within the minor groove and commonly interact with adenine and thymine, especially where the minor groove narrows (57). Although mutating adenine 10 to cytosine in the *nan*E recognition sequence should be tolerated because it does not introduce a steric clash and offers hydrogen bonding interactions with Arg67, it too abolishes binding (*nan*E^A10C^, **Figure 6D**, open red box). TpA steps, as found in the *nan*E recognition sequence at positions 9/10 and 15/16, increase flexibility in B-DNA molecules (59–61), reviewed here (62). We reason that the mutation *nan*E^A10C^ breaks the TpA step and stabilizes the B-DNA structure such that *Sp*NanR cannot recognize or engage a distorted structure.

**Figure 7.**
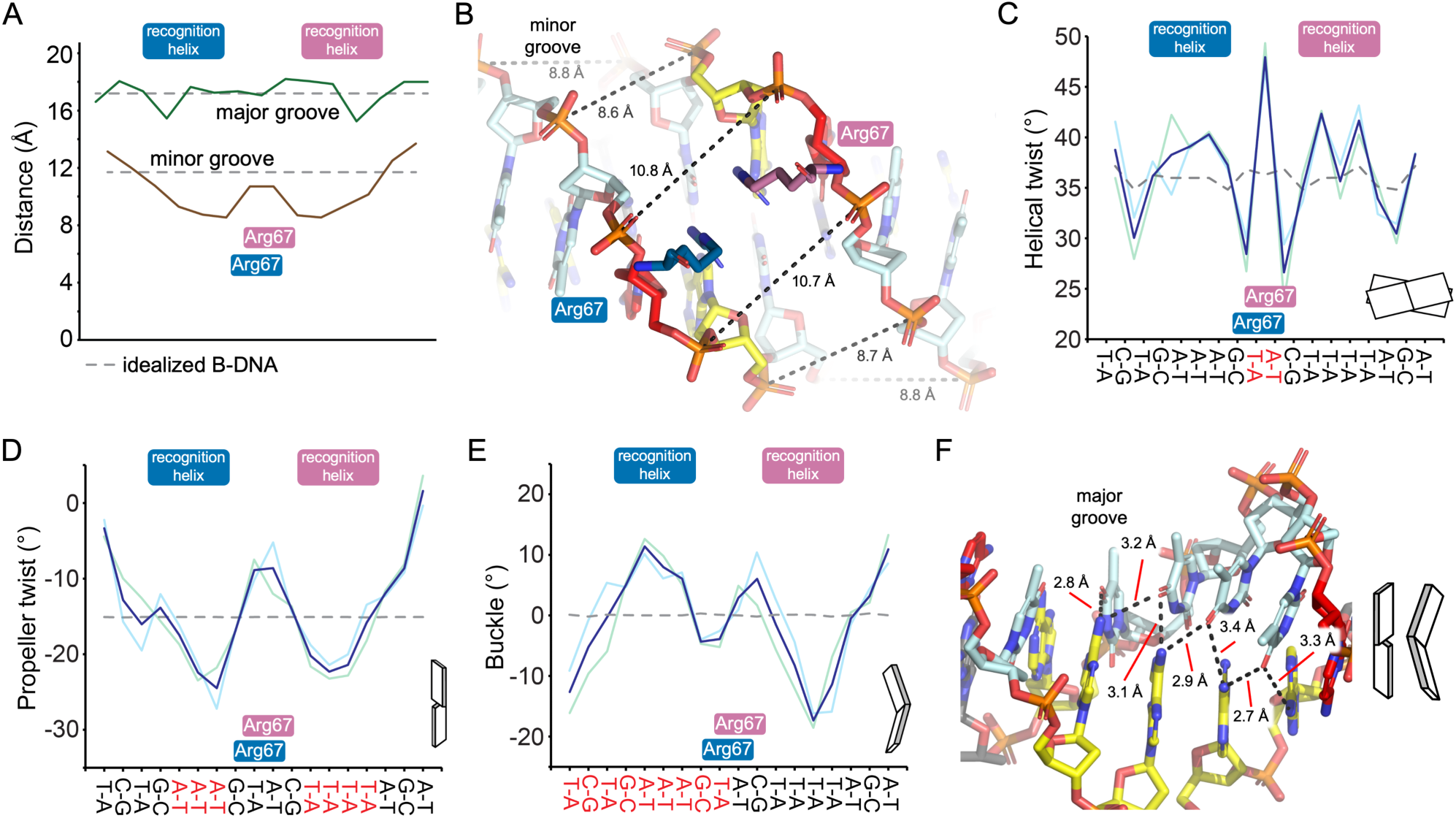
*Sp*NanR in complex with the *nan*E recognition sequence. **A**, Major and minor groove widths across the *nan*E recognition sequence bound to *Sp*NanR. **B**, The distances across the minor groove, measured from opposing backbone phosphate, at the point of Arg167 insertion. **C**, Degree of helical twisting between base pairs. **D**, Degree of propeller twist within base pairs. **E**, Degree of buckle within base pairs. **F**, Disruption of the base pairing within the major groove below the recognition helix. For **A**, **C**, **D**, and **E**, the dotted line represents the norm for an idealized B-DNA molecule of the same sequence and length. The approximate positions of the recognition helices and Arg167 from each monomer are also shown.

Although the major groove width is close to ideal (**Figure 7A**), the bases are buckled and twisted, affecting the pattern of base pairing. The alignment of the base pairs (propeller twist) is especially altered where the recognition helices bind and where the Agr67 sidechain insert (**Figure 7D**). In addition, the base pairs are significantly buckled through the A—T track below the recognition helices (**Figure 7E**). This causes a deviation away from the Watson-Crick pattern in an idealized B-DNA molecule, such that the bases also form hydrogen bonds with opposing, but neighboring bases (**Figure 7F**).

In summary, the DNA binding domains of *Sp*NanR remain disengaged from the sugar isomerase domains when bound to the *nan*E recognition DNA sequence, explaining why *N*-acetylmannosamine-6-phosphate does not affect DNA binding. The *Sp*NanR recognition helix binds the major groove, as expected, but with only one long-range interaction with a base each side of the near palindromic sequence. Two arginine residues (Arg67 from each domain) insert into the minor groove, form an unusual π-stack conformation, each make an additional long-range hydrogen bond with a base, and together disrupt the minor groove width. These observations suggest that specificity is achieved through recognition of a sequence-dependent distorted B-DNA structure—the ‘indirect read-out’ mechanism—while affinity is primarily driven by interactions between the DNA binding domain and the phosphate backbone.

## Discussion

The import and catabolism of sialic acid is an adaptive mechanism for many bacterial pathogens [recent reviews (4,14,63,64)]. For *S. pneumoniae*, the link between sialic acid utilization and both colonization and pathogenesis is well-established. For example, addition of exogenous sialic acid leads to a significant increase in *nan* cluster transcription in naturally colonizing *Streptococci* (13), and it has been long known that sialidase enzymes (in this context the *sia*A and *nan*B genes), which cleave sialic acids from host glycoconjugates, are virulence factors for *S. pneumoniae* pathogenesis (14). Disrupting genes that import and metabolize sialic acid results in loss of fitness during colonization for many pathogenic bacteria (14,63,64), including *S. pneumoniae* (65). When in the presence of *N*-acetylmannosamine or sialic acid, *S. pneumoniae* upregulates expression of the machinery needed to catabolize sialic acid encoded by the *nan* gene cluster (23), and knocking out the *nan*R gene abolishes this response (22). Given the importance of sialic uptake for bacterial colonization and pathogenesis, including for *S. pneumoniae*, we set out to define the molecular mechanism by which *S. pneumoniae* senses sialic acid in the environment and controls the expression of the genes required for its metabolism. Here we propose a mechanism by which *Sp*NanR regulates sialic acid uptake that is consistent with our structural and biophysical data, and the previously reported functional data (22).

### Architecture and regulation across the nan gene cluster

The *nan* gene cluster comprises three operons (*nan* operon-I, *nan* operon-II and *sia*A) that generate three transcripts. Three *Sp*NanR recognition sites were previously reported across the cluster (22) and we identify two additional sites. Among the five identified sites, three are located within the promoter regions of the respective operons, while the remaining two are positioned at cryptic loci whose regulatory roles are less well defined (**Figure 1A**).

The *nan* operon-I and *sia*A operon transcripts are upregulated when *S. pneumoniae* is grown on sialic acid rich media (22). Their promoter regions are similar in architecture, with functional *Sp*NanR recognition sequences ∼55 base pairs upstream of the −35 box (**Supplementary Figure 17A-C**). This general structure is conserved across *S. pneumoniae* strains and other *Streptococcus* bacteria, especially for the *nan* operon-I (**Supplementary Table 10**). We demonstrate that *Sp*NanR binds the *nan* operon-I promoter region upstream of the RNA polymerase binding site, consistent with its role as an activator of transcription. Although we did not study the *Sp*NanR binding site within the *sia*A operon promoter site, it very likely functions identically to the *nan* operon-I promoter given their shared architecture.

Both promoter regions have a CcpA binding site (*cre* box) overlapping the −35 box (**Supplementary Figure 17B,C**), consistent with the observation that CcpA also controls these operons (**Figure 8A**) (22). CcpA knockout experiments demonstrate that both operons are repressed in the presence of glucose through binding of CcpA to the *cre* box (22). Glucose-induced repression of the genetic machinery specific for alternative carbon sources such as sialic acid is a theme in bacteria, first discovered in the well-characterized *E. coli lac* operon (66,67). This phenomenon exists in Gram-positive bacteria as well (68,69), and in *S. pneumoniae* CcpA controls global gene expression in response to glucose concentrations (30). Here, *S. pneumoniae* prioritizes glucose uptake and metabolism when present, consistent with the first growth phase of the diauxic pattern seen in **Figure 1C**.

**Figure 8.**
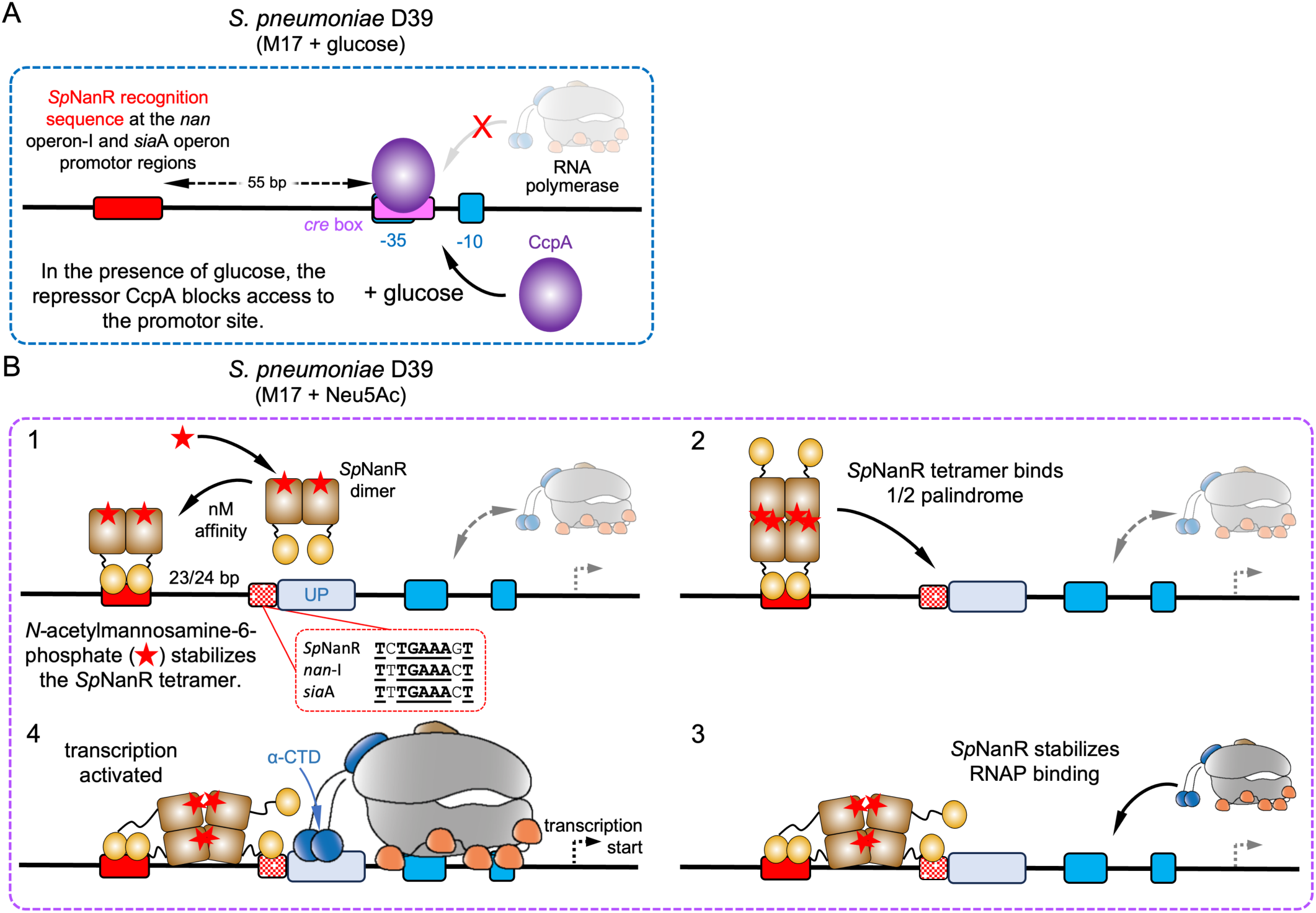
Proposed mechanisms for regulating the transcription of nan operons. **A**, In response to glucose availability, the global regulator CcpA (purple) binds to the *cre* site (pink) that overlaps the −35 box (blue) and represses transcription of *nan* operon-I by preventing RNA polymerase from binding. **B**, In the presence of Neu5Ac in media, the *nan* operon-I and *sia*A operon are upregulated. To explain this phenotype, we propose that *Sp*NanR binds *N*-acetylmannosamine-6-phosphate and stabilizes the tetramer, which then binds a degenerate half-site 23 base pairs downstream of the *Sp*NanR recognition sequence (steps 1 & 2). The half palindrome sequences found in the *nan* operon-I and *sia*A operon are compared to the *Sp*NanR recognition sequence (inset). The half palindrome is adjacent to the α-CTD binding site (UP element, light blue) and the *Sp*NanR DNA binding domain stabilizes the α-CTD, promoting transcription (steps 3 & 4).

In contrast, the *nan* operon-II transcript is not upregulated when *S. pneumoniae* is grown on sialic acid rich media (22). We identified a possible *Sp*NanR recognition sequence site within the *nan* operon-II promoter region (**Supplementary Figure 17D**), however it is the most divergent of the sequences (**Figure 1A**) and includes a T to G mutation at position 10 that we demonstrate abolishes *Sp*NanR binding to the *nan*E recognition sequence (**Figure 6D**, open red box) and therefore this site is likely to be nonfunctional. Also, this potential *Sp*NanR recognition sequence is downstream of the RNA polymerase −35 and −10 boxes, which suggests that if there is *Sp*NanR binding to this site it likely represses transcription. It is possible that *nan* operon-II, which encodes an ABC transporter, neuramidase, and an oxidoreductase that are predicted to metabolize unusual sialic acids or other amino sugars, is controlled by a transcriptional regulator that is yet to be unidentified.

Across the *nan* gene cluster there are two further *Sp*NanR recognition sequences with potentially cryptic functions and we demonstrate that *Sp*NanR can bind to both sites with nanomolar affinity. The functional *Sp*NanR recognition sequence upstream of the *nan*A gene is within *nan* operon-I (**Supplementary Figure 17E**), is not upregulated when *S. pneumoniae* is grown on sialic acid, and is considered nonfunctional (22). This makes sense because it lies within the *nan* operon-I transcript, which is controlled by the promoter upstream of the *nan*E gene. However, the region has identifiable −35 and −10 boxes, suggesting it may be functional as a promoter for the final two genes in the operon, perhaps with the help of additional unknown regulators. The *sia*A operon also has a *bona fide* functional *Sp*NanR recognition sequence located at the upstream of the operon with pseudo −35 and −10 boxes for transcription in the opposite direction (**Supplementary Figure 17F**), although there is no evidence that this protein is transcribed and the sequence contains early stop codons in all three frames. Overall, the functional significance of having *Sp*NanR recognition sequences located within *nan* operon-I upstream of the *nan*A gene and downstream of the *sia*A operon remains unclear.

### Trends and differences between SpNanR and other RpiR regulators

All RpiR regulators reported to date are repressors, whereas *Sp*NanR is an activator. Moreover, effector binding to RpiR regulators is reported to modulate DNA binding, whereas our biochemical and structural data demonstrate that effector binding to *Sp*NanR does not modulate DNA binding and instead stabilizes a higher order tetrameric state. To varying degrees, the molecular basis for repression has been studied in three cases: *V. vulnificus* NanR (*Vv*NanR), which also controls sialic acid metabolism (41); *E. coli* MurR (*Ec*MurR), which controls cell-wall recycling (36,40); and *P. aeruginosa* RccR (*Pa*RccR), which is a global regulator of carbon metabolism (42,43). Our structural and functional characterization of *Sp*NanR highlights conserved RpiR features as well as distinct differences.

Given the sequence similarity between these RpiR regulators, it is unsurprising that they share very similar DNA-binding and sugar isomerase domain structures. In all four cases (*Sp*NanR, *Vv*NanR, *Ec*MurR, and *Pa*RccR) the effector ligands are phosphorylated carbohydrates (*N*-acetylmannosamine-6-phosphate, *N*-acetylmuramate-6-phosphate, and 2-keto-3-deoxy-6-phosphogluconate) that bind to a site situated identically in their sugar isomerase domains.

Moreover, the phosphate binding loop (*i.e.*, loop-7 of *Sp*NanR) is highly conserved, comprising sidechain and mainchain contacts to the phosphate moiety. In contrast, the loops that bind the effector ligand distal from the phosphate are rearranged depending on the ligand the RpiR binds. Nonetheless, because of the location of the effector binding site at the tetrameric interface, all effector ligands bridge residues from three monomers across the tetramer. For *Sp*NanR, at least, this provides a mechanism by which the effector ligand stabilizes the tetrameric form.

All crystal structures of full-length RpiR regulators or independent sugar isomerase domains are tetrameric, including *Sp*NanR since the concentration during crystallization is very high (>>*K*_D_^4-2^). Some solution studies have been conducted, usually at high protein concentrations, to validate the tetrameric structure in their respective systems. For example, size exclusion chromatography and analytical ultracentrifugation experiments demonstrate that *Ec*MurR (36,40) and *Pa*RccR (42,70) are tetrameric, and this is the assumed biological unit. While a *Vv*NanR dimer is proposed to bind DNA, multiple oligomeric states are reported (dimer, tetramer, hexamer) when studied by size exclusion chromatography (41,71). How effector ligand binding affects oligomeric state has not been probed for these RpiR regulators—given the behavior of *Sp*NanR and the observation that the effector ligands bridge the tetramer, the oligomeric state of the RpiR family may, in general, be more important than first thought.

The reported RpiR structures also highlight another important difference. In both *Vv*NanR and *Pa*RccR, the DNA-binding domains are tightly coupled to the sugar isomerase domain (**Supplementary Figure 12**), enabling direct communication between the two domains. This structural arrangement provides a mechanism for allosteric control of DNA binding in response to effector ligand binding. In contrast, in-crystal and in-solution studies show the *Sp*NanR DNA binding domains are extended away and are not engaged with the isomerase domain, irrespective of effector ligand or DNA binding, explaining why the *N*-acetylmannosamine-6-phosphate effector when bound to the sugar isomerase domain has no effect on DNA binding.

### How does SpNanR sense sialic acid and upregulate transcription of nan operon-I?

Because the *nan* operon-I promoter region is responsive to sialic acid in the growth media and is best characterized by us (biophysically/structurally) and others (functionally) we consider how this locus is affected by the presence of a *Sp*NanR recognition sequence. Despite the *Sp*NanR recognition sequence being distal to the RNA polymerase −35 box of *nan* operon-I (55 base pairs) and the effector *N*-acetylmannosamine-6-phosphate not modulating DNA affinity, functional studies demonstrate that sialic acid significantly upregulates the *nan* operon-I transcript and that activation is abolished in a *nan*R knockout strain (22). We demonstrate that *N*-acetylmannosamine-6-phosphate does, however, stabilize a tetrameric form. We consider two possibilities.

An obvious effect of having multiple binding sites distributed across the *nan* gene cluster is that, upon forming a tetramer, *Sp*NanR can bridge these sites. This interaction effectively loops the 2D linear arrangement of the cluster to form a 3D structure. This behavior has been established in other bacterial transcriptional regulators. AraC, for instance, forms DNA loops in its tetrameric form, repressing transcription until the effector arabinose binds, which disrupts the tetramer and the subsequent dimers promote RNA polymerase binding and transcription (72–76). Unlike AraC, which dissociates into dimers upon ligand binding and activates transcription, the effector *N*-acetylmannosamine-6-phosphate stabilizes the tetrameric form of *Sp*NanR, thereby enhancing its potential to loop DNA without directly altering its binding affinity. An advantage of DNA looping is that it leads to higher concentration of *Sp*NanR within the vicinity of the *nan* gene cluster, although how this specifically activates transcription of *nan* operon-I is unclear. Another possibility is that the formation of the loop affects the local DNA structure, enhancing the affinity of σ factors or auxiliary proteins for the site and thereby promoting RNA polymerase function.

Although *Ec*MurR and *Vv*NanR are repressors, where effector binding decreases their affinity for DNA, their binding mode and promoter architecture is interesting. It is proposed that an *Ec*MurR tetramer bridges two adjacent DNA binding sites that overlap the polymerase binding sites (−35 and −10 boxes), explaining repression by *Ec*MurR. These sites are separated by 17 base pairs, and the tetramer would span a total 47 base pairs (36). Interestingly, *Vv*NanR also binds two adjacent sites (NANRBI and NANRBII) each ∼55 base pairs in length (71). In this case, *Vv*NanR dimers are proposed to bind to each site, although *Vv*NanR tetramers could plausibly bind (71). The minimal sequence that *Vv*NanR binds has not been defined, although the two regions span ∼107 base pairs.

Inspection of both the *nan* operon-I and the *sia*A operon promoter region do not identify adjacent *Sp*NanR binding sites, as found in the promoter regions that *Ec*MurR and *Vv*NanR bind, although there is a degenerate half palindrome ∼23 base pairs downstream of the *Sp*NanR sites in each operon (**Figure 8B**). Despite being degenerate and unlikely to bind *Sp*NanR in isolation, the affinity would be increased through avidity—that is, the proximity of the *Sp*NanR binding site increases the local concentration of *Sp*NanR DNA binding domains making even weak interactions more likely. Analysis of our *Sp*NanR structure suggests that the affinity of the DNA binding domains for DNA is determined largely through backbone interactions (**Figure 6**), suggesting even a degenerate half palindrome sequence could bind a *Sp*NanR DNA binding domain, where specificity is driven by the *bona fide* sequence downstream. Given that this site is immediately adjacent to the RNA polymerase α-CTD binding site (UP element) in *nan* operon-I and the *sia*A operon (**Supplementary Figure 17B-C**), one hypothesis is that *N*-acetylmannosamine-6-phosphate stabilizes the tetramer, which can bridge across and one of the DNA binding domains could bind the half palindrome (**Figure 8B**, steps 1 & 2). The distance *Sp*NanR would need to span across a linear B-DNA sequence is ∼150 Å, which is consistent with the distance of the extended tetramer in solution (**Supplementary Table 9**, **Supplementary Figure 13**, ∼160 Å). The flexibility of the *Sp*NanR linker region (residues 77–88), as demonstrated by our structural studies, would provide the domains significant flexibility to bind at both sites. The presence of a *Sp*NanR DNA binding domain at the half palindrome could enhance RNA polymerase binding through direct interactions with the α-CTD and stabilizes binding (**Figure 8B**, steps 3 & 4). In *E. coli*, binding of the RNA polymerase α-CTD to sequences upstream of the −35 box dramatically effects transcription rates (77,78). This hypothesis is consistent with our structural and biophysical data, and the previously reported activation of *nan* operon-I and the *sia*A operon in sialic acid containing media. A half-palindromic sequence is absent from the *nan*A promoter region (**Supplementary Figure 17E**), consistent with the observation that this region is not activated by SpNanR. It is also, to date, a unique mechanism for an RpiR regulator.

This model is also consistent with *Sp*NanR knockout studies, which result in significant decreases in expression of the *nan* operon-I transcript in M17 media with or without glucose or sialic acid supplementation [(22), see Figure 2A-B in their work], meaning that simply binding *Sp*NanR to the *nan* operon-I promoter region induces some expression. We demonstrate that *Sp*NanR binds to its recognition sequence in the *nan* operon-I promoter region with nanomolar affinity even in the absence of its effector *N*-acetylmannosamine-6-phosphate (**Figure 4**). Although the dissociation constant for *Sp*NanR in the absence of *N*-acetylmannosamine-6-phosphate is weak (*K*_D_^4-2^ = 97 μM), a tetramer can form and would be expected to reside intermittently at the *Sp*NanR recognition sequence. Again, the tetrameric *Sp*NanR could bridge across and one of the DNA binding domains binds the half palindrome. Here, avidity could play a role in stabilizing and promoting complex formation. From a biological perspective, this means that *Sp*NanR stimulates baseline transcription of the *nan* operon-I. The low-level expression of the sialic acid machinery, including a sialic acid ABC transporter, presumably allows *S. pneumoniae* to sense the presence of sialic acid in the environment.

## Conclusions

Our study reveals that *S. pneumoniae* NanR is a unique case within the RpiR family, functioning as a transcriptional activator rather than a repressor. Structural and biochemical analyses demonstrate that the effector, *N*-acetylmannosamine-6-phosphate, does not alter DNA binding affinity but instead stabilizes tetramer formation, establishing a ligand-dependent oligomerization mechanism distinct from previously characterized RpiR proteins. This decoupling of effector sensing from direct modulation of DNA binding expands the known repertoire of bacterial transcriptional regulation strategies. These findings provide a molecular framework for understanding sialic acid-responsive gene activation in pneumococci, and future work should aim to validate these mechanisms *in vivo*.

## Methods and Materials

DNA oligonucleotides were synthesized and annealed by GenScript. Upon receipt they were resuspended in SEC buffer (20 mM Tris-HCl (pH 8.0), 300 mM NaCl) at 100 mM concentrations and stored at −20 °C. DNA that was used in fluorescent experiments was labeled with 5’-FAM on the sense strand.

### Expression and purification of recombinant SpNanR

The protein sequence corresponding to *S. pneumoniae* NanR (gene: SPN23F_RS08445 NCBI Reference Sequence: WP_000360349.1) was synthesized by GenScript and cloned into pET30ΔSE. Mutant sequences were either ordered from GenScript directly or made in-house using mutagenesis PCR primers and ligation-independent cloning described previously (79).

The pET30ΔSE-*Sp*NanR and mutated constructs were transformed into *E. coli* BL21(DE3) competent cells (Agilent) and grown in LB media with 30 mg.ml^-1^ kanamycin at 37 °C and shaking at 220 rpm. Expression of *Sp*NanR was induced mid-log phase (OD600 ≈ 0.6) by the addition of IPTG to a final concentration of 1 mM and incubated overnight (14–16 h) at 25 °C with shaking at 220 rpm. Cells were harvested by centrifugation (Sorvall LYNX 4000 Superspeed) at 8,000 rpm for 10 min. Following centrifugation, the supernatant was discarded, and the cell pellet was flash cooled in liquid nitrogen for storage until needed.

To prepare the protein for purification, the cell pellet was resuspended in lysis buffer (20 mM Tris-HCl pH 8.0, 150 mM NaCl), supplemented with cOmplete™ mini protease cocktail inhibitor (Roche), and lysed by sonication (Hielscher UP200S Ultrasonic Processor) on ice. Cell debris and insoluble material was pelleted by centrifugation at 16,000 rpm for 30 min. The resulting cell lysate was then saturated to 50% ammonium sulphate and left to equilibrate for 1 h at 4 °C. Following precipitation, protein was pelleted at 10,000 rpm for 15 min and resuspended in buffer A (20 mM Tris-HCl pH 8.0, 100 mM NaCl), while the supernatant was discarded. This resuspended sample was dialyzed overnight in buffer A at 4 °C.

Purification of *S. pneumoniae* NanR was conducted using a three-step procedure on an ÄKTApure (Cytiva): anion exchange chromatography, heparin affinity chromatography and size-exclusion chromatography (**Supplementary Figure 1A**). The dialyzed sample was applied to a HisTrap Q FF column (GE Healthcare), pre-equilibrated in buffer A. Weakly associated or positively charged proteins were removed by washing the column for three column volumes of buffer A. Subsequently, the bound protein was eluted using a continuous gradient of buffer B (20 mM Tris-HCl pH 8.0, 1 M NaCl) over 60 min. Fractions containing protein were identified by SDS-PAGE and then pooled. The pooled sample was then applied to a Hi-Trap Heparin HP column (GE Healthcare), pre-equilibrated in buffer A. Next, the column was washed using three column volumes of buffer A, while the bound protein was eluted using a continuous gradient of buffer B over 20 min. Following concentration to >500 μL using ultrafiltration (Sartorius), the pooled sample was applied to a Superdex 200 Increase 10/300 GL column in buffer C (20 mM Tris-HCl pH 8.0, 300 mM NaCl) where *Sp*NanR eluted as a single peak. The absorbance at 280 nm of the purified protein was measured using a NanoDrop spectrophotometer and the concentration was estimated using a molar extinction coefficient of 27,390 M^-1^.cm^-1^ at 280 nm as calculated by ProtParam (80). All purification steps were carried out at 4 °C. Protein that was not immediately used in experiments was flash-frozen in liquid nitrogen and stored at −80 °C. We verified the mass of our protein using intact mass spectrometry, which shows that the protein has a mass of 32,611 Da (**Supplementary Figure 1B**).

### Differential scanning fluorimetry

A QuantStudio 3 real-time PCR system (ThermoFisher Scientific) was used to record the thermal stability of the purified protein with or without potential substrates. Reactions consisted of *Sp*NanR (0.65 mg.ml^-1^, 20 μM), 5× SYPRO® Orange (prepared as a 50× stock) and 1 mM effector in a final reaction volume of 25 μL. Once prepared these were loaded into a 96-well microplate (Applied Biosystems). Samples were heated from 4 °C to 95 °C at a ramp rate of 0.05 °C, taking fluorescent readings at each time point. Triplicate measurements were performed for each sample. To validate the experiment three negative controls (protein only, dye only, ligand only) were conducted. Data analysis was performed using Protein Thermal Shift software (version 1.3–Applied Biosystems) where an apparent melting point (T_m_) of each sample in °C was obtained from the highest point of the first derivative plot.

### Isothermal titration calorimetry

All experiments were performed in an ITC buffer (30 mM Tris, 300 mM NaCl,) while stirring at 250 rpm. The powdered stock of *N*-acetyl-D-mannosamine-6-phosphate was solubilized with the ITC buffer to a concentration of 2-5 mM and protein concentration of 200–500 μM before titrations. All titrations were performed using an initial injection of 1 μL followed by 125 identical injections of 2 μL with a duration of 4 seconds per injection and a spacing of 250 seconds between injections. The last four data points were averaged and subtracted from each titration to account for the heat of dilution (ligand into protein). Additional background experiments where buffer was titrated into the protein solution revealed no significant shift in the baseline during the measurements.

### Sedimentation velocity experiments

Sedimentation velocity experiments were performed with a Beckman Coulter XL-I with UV/visible detection optics, XL-A with a fluorescence detection system (Aviv Biomedical), or a Beckman Coulter Optima analytical ultracentrifuge with multi-wavelength UV/visible detection optics using either an An-60 Ti 4-hole or AN-50 Ti 8-hole rotor. Sample (380–460 μL) were loaded into a quartz cell with a 12 mm epon 2-channel centerpiece. Data were collected at 20 °C and at 42,000 rpm, acquired in intensity mode, using radial absorbance scans at a step size of 0.001 cm (Optima) or 0.003 cm (XL-I). Diluted samples were incubated at 4 °C overnight to ensure the self-associating protein molecules approached equilibrium and to minimize protein aggregation, followed by a further two hours at room temperature as the sedimentation experiment was setup and the rotor came to temperature (20 °C). SEDNTERP (81) and UltraScan (82) was used to calculate the v-bar of 0.7318 cm^3^/g for *Sp*NanR, buffer density of 1.0059 g/mL, and viscosity of 0.001034 Pa×s.

For the concentration series in the absence of *N*-acetylmannosamine-6-phosphate (**Figure 3, Supplementary Figure 4**), *Sp*NanR (4.3–227.7 μM) was incubated in size exclusion buffer C (20 mM Tris–HCl pH 8.0, 300 mM NaCl). For the concentration series with *N*-acetylmannosamine-6-phosphate (**Figure 3** and **Supplementary Figure 5**), *Sp*NanR (0.27–153.5 μM) was incubated in size exclusion buffer C plus 2 mM *N*-acetylmannosamine-6-phosphate. Data were collected using the Optima or XL-I instruments at wavelengths outlined in **Supplementary Table 3**.

For the experiments shown in **Figure 4A**, *Sp*NanR was serially diluted at a 2× concentration from 5 μM to 0.019 μM in similar fashion to above. The protein dilutions (500 μL) were mixed with 500 μL of 0.01 μM (2× final concentration) FAM-labeled *nan*E promoter sequence DNA (**Supplementary Table 4**) in SEC buffer. Following overnight incubation at 4 °C, the samples were either prepared for the analytical ultracentrifuge experiment as above (*Sp*NanR + DNA experiment), or (*Sp*NanR + DNA + *N*-acetylmannosamine-6-phosphate experiment) each 1 mL sample was spiked with 10 μL of 200 mM *N*-acetylmannosamine-6-phosphate and incubated for four extra hours at 4 °C followed by a further two hours at room temperature as the sedimentation experiment was setup and the rotor came to temperature (20 °C).

For the multiwavelength experiments in **Figure 4C** and **Supplementary Figure 10**, absorption spectra (210–310 nm) were collected using a Genesys S10 UV-Vis spectrophotometer (Thermo Fisher Scientific) for each component: *Sp*NanR protein, *nan*E promoter DNA sequence, and size exclusion buffer C with or without *N*-acetylmannosamine-6-phosphate. The spectra were globally fit using a nonlinear least square optimization to yield a global extinction profile for each analyte from 210–310 nm as described in (47), which was then used to deconvolute the combined sedimentation absorbance profiles acquired in the multiwavelength centrifuge experiment, as described in (83). All data were analyzed using UltraScan 4.0 (82). Initially, multi-wavelength sedimentation velocity datasets from each wavelength were analyzed using 2DSA (84) to remove systematic noise components, and to determine boundary conditions of the sample column as reported above. Iteratively refined 2DSA models from each wavelength were used to generate a sedimentation profile for each wavelength mapped to a common time grid Spectral deconvolution of the multiwavelength data using the molar extinction coefficient profiles of each spectral contributor generates the hydrodynamic results for each contributor. Wavelengths monitored in this experiment were 223, 225, 227, 229, 231, 233, 258, 260, 262, 276, 278, 280, 282 nm. The data were collected and processed as described previously (83,85).

All data were analyzed using UltraScan 4.0, release 7529 (82). Sedimentation data were evaluated by the two-dimensional spectrum analysis (2DSA) (44), which affords an unbiased hydrodynamic model, where the sedimentation coefficient and frictional ratio (*f/f*_0_) can be floated independently. The models were further subjected to iterative refinement (2DSA-IT). All fitting procedures were completed using the UltraScan LIMS cluster. Each 2DSA-IT model was visually assessed to ensure a good fit and a low root mean squared deviation using the FE Model Viewer utility in UltraScan (**Supplementary Figures 4B and 5B**). Peak integration of sample components was performed using the Genetic Algorithm Initialization Utility in UltraScan.

The weight-averaged partial specific volume was estimated for each complex using the following equation:

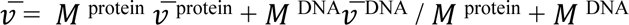

where *M* ^protein^ is the molar mass of the protein component (Da), *M* ^DNA^ is the molar mass of the DNA component (Da), *v̅* ^protein^ is the partial specific volume of the protein (mL/g), *v̅* ^DNA^ is the partial specific volume of the DNA, which is 0.55 mL.g^-1^.

### Microscale thermophoresis

Microscale thermophoresis (MST) was carried out using a Monolith NT.115 series instrument (NanoTemper) at 20 °C. FAM-labeled nucleotides identical to those used in the fluorescent AUC experiments were used as the target. Serial dilution of *Sp*NanR was performed in PCR 8-strip tubes with individual caps (USA Scientific) following the user manual provided by NanoTemper. 2× FAM-labeled DNA oligonucleotides (40 nM) were mixed with titrated protein overnight. Samples with a final dye concentration of 20 nM were then loaded into MO-K003 Monolith NT.115 standard capillaries (NanoTemper) and were measured at 60% LED and 40% MST power. SEC buffer (20 mM Tris-HCl pH 8.0, 300 mM NaCl) with 0.05% (v/v) Tween-20 was used for MST studies.

### Native mass spectrometry

Samples for native mass spectrometry were exchanged from the SEC buffer into 0.5 M ammonium acetate pH 7.4 *via* a single-pass through a Bio-Spin P-6 column (Bio-Rad). Final concentrations are as quoted in **Supplementary Figure 6**. Buffer-exchanged samples were then pipetted into gold-coated glass capillaries (1.0 mm outer diameter/0.75 mm inner diameter with filament), fabricated with orifice sizes of 1–5 µm, using a P-2000 puller (Sutter). Spectra were acquired using a Synapt XS (Waters) equipped with a 32k quad, with instrument settings optimized for the transmission of high-mass ions, and soft ionization and activation conditions as previously described (86). The instrument was mass calibrated using CsI. Spectra were smoothed (3 × 20, mean) and mass assigned using a combination of MassLynx (Waters), standard deviation analysis (87), and UniDec (88). Due to the salty nature of the peaks in the spectra, only those which could be unambiguously assigned have been labeled.

### Crystallography

Protein crystallization. **Supplementary Table 7** summarizes the temperature, crystallization buffer, concentration cryoprotectant and other parameters for the crystallization of *Sp*NanR with different ligands reported in this study. Briefly, 1 μL protein solution was mixed with 1 μL reservoir solution. The drops were incubated for 4–16 weeks before harvest. During the crystal harvest, each crystal was soaked in cryoprotectant (reservoir buffer + 20% (v/v) ethylene glycol/glycerol mix) for 2–5 seconds before snap-freezing in liquid nitrogen and data collection.

Data collection. Diffraction data were collected on the Australian Synchrotron MX2 beamline equipped with a Dectris EIGER 16M detector, and initial scaling and quality assessment performed using the XDS pipeline and AIMLESS (89,90). The first structure (sugar isomerase domain of *Sp*NanR + *N*-acetylmannosamine-6-phosphate, PDB ID: 8TX9) was phased experimentally (see below) and this was used as a template to solve subsequent structures by molecular replacement (PHASER (91), from the CCP4 suite of programs). The models were further refined using Refmac5 (92) and phenix.refine (93) of the CCP4 and PHENIX suites, respectively. The models were visually inspected by COOT before being deposited in the PDB (94). Structural figures were prepared using Pymol (v 3.1.4.1).

Experimental phasing. Protein crystals of *Sp*NanR + *N*-acetylmannosamine-6-phosphate were added to mother liquor that had been supplemented with triiodide (10 mM) and soaked from 1–5 minutes; the crystals cracked if soaked longer. The crystals were washed in their mother liquor and then a cryoprotectant mix of 20% ethylene glycol/glycerol, both supplemented with triiodide, and flash frozen. We used a single wavelength anomalous diffraction (SAD) strategy. Diffraction data were collected to a resolution of 2.6–3.0 Å and processed in space group *P*2_1_. To enhance the weak anomalous signal, seven data sets were merged (see **Supplementary Table 8**). Initial phases were estimated using the Auto-Rickshaw SAD phasing protocol (95,96), which incorporates SHELX (97). The resulting partial structure, comprising only the sugar isomerase domain, was used as a starting model for the MRSAD phasing protocol in Auto-Rickshaw (95,96), with the same anomalous data. This protocol identified 16 monomers (four tetramers) in the asymmetric unit and located 38 triiodide molecules, corresponding to two triiodide sites per monomer. A single tetramer of the sugar isomerase domains was subsequently used as a template for molecular replacement with the higher resolution *I*4 dataset (*Sp*NanR + *N*-acetylmannosamine-6-phosphate, PDB id: 8TX9), again using Auto-Rickshaw (95,96).

Structure determination and refinement. Phaser (91) from the CCP4 suite of programs was used to phase structures using the sugar isomerase domain of *Sp*NanR + *N*-acetylmannosamine-6-phosphate (PDB id: 8TX9) as the search model. The models were further refined using Refmac5 (92) and phenix.refine (93) of the CCP4 and PHENIX suites respectively. The models were visually inspected by COOT before being deposited in the PDB (94).

For the structure with *Sp*NanR + *N*-acetylmannosamine-6-phosphate, weak density is observed for the DNA-binding domains (∼30% of the structure) for the 2.01 Å resolution structure, although the crystal has ample space for these domains (**Supplementary Figure 11A**). As a result, the *R*_free_ is understandably high (27.1%) despite the high-resolution data. Further crystallization trials yielded crystals that diffracted to a lower 2.95 Å resolution, but only two of the four DNA-binding domains could now be modeled (*R*_free_ = 30.1%). There was residual density for the other two DNA binding domains, which could not be modeled.

Crystallization of the apo structure was significantly more difficult given the oligomeric heterogeneity (*K*_D_^4-2^ = 97 μM, 3.8 mg.mL^-1^), but crystals that yielded sufficient resolution data were eventually achieved (2.55 Å). Again, the *R*_free_ is rather high (31.5%), which we attribute to the structure missing one of the four DNA binding domains and the density for the other three being weak due to flexibility (average B-factor for the DNA domains Cα atoms is 85.1 Å^2^, whereas the average B-factor for the isomerase domains is 68.3 Å^2^). We determined the structure in the space group P1 and tested various orthorhombic and monoclinic space group possibilities. In all cases, the fourth DNA binding domain of the tetramer could not be resolved. Xtriage did not suggest twinning or pseudosymmetry.

The structure of *Sp*NanR in complex with DNA was refined using phenix.refine using NCS and Ramachandran restraints, given the low resolution (2.97 Å), and manually built using Coot. The final model has good geometry and just four Ramachandran outliers (0.4%); these residues reside in difficult to model loops. The N- and C-terminal residues were trimmed as there was little density to guide building. Density inconsistent with waters was modeled with Mg^2+^ and several ligands could be confidently placed—four glycerol molecules (from the cryoprotection solution) and a HEPES from the crystallization conditions.

## Data Availability

All reagents are available under material transfer agreement. All data relating to this work is available upon reasonable request. The atomic models for *Sp*NanR alone, bound to *N*-acetylmannosamine-6-phosphate and the *Sp*NanR/*nan*E DNA recognition sequence complex are available through the Protein Data Bank with the accession codes 8V5F, 8TX9, 8V4V and 9DQN, respectively.

## Funding

This work was supported by the following funding to R.C.J.D: 1) the New Zealand Royal Society Marsden Fund (contract UOC1506); 2) a Ministry of Business, Innovation and Employment Smart Ideas grant (contract UOCX1706); and 3) the Biomolecular Interaction Centre (University of Canterbury). C.R.H acknowledges: 1) the Canterbury Medical Research Foundation; 2) the Maurice Wilkins Centre; and 3) National Health and Medical Research Council of Australia for grant (2034044). B.D. acknowledges support from grants from the National Institutes of Health 1R01GM120600, the Canada Research Chairs program C150-2017-00015, and the Canadian Natural Science and Engineering Research Council Discovery Grant DG-RGPIN-2019-05637. The Canadian Center for Hydrodynamics is funded by the Canada Foundation for Innovation grant CFI-37589 (B.D.). UltraScan supercomputer calculations were supported through NSF/ACCESS grant TG-MCB070039N (B.D.). We thank Prof. Aron Fenton for insightful comments and ideas on cooperativity and allostery.

## Acknowledgements

We acknowledge the Australian Synchrotron, the staff at the MX2 beamline and Dr Nigel Kirby and Dr Tim Ryan for provision of synchrotron facilities and for their assistance in using the beamlines. Fluorescence-detection sedimentation velocity experiments at the Melbourne Protein Characterisation Platform, Bio21 Institute, University of Melbourne. MWL-SV experiments were performed at the Canadian Center for Hydrodynamics, University of Lethbridge. We thank Prof. Dr. Sven Hammerschmidt (Universität Greifswald) for advice.

## Notes

### Competing Interest Statement

The authors have declared no competing interest.

## References

1. Buckwalter, C.M. and King, S.J. (2012) Pneumococcal carbohydrate transport: food for thought. Trends Microbiol, 20, 517–522.

2. Pezzulo, A.A., Gutierrez, J., Duschner, K.S., McConnell, K.S., Taft, P.J., Ernst, S.E., Yahr, T.L., Rahmouni, K., Klesney-Tait, J., Stoltz, D.A. et al. (2011) Glucose depletion in the airway surface liquid is essential for sterility of the airways. PLoS One, 6, e16166.

3. Trappetti, C., Kadioglu, A., Carter, M., Hayre, J., Iannelli, F., Pozzi, G., Andrew, P.W. and Oggioni, M.R. (2009) Sialic acid: a preventable signal for pneumococcal biofilm formation, colonization, and invasion of the host. J Infect Dis, 199, 1497–1505.

4. McDonald, N.D., Lubin, J.B., Chowdhury, N. and Boyd, E.F. (2016) Host-Derived Sialic Acids Are an Important Nutrient Source Required for Optimal Bacterial Fitness In Vivo. mBio, 7, e02237–02215.

5. Sillanaukee, P., Ponnio, M. and Jaaskelainen, I.P. (1999) Occurrence of sialic acids in healthy humans and different disorders. Eur J Clin Invest, 29, 413–425.

6. Bulai, T., Bratosin, D., Pons, A., Montreuil, J. and Zanetta, J.P. (2003) Diversity of the human erythrocyte membrane sialic acids in relation with blood groups. FEBS Lett, 534, 185–189.

7. Blaum, B.S., Hannan, J.P., Herbert, A.P., Kavanagh, D., Uhrin, D. and Stehle, T. (2015) Structural basis for sialic acid-mediated self-recognition by complement factor H. Nat Chem Biol, 11, 77–82.

8. Parker, D., Soong, G., Planet, P., Brower, J., Ratner, A.J. and Prince, A. (2009) The NanA neuraminidase of Streptococcus pneumoniae is involved in biofilm formation. Infect Immun, 77, 3722–3730.

9. Wren, J.T., Blevins, L.K., Pang, B., Basu Roy, A., Oliver, M.B., Reimche, J.L., Wozniak, J.E., Alexander-Miller, M.A. and Swords, W.E. (2017) Pneumococcal Neuraminidase A (NanA) Promotes Biofilm Formation and Synergizes with Influenza A Virus in Nasal Colonization and Middle Ear Infection. Infect Immun, 85.

10. North, R.A., Horne, C.R., Davies, J.S., Remus, D.M., Muscroft-Taylor, A.C., Goyal, P., Wahlgren, W.Y., Ramaswamy, S., Friemann, R. and Dobson, R.C.J. (2018) “Just a spoonful of sugar…”: import of sialic acid across bacterial cell membranes. Biophys Rev, 10, 219–227.

11. Dhanabalan, K., Cheng, Y., Thach, T. and Subramanian, R. (2024) Many locks to one key: N-acetylneuraminic acid binding to proteins. IUCrJ, 11, 664–674.

12. Kim, J. and Kim, B.S. (2023) Bacterial Sialic Acid Catabolism at the Host-Microbe Interface. J Microbiol, 61, 369–377.

13. Pezzicoli, A., Ruggiero, P., Amerighi, F., Telford, J.L. and Soriani, M. (2012) Exogenous sialic acid transport contributes to group B streptococcus infection of mucosal surfaces. J Infect Dis, 206, 924–931.

14. Vimr, E.R. (2013) Unified theory of bacterial sialometabolism: how and why bacteria metabolize host sialic acids. ISRN Microbiol, 2013, 816713.

15. Almagro-Moreno, S. and Boyd, E.F. (2009) Sialic acid catabolism confers a competitive advantage to pathogenic vibrio cholerae in the mouse intestine. Infect Immun, 77, 3807–3816.

16. Jeong, H.G., Oh, M.H., Kim, B.S., Lee, M.Y., Han, H.J. and Choi, S.H. (2009) The Capability of Catabolic Utilization of N- Acetylneuraminic Acid, a Sialic Acid, Is Essential for Vibrio vulnificus Pathogenesis. Infection and Immunity, 77, 3209–3217.

17. Vimr, E., Lichtensteiger, C. and Steenbergen, S. (2000) Sialic acid metabolism’s dual function in Haemophilus influenzae. Mol Microbiol, 36, 1113–1123.

18. Troy, F.A. and McCloskey, M.A. (1979) Role of a membranous sialyltransferase complex in the synthesis of surface polymers containing polysialic acid in Escherichia coli. Temperature-induced alteration in the assembly process. J Biol Chem, 254, 7377–7387.

19. Olson, M.E., King, J.M., Yahr, T.L. and Horswill, A.R. (2013) Sialic acid catabolism in Staphylococcus aureus. J Bacteriol, 195, 1779–1788.

20. Severi, E., Randle, G., Kivlin, P., Whitfield, K., Young, R., Moxon, R., Kelly, D., Hood, D. and Thomas, G.H. (2005) Sialic acid transport in Haemophilus influenzae is essential for lipopolysaccharide sialylation and serum resistance and is dependent on a novel tripartite ATP-independent periplasmic transporter. Mol Microbiol, 58, 1173–1185.

21. Shell, D.M., Chiles, L., Judd, R.C., Seal, S. and Rest, R.F. (2002) The Neisseria lipooligosaccharide-specific alpha-2,3-sialyltransferase is a surface-exposed outer membrane protein. Infect Immun, 70, 3744–3751.

22. Afzal, M., Shafeeq, S., Ahmed, H. and Kuipers, O.P. (2015) Sialic acid-mediated gene expression in Streptococcus pneumoniae and role of NanR as a transcriptional activator of the nan gene cluster. Appl Environ Microbiol, 81, 3121–3131.

23. Gualdi, L., Hayre, J.K., Gerlini, A., Bidossi, A., Colomba, L., Trappetti, C., Pozzi, G., Docquier, J.D., Andrew, P., Ricci, S. et al. (2012) Regulation of neuraminidase expression in Streptococcus pneumoniae. BMC Microbiol, 12, 200.

24. Coombes, D., Davies, J.S., Newton-Vesty, M.C., Horne, C.R., Setty, T.G., Subramanian, R., Moir, J.W.B., Friemann, R., Panjikar, S., Griffin, M.D.W. et al. (2020) The basis for non-canonical ROK family function in the N-acetylmannosamine kinase from the pathogen Staphylococcus aureus. J Biol Chem, 295, 3301–3315.

25. Horne, C.R., Kind, L., Davies, J.S. and Dobson, R.C.J. (2020) On the structure and function of Escherichia coli YjhC: An oxidoreductase involved in bacterial sialic acid metabolism. Proteins, 88, 654–668.

26. Currie, M.J., Manjunath, L., Horne, C.R., Rendle, P.M., Subramanian, R., Friemann, R., Fairbanks, A.J., Muscroft-Taylor, A.C., North, R.A. and Dobson, R.C.J. (2021) N-acetylmannosamine-6-phosphate 2-epimerase uses a novel substrate-assisted mechanism to catalyze amino sugar epimerization. J Biol Chem, 297, 101113.

27. North, R.A., Watson, A.J., Pearce, F.G., Muscroft-Taylor, A.C., Friemann, R., Fairbanks, A.J. and Dobson, R.C. (2016) Structure and inhibition of N-acetylneuraminate lyase from methicillin-resistant Staphylococcus aureus. FEBS Lett, 590, 4414–4428.

28. Gangi Setty, T., Sarkar, A., Coombes, D., Dobson, R.C.J. and Subramanian, R. (2020) Structure and Function of N-Acetylmannosamine Kinases from Pathogenic Bacteria. ACS Omega, 5, 30923–30936.

29. Paysan-Lafosse, T., Blum, M., Chuguransky, S., Grego, T., Pinto, B.L., Salazar, G.A., Bileschi, M.L., Bork, P., Bridge, A., Colwell, L. et al. (2023) InterPro in 2022. Nucleic Acids Res, 51, D418–D427.

30. Carvalho, S.M., Kloosterman, T.G., Kuipers, O.P. and Neves, A.R. (2011) CcpA ensures optimal metabolic fitness of Streptococcus pneumoniae. PLoS One, 6, e26707.

31. Monod, J. and Bordet, J. (1942).

32. Monod, J.L. (1978).

33. Aravind, L., Anantharaman, V., Balaji, S., Babu, M. and Iyer, L. (2005) The many faces of the helix-turn-helix domain: Transcription regulation and beyond. FEMS Microbiology Reviews, 29, 231–262.

34. Bateman, A. (1999) The SIS domain: a phosphosugar-binding domain. Trends Biochem Sci, 24, 94–95.

35. Kohler, P.R., Choong, E.L. and Rossbach, S. (2011) The RpiR-like repressor IolR regulates inositol catabolism in Sinorhizobium meliloti. J Bacteriol, 193, 5155–5163.

36. Jaeger, T. and Mayer, C. (2008) The transcriptional factors MurR and catabolite activator protein regulate N-acetylmuramic acid catabolism in Escherichia coli. J Bacteriol, 190, 6598–6608.

37. Shimada, T., Ogasawara, H. and Ishihama, A. (2018) Single-target regulators form a minor group of transcription factors in Escherichia coli K-12. Nucleic Acids Res, 46, 3921–3936.

38. Flores-Bautista, E., Cronick, C.L., Fersaca, A.R., Martinez-Nunez, M.A. and Perez-Rueda, E. (2018) Functional Prediction of Hypothetical Transcription Factors of Escherichia coli K-12 Based on Expression Data. Comput Struct Biotechnol J, 16, 157–166.

39. Sorensen, K.I. and Hove-Jensen, B. (1996) Ribose catabolism of Escherichia coli: characterization of the rpiB gene encoding ribose phosphate isomerase B and of the rpiR gene, which is involved in regulation of rpiB expression. J Bacteriol, 178, 1003–1011.

40. Zhang, Y., Chen, W., Wu, D., Liu, Y., Wu, Z., Li, J., Zhang, S.Y. and Ji, Q. (2022) Molecular basis for cell-wall recycling regulation by transcriptional repressor MurR in Escherichia coli. Nucleic Acids Res, 50, 5948–5960.

41. Hwang, J., Kim, B.S., Jang, S.Y., Lim, J.G., You, D.J., Jung, H.S., Oh, T.K., Lee, J.O., Choi, S.H. and Kim, M.H. (2013) Structural insights into the regulation of sialic acid catabolism by the Vibrio vulnificus transcriptional repressor NanR. Proc Natl Acad Sci U S A, 110, E2829–2837.

42. Zhu, Y., Mou, X., Song, Y., Zhang, Q., Sun, B., Liu, H., Tang, H. and Bao, R. (2024) Molecular mechanism of the one-component regulator RccR on bacterial metabolism and virulence. Nucleic Acids Res, 52, 3433–3449.

43. Wang, Y., Wang, Z., Chen, W., Ren, Z.H., Gao, H., Dai, J., Luo, G.Z., Wu, Z. and Ji, Q. (2024) A KDPG sensor RccR governs Pseudomonas aeruginosa carbon metabolism and aminoglycoside antibiotic tolerance. Nucleic Acids Res, 52, 967–976.

44. Brookes, E., Cao, W. and Demeler, B. (2010) A two-dimensional spectrum analysis for sedimentation velocity experiments of mixtures with heterogeneity in molecular weight and shape. Eur Biophys J, 39, 405–414.

45. Demeler, B. and Brookes, E. (2008) Monte Carlo analysis of sedimentation experiments. Colloid and Polymer Science, 286, 129–137.

46. Demeler, B. and van Holde, K.E. (2004) Sedimentation velocity analysis of highly heterogeneous systems. Anal Biochem, 335, 279–288.

47. Demeler, B. (2024) Methods for the Design and Analysis of Analytical Ultracentrifugation Experiments. Curr Protoc, 4, e974.

48. Demeler, B. (2010) Methods for the design and analysis of sedimentation velocity and sedimentation equilibrium experiments with proteins. *Curr Protoc Protein Sci*, **Chapter** 7, Unit 7 13.

49. Zhao, H., Piszczek, G. and Schuck, P. (2015) SEDPHAT--a platform for global ITC analysis and global multi-method analysis of molecular interactions. Methods, 76, 137–148.

50. Zhao, H., Li, W., Chu, W., Bollard, M., Adao, R. and Schuck, P. (2020) Quantitative Analysis of Protein Self-Association by Sedimentation Velocity. Curr Protoc Protein Sci, 101, e109.

51. Brookes, E.H. and Demeler, B. (2007), *Proceedings of the 9th annual conference on Genetic and evolutionary computation*. Association for Computing Machinery, London, England, pp. 361–368.

52. Harrison, S.C. (1991) A structural taxonomy of DNA-binding domains. Nature, 353, 715–719.

53. Teplyakov, A., Obmolova, G., Badet-Denisot, M.A. and Badet, B. (1999) The mechanism of sugar phosphate isomerization by glucosamine 6-phosphate synthase. Protein Sci, 8, 596–602.

54. Natalello, A., Santambrogio, C. and Grandori, R. (2017) Are Charge-State Distributions a Reliable Tool Describing Molecular Ensembles of Intrinsically Disordered Proteins by Native MS? J Am Soc Mass Spectrom, 28, 21–28.

55. Testa, L., Brocca, S., Santambrogio, C., D’Urzo, A., Habchi, J., Longhi, S., Uversky, V.N. and Grandori, R. (2013) Extracting structural information from charge-state distributions of intrinsically disordered proteins by non-denaturing electrospray-ionization mass spectrometry. Intrinsically Disord Proteins, 1, e25068.

56. Garvie, C.W. and Wolberger, C. (2001) Recognition of specific DNA sequences. Mol Cell, 8, 937–946.

57. Rohs, R., West, S.M., Sosinsky, A., Liu, P., Mann, R.S. and Honig, B. (2009) The role of DNA shape in protein-DNA recognition. Nature, 461, 1248–1253.

58. Otwinowski, Z., Schevitz, R.W., Zhang, R.G., Lawson, C.L., Joachimiak, A., Marmorstein, R.Q., Luisi, B.F. and Sigler, P.B. (1988) Crystal structure of trp repressor/operator complex at atomic resolution. Nature, 335, 321–329.

59. Ngo, T.T., Zhang, Q., Zhou, R., Yodh, J.G. and Ha, T. (2015) Asymmetric unwrapping of nucleosomes under tension directed by DNA local flexibility. Cell, 160, 1135–1144.

60. Olson, W.K., Gorin, A.A., Lu, X.J., Hock, L.M. and Zhurkin, V.B. (1998) DNA sequence-dependent deformability deduced from protein-DNA crystal complexes. Proc Natl Acad Sci U S A, 95, 11163–11168.

61. Chua, E.Y., Vasudevan, D., Davey, G.E., Wu, B. and Davey, C.A. (2012) The mechanics behind DNA sequence-dependent properties of the nucleosome. Nucleic Acids Res, 40, 6338–6352.

62. Marin-Gonzalez, A., Vilhena, J.G., Perez, R. and Moreno-Herrero, F. (2021) A molecular view of DNA flexibility. Q Rev Biophys, 54, e8.

63. Jennings, M.P., Day, C.J. and Atack, J.M. (2022) How bacteria utilize sialic acid during interactions with the host: snip, snatch, dispatch, match and attach. Microbiology (Reading), 168.

64. Severi, E., Rudden, M., Bell, A., Palmer, T., Juge, N. and Thomas, G.H. (2021) Multiple evolutionary origins reflect the importance of sialic acid transporters in the colonization potential of bacterial pathogens and commensals. Microb Genom, 7.

65. Marion, C., Burnaugh, A.M., Woodiga, S.A. and King, S.J. (2011) Sialic acid transport contributes to pneumococcal colonization. Infect Immun, 79, 1262–1269.

66. Loomis, W.F., Jr. and Magasanik, B. (1967) Glucose-lactose diauxie in Escherichia coli. J Bacteriol, 93, 1397–1401.

67. Magasanik, B. and Neidhardt, F.C. (1956) Inhibitory effect of glucose on enzyme formation. Nature, 178, 801–802.

68. Miwa, Y., Nagura, K., Eguchi, S., Fukuda, H., Deutscher, J. and Fujita, Y. (1997) Catabolite repression of the Bacillus subtilis gnt operon exerted by two catabolite-responsive elements. Mol Microbiol, 23, 1203–1213.

69. Gosseringer, R., Kuster, E., Galinier, A., Deutscher, J. and Hillen, W. (1997) Cooperative and non-cooperative DNA binding modes of catabolite control protein CcpA from Bacillus megaterium result from sensing two different signals. J Mol Biol, 266, 665–676.

70. Hwang, W., Yong, J.H., Min, K.B., Lee, K.M., Pascoe, B., Sheppard, S.K. and Yoon, S.S. (2021) Genome-wide association study of signature genetic alterations among pseudomonas aeruginosa cystic fibrosis isolates. PLoS Pathog, 17, e1009681.

71. Kim, B.S., Hwang, J., Kim, M.H. and Choi, S.H. (2011) Cooperative regulation of the Vibrio vulnificus nan gene cluster by NanR protein, cAMP receptor protein, and N-acetylmannosamine 6-phosphate. J Biol Chem, 286, 40889–40899.

72. Martin, K., Huo, L. and Schleif, R.F. (1986) The DNA loop model for ara repression: AraC protein occupies the proposed loop sites in vivo and repression-negative mutations lie in these same sites. Proc Natl Acad Sci U S A, 83, 3654–3658.

73. Saviola, B., Seabold, R. and Schleif, R.F. (1998) Arm-domain interactions in AraC. J Mol Biol, 278, 539–548.

74. Seabold, R.R. and Schleif, R.F. (1998) Apo-AraC actively seeks to loop. J Mol Biol, 278, 529–538.

75. Reed, W.L. and Schleif, R.F. (1999) Hemiplegic mutations in AraC protein. J Mol Biol, 294, 417–425.

76. Wu, M. and Schleif, R. (2001) Mapping arm-DNA-binding domain interactions in AraC. J Mol Biol, 307, 1001–1009.

77. Estrem, S.T., Gaal, T., Ross, W. and Gourse, R.L. (1998) Identification of an UP element consensus sequence for bacterial promoters. Proc Natl Acad Sci U S A, 95, 9761–9766.

78. Ross, W. and Gourse, R.L. (2005) Sequence-independent upstream DNA-alphaCTD interactions strongly stimulate Escherichia coli RNA polymerase-lacUV5 promoter association. Proc Natl Acad Sci U S A, 102, 291–296.

79. Park, J., Throop, A.L. and LaBaer, J. (2015) Site-specific recombinational cloning using gateway and in-fusion cloning schemes. Curr Protoc Mol Biol, 110, 3 20 21–23 20 23.

80. Walker, J.M. (2005) The proteomics protocols handbook. Humana Press, Totowa, N.J.

81. Philo, J.S. (2023) SEDNTERP: a calculation and database utility to aid interpretation of analytical ultracentrifugation and light scattering data. Eur Biophys J, 52, 233–266.

82. Demeler, B. and Gorbet, G.E. (2016) In Uchiyama, S., Arisaka, F., Stafford, W. F. and Laue, T. (eds.), Analytical Ultracentrifugation: Instrumentation, Software, and Applications. Springer Japan, Tokyo, pp. 119–143.

83. Horne, C.R., Henrickson, A., Demeler, B. and Dobson, R.C.J. (2020) Multi-wavelength analytical ultracentrifugation as a tool to characterise protein-DNA interactions in solution. Eur Biophys J, 49, 819–827.

84. Demeler, B., Aziz, Z., Behlke, J., Bernardi, G., Bourdillon, L., Butler, P.J.G., Carels, N., Clay, O., Colfen, H., Correia, J.J., et al. (2005) In Scott, D. J., Harding, S. E., Rowe, A. J., Scott, D., Harding, S. E. and Rowe, A. (eds.), Analytical Ultracentrifugation. The Royal Society of Chemistry, pp. 0.

85. Henrickson, A., Gorbet, G.E., Savelyev, A., Kim, M., Hargreaves, J., Schultz, S.K., Kothe, U. and Demeler, B. (2022) Multi-wavelength analytical ultracentrifugation of biopolymer mixtures and interactions. Anal Biochem, 652, 114728.

86. Ruotolo, B.T., Benesch, J.L., Sandercock, A.M., Hyung, S.J. and Robinson, C.V. (2008) Ion mobility-mass spectrometry analysis of large protein complexes. Nat Protoc, 3, 1139–1152.

87. McKay, A.R., Ruotolo, B.T., Ilag, L.L. and Robinson, C.V. (2006) Mass measurements of increased accuracy resolve heterogeneous populations of intact ribosomes. J Am Chem Soc, 128, 11433–11442.

88. Marty, M.T., Baldwin, A.J., Marklund, E.G., Hochberg, G.K., Benesch, J.L. and Robinson, C.V. (2015) Bayesian deconvolution of mass and ion mobility spectra: from binary interactions to polydisperse ensembles. Anal Chem, 87, 4370–4376.

89. Evans, P. (2006) Scaling and assessment of data quality. Acta Crystallogr D Biol Crystallogr, 62, 72–82.

90. Kabsch, W. (2010) Integration, scaling, space-group assignment and post-refinement. Acta Crystallogr D Biol Crystallogr, 66, 133–144.

91. McCoy, A.J., Grosse-Kunstleve, R.W., Adams, P.D., Winn, M.D., Storoni, L.C. and Read, R.J. (2007) Phaser crystallographic software. J Appl Crystallogr, 40, 658–674.

92. Murshudov, G.N., Vagin, A.A. and Dodson, E.J. (1997) Refinement of macromolecular structures by the maximum-likelihood method. Acta Crystallogr D Biol Crystallogr, 53, 240–255.

93. Liebschner, D., Afonine, P.V., Baker, M.L., Bunkoczi, G., Chen, V.B., Croll, T.I., Hintze, B., Hung, L.W., Jain, S., McCoy, A.J. et al. (2019) Macromolecular structure determination using X-rays, neutrons and electrons: recent developments in Phenix. Acta Crystallogr D Struct Biol, 75, 861–877.

94. Emsley, P., Lohkamp, B., Scott, W.G. and Cowtan, K. (2010) Features and development of Coot. Acta Crystallogr D Biol Crystallogr, 66, 486–501.

95. Panjikar, S., Parthasarathy, V., Lamzin, V.S., Weiss, M.S. and Tucker, P.A. (2005) Auto-rickshaw: an automated crystal structure determination platform as an efficient tool for the validation of an X-ray diffraction experiment. Acta Crystallogr D Biol Crystallogr, 61, 449–457.

96. Panjikar, S., Parthasarathy, V., Lamzin, V.S., Weiss, M.S. and Tucker, P.A. (2009) On the combination of molecular replacement and single-wavelength anomalous diffraction phasing for automated structure determination. Acta Crystallogr D Biol Crystallogr, 65, 1089–1097.

97. Uson, I. and Sheldrick, G.M. (2018) An introduction to experimental phasing of macromolecules illustrated by SHELX; new autotracing features. Acta Crystallogr D Struct Biol, 74, 106–116

