## Supplementary data for "Mechanism of NanR transcriptional activation of sialic acid metabolism in *Streptococcus pneumoniae*"

[a] Biomolecular Interaction Centre and School of Biological Sciences, University of Canterbury, PO Box 4800, Christchurch 8140, New Zealand.
[b] Mātai Hāora - Centre for Redox Biology and Medicine, Department of Pathology and Molecular Medicine, University of Otago, Christchurch 8041, New Zealand.
[c] Australian Synchrotron, ANSTO, Clayton, 800 Blackburn Road, Victoria 3168, Australia.
[d] Department of Biochemistry and Molecular Biology, Monash University, Melbourne, 3800, VIC, Australia.
[e] Department of Biochemistry and Pharmacology, Bio21 Molecular Science and Biotechnology Institute, University of Melbourne, Parkville, Victoria 3010, Australia.
[f] Biomolecular Interaction Centre, School of Physical and Chemical Sciences, University, of Canterbury, Christchurch, New Zealand.
[g] School of Medical Sciences, Faculty of Medicine and Health, University of Sydney, New South Wales 2006, Australia.
[h] Department of Chemistry and Biochemistry, University of Lethbridge, 4401 University Drive, Lethbridge, AB T1K 3M4, Canada.
[i] Department of Chemistry and Biochemistry, University of Montana, Missoula, MT 59812, USA.
[j] Walter and Eliza Hall Institute of Medical Research, 1G Royal Parade, Parkville, VIC 3050, Australia.
[k] Drug Discovery Biology, Monash Institute of Pharmaceutical Sciences, Monash University, Victoria 3052, Australia.

****Corresponding authors:*** *Prof. Renwick Dobson*, Biomolecular Interaction Centre and School of Biological Sciences, University of Canterbury, Christchurch 8140, New Zealand.; *Dr Christopher Horne*, Walter and Eliza Hall Institute of Medical Research, 1G Royal Parade, Parkville, VIC 3050, Australia.

**Supplementary Tables**

**Supplementary Table 1 | Protein sequence of *Sp*NanR expressed and purified in this study.**

| Protein | Protein sequence |
|---|---|
| *Sp*NanR (www.uniprot.org/uniprotkb/A0A0H2ZPE9/) | MDKPDIATVIDSHFEEMTDLEQEIARYFLQAETIQDDLSSQQVT QKLHISQAALTRFAKKCGFTGYREFIFQYQHEAENQANQVSKH SPLTKRVLRSYSNMREQTQDLIDEVQLERIAQLIEDAERVYFFG TGSSGLVAREMKLRFMRLGVVCEALTDQDGFAWTTSIMDENC LVLGFSLSGSTPSILDSLLDAKEMGAKTVLFSSVPNKDSQAYTE TVLVATHSQPSYIQRISAQLPMLFFIDLIYAYFLEINRESKEKIFN SYWENKKLNGYRRQKRVRKS |

**Supplementary Table 2 | *N*-Acetylmannosamine-6-phosphate (manNAc-6-P) stabilizes *Sp*NanR.** These differential scanning fluorimetry data are shown in **Figure 2A** and **Supplementary Figure 2**. Ligand concentrations were 1 mM and the *Sp*NanR concentration was 20 μM.

| ligand | Tm (°C) |
|---|---|
| *Sp*NanR only | 46.4 ± 0.1 |
| + Neu5Ac | 45.8 ± 0.5 |
| + manNAc | 46.2 ± 0.2 |
| + manNAc-6-P | **50.5 ± 0.1** |
| + glcNAc | 46.4 ± 0.1 |
| + glcNAc-6-P | 46.0 ± 0.1 |

**Supplementary Table 3 | Sedimentation velocity data and fits to the 2DSA model for *Sp*NanR with increasing concentrations as plotted in Supplementary Figure 4A and 5A.** UltraScan software was used to calculate the v-bar ($cm^3/g$ at 20 °C) of 0.7318, buffer density of 1.0059 (g/mL at 20 °C), and viscosity of 0.001034 (Pa·s). The weight-averaged S value was determined over an S range of 3–7 S.

| concentration (µM) | wavelength (nm) | weight average S | $f/f_o$ | variance | r.m.s.d. |
|---|---|---|---|---|---|
| ***Sp*NanR in size-exclusion buffer (Supplementary Figure 4)** | | | | | |
| 4.3 | 220 | 4.35 | 1.32 | 0.00017 | 0.013 |
| 9.2 | 278 | 4.48 | 1.88 | 0.000010 | 0.0031 |
| 15.4 | 278 | 4.70 | 1.27 | 0.000010 | 0.0031 |
| 21.5 | 281 | 4.73 | 1.15 | 0.000017 | 0.0041 |
| 36.9 | 281 | 5.04 | 1.22 | 0.000019 | 0.0044 |
| 52.3 | 291 | 5.23 | 1.30 | 0.000015 | 0.0039 |
| 101.5 | 291 | 5.32 | 1.36 | 0.000040 | 0.0063 |
| 138.5 | 296 | 5.69 | 1.13 | 0.000001 | 0.0039 |
| 227.7 | 298 | 5.68 | 1.18 | 0.000011 | 0.0032 |
| ***Sp*NanR + *N*-acetylmannosamine-6-phosphate (2 mM) (Supplementary Figure 5)** | | | | | |
| 0.27* | 220 | 6.42 | 1.42 | 0.000010 | 0.0022 |
| 0.54* | 220 | 6.39 | 1.46 | 0.0000034 | 0.0023 |
| 1.1* | 228 | 6.39 | 1.43 | 0.000062 | 0.0016 |
| 2.2* | 228 | 6.38 | 1.44 | 0.0000053 | 0.0025 |
| 4.4 | 232 | 6.36 | 1.48 | 0.0000061 | 0.0023 |
| 9.1 | 232 | 6.34 | 1.49 | 0.0000025 | 0.0079 |
| 17.9 | 280 | 6.36 | 1.47 | 0.0000052 | 0.0019 |
| 36.1 | 280 | 6.36 | 1.42 | 0.0000051 | 0.0032 |
| 152.5 | 289 | 6.34 | 1.56 | 0.00076 | 0.028 |

***** At the very low concentrations the data were collected at low wavelengths (220 and 228 nm) where the *N*-acetylmannosamine-6-phosphate containing buffer showed significant absorbance and gave a signal at ~0.2 S.

**Supplementary Table 4 | DNA sequence of *nan*E recognition region used for binding and crystallization studies.**

| oligonucleotide | DNA sequence |
|---|---|
| *nan*E recognition sequence | 5'TCTGAAAGTACTTTTAGA3' |
| Fluorescently labeled *nan*E recognition sequence | 5'6-FAM-TCTGAAAGTACTTTTAGA3' |

**Supplementary Table 5 | Fluorescent sedimentation velocity data and fits to the 2DSA model for *Sp*NanR with FAM-labeled *nan*E recognition DNA sequence (Supplementary Figure 8 & 9) using UltraScan.** UltraScan software was used to calculate the v-bar ($cm^3/g$ at 20 °C) of 0.7318 for *Sp*NanR protein (0.015–5 µM) and 0.55 was used for the 18 base pairs *nan*E DNA (0.05 µM), buffer density of 1.0059 (g/mL at 20 °C), and viscosity of 0.001034 (Pa·s). The weight-averaged S value was determined over an S range of 1.5–7 S.

| concentration (µM) | wavelength (nm) | weight average S | $f/f_o$ | variance | r.m.s.d. |
|---|---|---|---|---|---|
| ***Sp*NanR + 50 nM *nan*E DNA in size-exclusion buffer (Figure 4A, left)** | | | | | |
| 5 | 488 | 6.43 | 1.09 | 275 | 16.6 |
| 2.5 | 488 | 6.21 | 1 | 356 | 18.8 |
| 1.25 | 488 | 6.20 | 1 | 214 | 14.6 |
| 0.625 | 488 | 4.89 | 1.42 | 73 | 8.6 |
| 0.3125 | 488 | 3.71 | 1.39 | 76 | 10.1 |
| 0.15 | 488 | 2.95 | 1.25 | 96. | 9.8 |
| 0.078 | 488 | 2.47 | 1.17 | 116 | 10.8 |
| 0.039 | 488 | 2.22 | 1.24 | 94 | 9.7 |
| 0.015 | 488 | 2.35 | 1.14 | 94 | 9.1 |
| ***Sp*NanR 50 nM *nan*E DNA in size-exclusion buffer + *N*-acetylmannosamine-6-phosphate (2 mM) (Figure 4A, right)** | | | | | |
| 5 | 488 | 5.74 | 1.30661 | 56.1768 | 7.50 |
| 2.5 | 488 | 5.50 | 1.28346 | 52.0462 | 7.2 |
| 1.25 | 488 | 5.36 | 1.30361 | 63.3914 | 8.0 |
| 0.625 | 488 | 4.45 | 1.40433 | 64.8972 | 8.1 |
| 0.3125 | 488 | 3.32 | 1.50004 | 92.6381 | 9.6 |
| 0.15 | 488 | 2.75 | 1.35738 | 109.375 | 10.4 |
| 0.078 | 488 | 2.50 | 1.33834 | 147.239 | 12.1 |
| 0.039 | 488 | 2.31 | 1.18934 | 113.531 | 10.7 |
| 0.015 | 488 | 2.39 | 1.19986 | 106.149 | 10.3 |

**Supplementary Table 6 | Hydrogen bond interactions between *Sp*NanR and *N*-acetylmannosamine-6-phosphate (PDB ID: 8TX9).** Averaged over the four binding sites in the *Sp*NanR tetramer.

| **Hydrogen bonding distances between *Sp*NanR and manNAc-6-P** | | |
|---|---|---|
| **donor (D)** | **acceptor (A)** | **D-A distance (Å)** |
| Ser 134 N | BMX O4 | 2.72 |
| Ser 135 OG | BMX O17 | 2.72 |
| Arg148 NH1 | BMX O7 | 3.07 |
| Arg 148 NH2 | BMX O3 | 3.13 |
| Asp 160 OD2 | BMX O1 | 2.55 |
| Ser 179 OG | BMX O19 | 2.83 |
| Leu 180 N | BMX O17 | 2.85 |
| Ser 181 N | BMX O18 | 2.81 |
| Ser 181 OG | BMX O19 | 2.72 |
| Thr 184 OG1 | BMX O19 | 2.68 |
| Tyr 229 OH | BMX O3 | 2.81 |

**Supplementary Table 7 | Crystallization conditions.** The concentrations represent the stock solutions that were mixed 1:1 with sample to produce the final drop. *N*-acetylmannosamine-6-phosphate abbreviated to manNAc-6-P.

| **crystallization conditions** | ***Sp*NanR + manNAc-6-P + triiodide** | ***Sp*NanR + manNAc-6-P, condition 1** | ***Sp*NanR + manNAc-6-P, condition 2** | ***Sp*NanR** | ***Sp*NanR + DNA** |
|---|---|---|---|---|---|
| buffer | 100 mM MES (pH 6.5), 12% PEG 20K | 100 mM MES (pH 6.5), 12% PEG 20K | 0.1 M phosphate/citrate (pH 4.2), 40% PEG 3350 | 0.1 M Bis-Tris (pH 5.5), 0.2 M lithium sulfate, 25% w/v PEG 3350 | 0.1 M sodium HEPES (pH 7.5), 0.2 M magnesium chloride hexahydrate. 30% v/v PEG 400 |
| buffer volume (μL) | 1 | 1 | 1 | 1 | 1 |
| sample concentrations | *Sp*NanR (15.5 mg.mL$^{-1}$, ~475 μM) + manNAc-6-P (10 mM) + triiodide (1- mM) | *Sp*NanR (15.5 mg.mL$^{-1}$, ~475 μM) + manNAc-6-P (10 mM) | *Sp*NanR (15.5 mg.mL$^{-1}$, ~475 μM) + manNAc-6-P (10 mM) | *Sp*NanR (6 mg.mL$^{-1}$, ~185 μM) | *Sp*NanR (5 mg.mL$^{-1}$, ~150 μM) + *nan*E DNA oligo (2 mg.mL$^{-1}$, ~150 μM) |
| sample volume (μL) | 1 | 1 | 1 | 1 | 1 |
| crystallization temperature | 20 °C | 20 °C | 20 °C | 8 °C | 8 °C |
| crystallization method | hanging drop | sitting drop | sitting drop | sitting drop | sitting drop |

**Supplementary Table 8 | Heavy-atom derivative individual dataset statistics.**

| dataset | 0079 | 0080 | 0083 | 0084 | 0085 | 0086 | 0087 | merged |
|---|---|---|---|---|---|---|---|---|
| wavelength (Å) | 1.4586 | 1.4586 | 1.4586 | 1.4586 | 1.4586 | 1.4586 | 1.4586 | 1.4586 |
| total frames | 3600 | 3600 | 3600 | 3600 | 3600 | 3600 | 3600 | 3600x 7 |
| oscillation (°) | 0.10 | 0.10 | 0.10 | 0.10 | 0.10 | 0.10 | 0.10 | 0.10 |
| space group | 4 | 4 | 4 | 4 | 4 | 4 | 4 | 4 |
| unit cell (a,b,c,β) | 147.32, 84.35, 148.03, 90.15 | 147.28, 84.27, 148.19, 90.09 | 147.52, 84.44, 148.02, 90.13 | 147.60, 84.54, 148.05, 90.15 | 147.53, 84.56, 147.99, 90.20 | 147.48, 84.50, 148.04, 90.17 | 147.40, 84.43, 148.10, 90.13 | 147.32, 84.35, 148.03, 90.15 |
| resolution (Å) | 2.64 | 2.74 | 2.86 | 2.75 | 2.87 | 2.84 | 2.87 | 2.65 |
| mosaicity (°) | 0.072 | 0.076 | 0.079 | 0.078 | 0.092 | 0.088 | 0.089 | - |
| total reflections | 738,642/117,362 | 662,426/105,604 | 578,246/88,461 | 657,097/103,350 | 572,883/86,735 | 591,995/91,704 | 573,586/88,146 | 4,373,231/51,665 |
| unique reflections | 207,224/32,849 | 185,734/29,678 | 163,971/25,562 | 185,012/29,343 | 161726/25,227 | 166,844/26,402 | 161,484/25,475 | 207,452/14,687 |
| redundancy | 3.56/3.57 | 3.57/3.56 | 3.53/3.46 | 3.55/3.52 | 3.54/3.44 | 3.55/3.47 | 3.55/3.46 | 21.0/3.5 |
| completeness (%) | 99.5/97.8 | 99.6/98.5 | 99.1/95.6 | 99.5/97.7 | 99.1/95.6 | 99.4/97.2 | 99.3/97.0 | 99.6/95.3 |
| $I/\sigma(I)$ | 7.38/1.19 | 5.85/0.97 | 6.44/1.08 | 6.42/0.97 | 6.00/1.05 | 6.71/1.05 | 6.64/1.02 | 12.96/0.96 |
| $CC_{anom}$ (%) | 15/0 | 12/0 | 15/0 | 14/0 | 13/-1 | 13/1 | 13/1 | 31/0 |
| $R_{merge}$ (%) | 11.4/89.1 | 14.4/107.9 | 13.5/99.9 | 13.3/112.3 | 14.6/106.5 | 13.2/103.3 | 13.5/106.2 | 17.2/103.5 |
| $R_{meas}$ (%) | 13.5/104.8 | 17.0/127.0 | 15.9/118.2 | 15.7/132.4 | 17.3/126.3 | 15.5/122.3 | 15.9/125.7 | 17.6/122.0 |
| Wilson-B | 62.40 | 63.10 | 66.08 | 67.02 | 69.05 | 69.12 | 69.37 | 63.29 |

**Supplementary Table 9 | Summary of small angle scattering experimental setup and measured parameters.** Includes *Sp*NanR, *Sp*NanR + *N*-acetylmannosamine-6-phosphate (2 mM), and *Sp*NanR + *N*-acetylmannosamine-6-phosphate (2 mM) + *nan*E recognition DNA sequence, as determined by SEC-SAXS analysis shown in **Supplementary Figure 13** and **Supplementary Figure 16**. The protein concentration loaded onto the SEC column was 10 mg.mL$^{-1}$ and the SAXS data collected at an estimated concentration of ~2–3 mg.mL$^{-1}$.

| **SAXS data collection parameters** | | | |
|---|---|---|---|
| Instrument | Australian Synchrotron SAXS/WAXS beamline | | |
| Detector | PILATUS 1M (Dectris) | | |
| Wavelength (Å) | 1.07812 | | |
| Maximum flux at sample | 8 × $10^{12}$ photons per second at 12 keV | | |
| Camera length (mm) | 2,500 | | |
| q-range (Å$^{-1}$) | 0.006–0.55 | | |
| Exposure time | Continuous 1 s frame measurements | | |
| Sample configuration | SEC-SAXS with co-flow | | |
| Sample temperature (°C) | 20 | | |
| **SAXS data analysis** | *Sp*NanR | *Sp*NanR + manNAc-6-P | *Sp*NanR + DNA + manNAc-6-P |
| *Guinier analysis* | | | |
| $I_o$ (cm$^{-1}$) | 0.08 | 0.11 | 0.16 |
| $R_g$ (Å) | 38.7 ±0.1 | 40.6 ±0.1 | 47.3 ±0.1 |
| *P(r) analysis* | | | |
| $I_o$ (cm$^{-1}$) | 0.08 | 0.11 | 0.16 |
| $R_g$ (Å) | 40.7 | 42.4 | 47.6 |
| $D_{max}$ (Å) | ~165 | ~168 | ~178 |
| alpha | 0.2 | 0.1 | 0.4 |
| Porod volume (Å$^{-3}$) | 181,800 | 228,900 | 230,100 |
| $D_{max}$, *crystal structure* (Å) | 160 | 160 | 175 |
| *Mw (from Porod)* (Da) | 104,200 | 127,900 | 134,300 |

**Supplementary Table 10 | Conservation of the *Sp*NanR recognition sites.** Where the species-specific recognition site differs from the *S. pneumoniae* downstream gene, an arrow notes the correct downstream gene.

| **species/strain** | **promoter regions** | | | **cryptic sites** | |
|---|---|---|---|---|---|
| | **nan operon-I (*nan*E)** | **nan operon-II** | **s*ia*A operon** | ***nan*A** | **end of 0084** |
| *S. pneu.*/D39 | TCTGAAAGTACTTTTAGA | TTTGAAGAGATTTTCAAA | GCTGAAACTACTTTCAAG | TCTGAAACTACTTTCAAA | AGACTTTGATGAAAGTCT |
| *S. pneu.*/R36A | TCTGAAAGTACTTTTAGA | TTTGAAGAGATTTTCAAA | GCTGAAACTACTTTCAAG | TCTGAAACTACTTTCAAA | AGACTTTGATGAAAGTCT |
| *S. pneu.*/TIGR4 | TCTGAAAGTACTTTTAGA | TTTGAAGAGATTTTCAAA | GCTGAAACTACTTTCAAG | TCTGAAACTACTTTCAAA | AGACTTTGATGAAAGTCT |
| *S. mitis* | TCTGAAACTACTTTCAGA | | | TCTGAAACTACTTTCAAA | ←*nan*K[#] |
| *S. agalactiae* | TCTGAAACTACTTTCACG | | | | |
| *S. equi* | AATGAAAGTACTTCCTAA | | | AATGAAAGTAGTTTCTAA | ←*nan*E2* |
| *S. sanguinis* | CCTGAAAGTACTTTCAGG | | | | |
| *S. suis* | TCTGAAACTATTTTCAGG | | | | |
| *S. uberis* | TCTGAAAATAGTTTCAGA | | | | |

[#] "←nanK" denotes the actual downstream gene, *nanK*, that follows the recognition sequence in *S. mitis*.

* ←nanE2 denotes the actual downstream gene, *nanE2* (a second annotated *N*-acetylmannosamine-6-phosphate 2-epimerase), that follows the recognition sequence *in S. equi*.

### Supplementary Figures

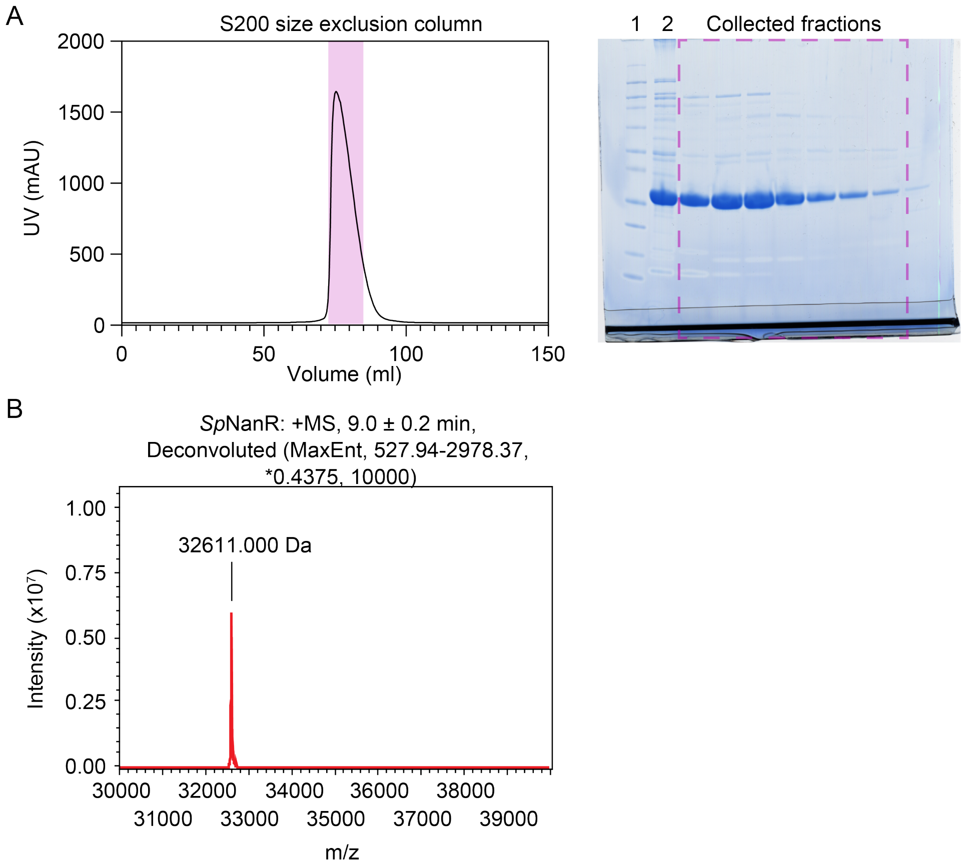


**Supplementary Figure 1 | SDS-PAGE analysis and mass spectrometry verifies that the sample was >95% pure and that the mass is as expected.** (**A**) Size-exclusion data (injected at ~20 mg.mL$^{-1}$, ~600 µM) indicates that *Sp*NanR forms multiple oligomeric species as it elutes as a highly asymmetric peak. The void volume for this column is ~45 mL. The shaded peak fractions were analyzed by SDS-PAGE (right) and contained *Sp*NanR. Lane 1 is the ladder and lane 2 represents what was loaded onto the SEC column. (**B**) Intact mass spectrometry verifies the presence of pure, full-length *Sp*NanR in the size-exclusion eluent fractions (theoretical molecular mass = 32,612.99, calculated molecular mass from spectra = 32,611.00.

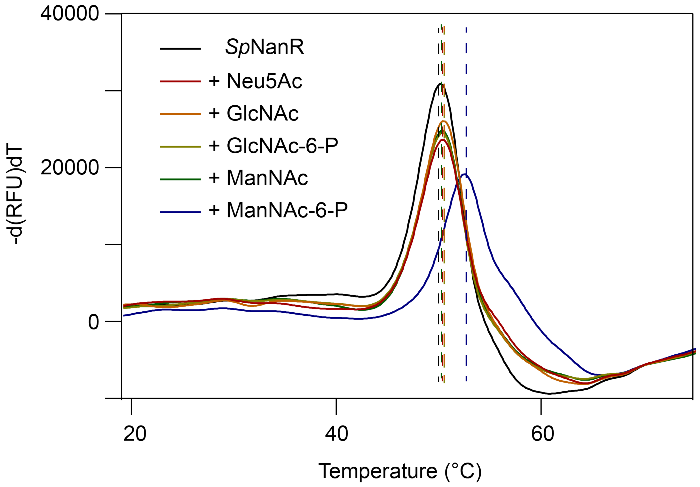


**Supplementary Figure 2 | Differential scanning fluorimetry.** Fluorescence as a function of temperature indicates protein stability when *Sp*NanR is incubated with each sialic acid derivative. Dotted lines corresponding to the peaks of each melt curve indicate the melting temperature. This shows the stabilizing effect of *N*-acetylmannosamine-6-phosphate (manNAc-6-P) on *Sp*NanR in comparison to *Sp*NanR alone and other sugars. Results are tabulated in **Supplementary Table 1** and plotted in **Figure 2A**.

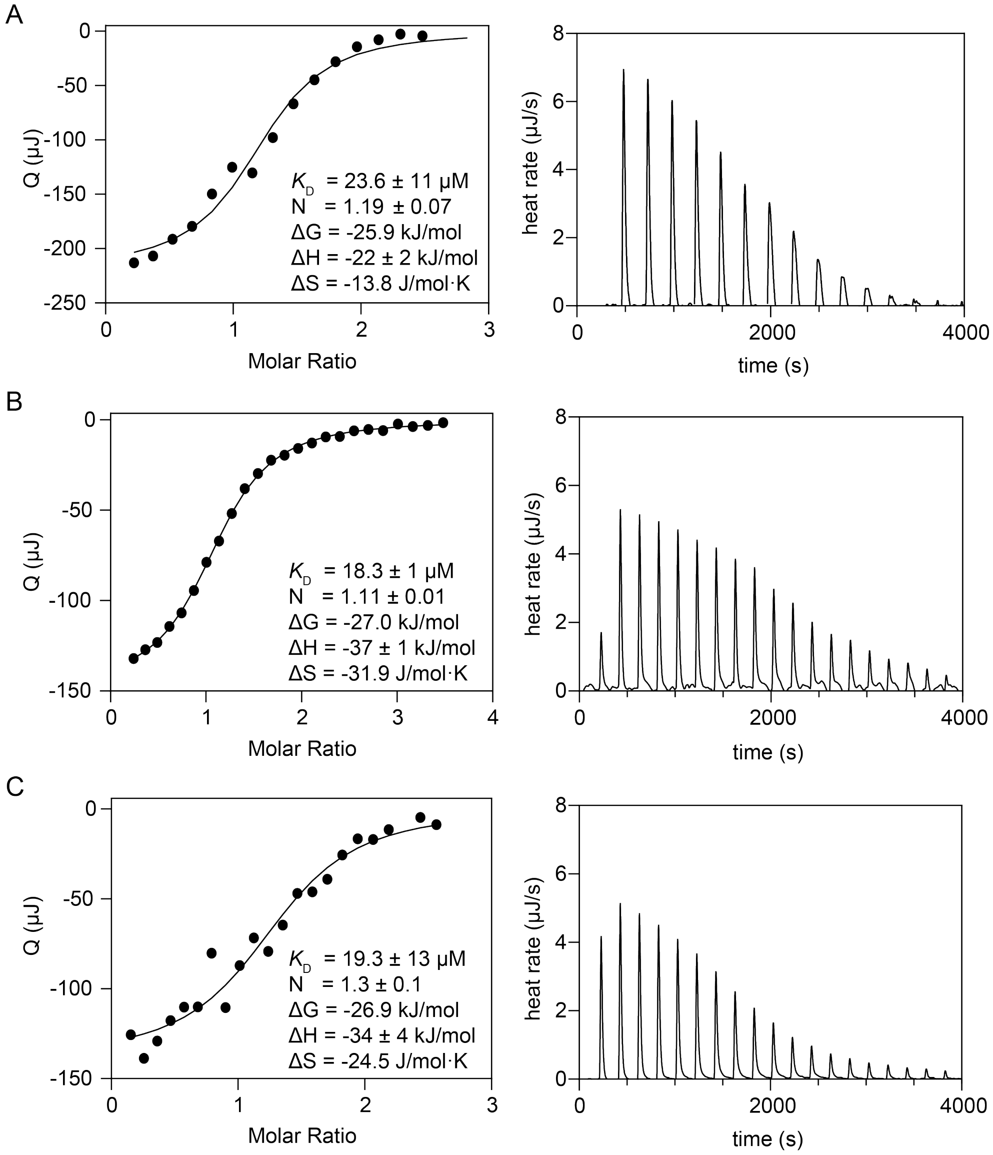


**Supplementary Figure 3 | Isothermal titration calorimetry.** (**A-C**) Three replicate experiments with the data fitted to an independent model and the fitted parameters inset. The error values represent the 95% confidence interval for each parameter.

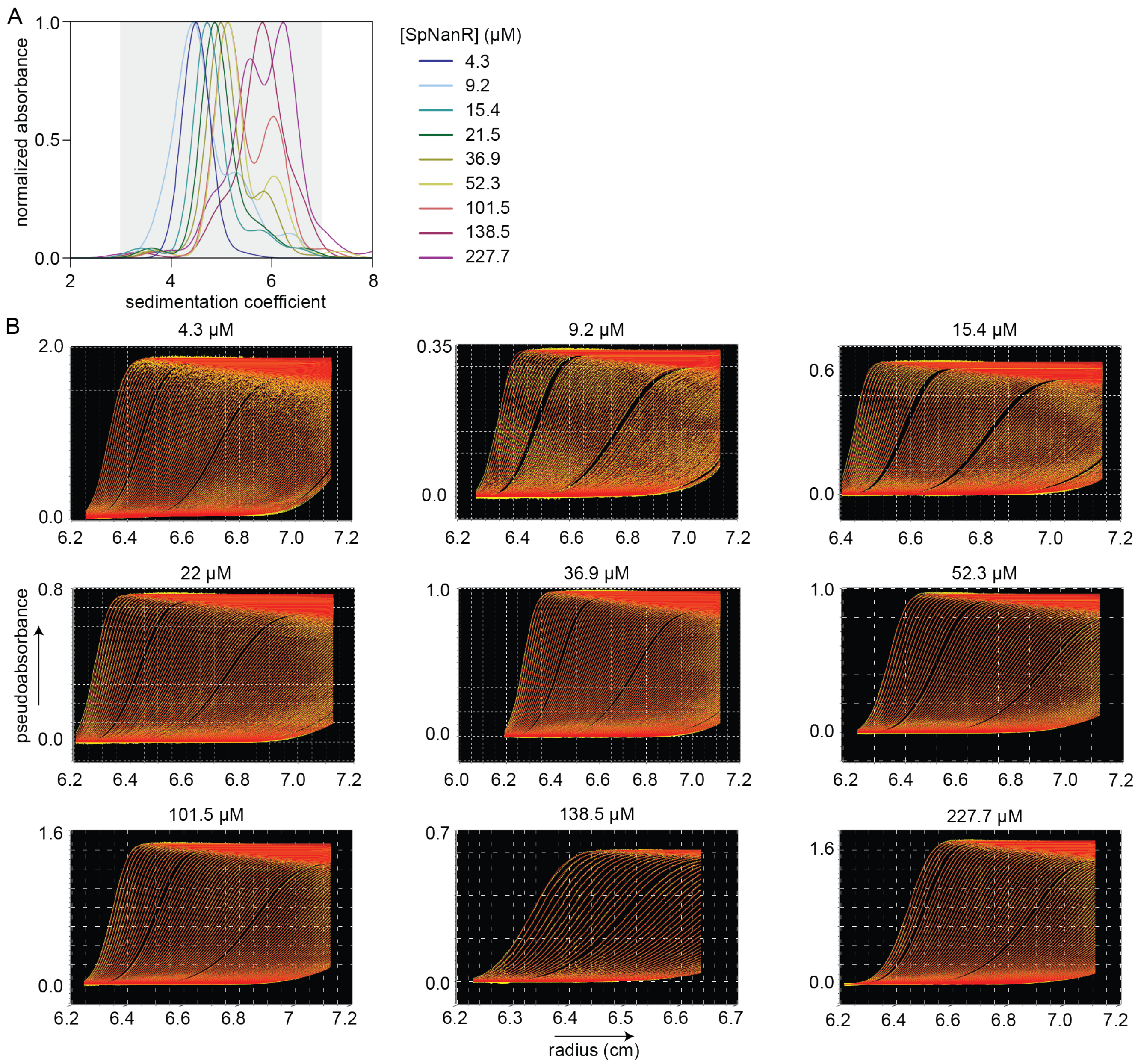


**Supplementary Figure 4 | 2DSA analysis of sedimentation velocity data for *Sp*NanR.** (**A**) The normalized sedimentation profiles across a range of *Sp*NanR concentrations (4.3–227.7 µM) as determined by analytical ultracentrifugation and analyzed with UltraScan 4.0 (4) using a 2DSA analysis method. Fit statistics are in **Supplementary Table 3**. The shaded area represents the S range used to determine the weight-averaged S plotted in **Figure 3B**, black circles. (**B**) Data and fits for the plots in **A**. Using UltraScan 4.0 (5), data from each sedimentation experiment was fit to a 2DSA-MC sedimentation model. The model (red) is fit to the experimental data (yellow). The residual r.m.s.d. is reported in **Supplementary Table 3** along with other statistical parameters.

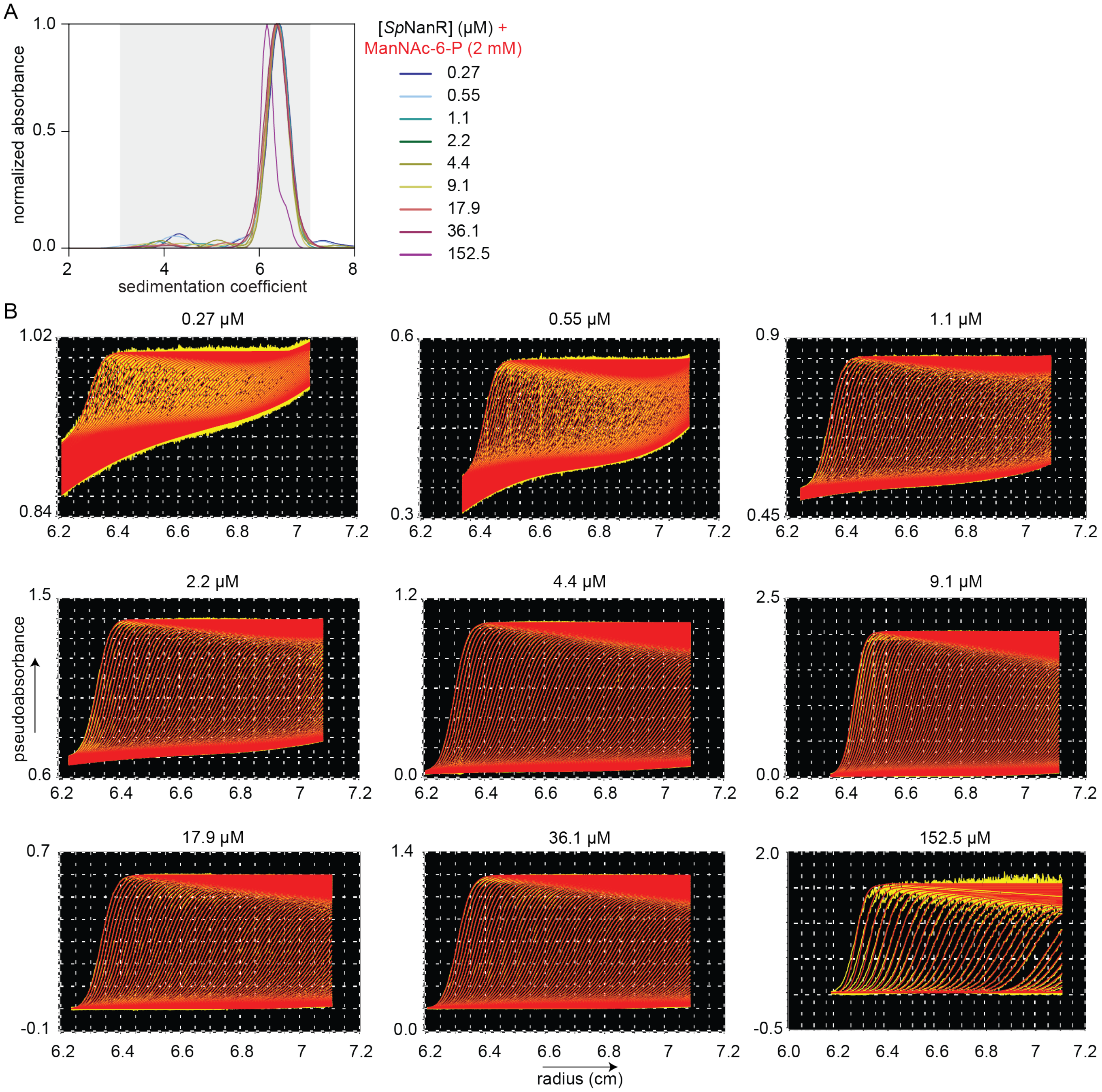


**Supplementary Figure 5 | 2DSA analysis of sedimentation velocity data for *Sp*NanR with *N*-acetylmannosamine-6-phosphate.** (**A**) The sedimentation profiles across a range of *Sp*NanR concentrations (0.27–152.3 µM) in the presence of *N*-acetylmannosamine-6-phosphate (2 mM) determined using analytical ultracentrifugation and analyzed with UltraScan 4.0 (4) and the 2DSA analysis method. The shaded area represents the S range used to determine the weight-averaged S plotted in **Figure 3B**, red squares. (**B**) Sedimentation velocity experiments of a concentration series of *Sp*NanR and 2 mM *N*-acetylmannosamine-6-phosphate. The sedimentation coefficients all exhibit similar results at ~6.2 S, calculated using UltraScan 2DSA analysis and iterative refinement. The data collected at 0.27–2.2 µM required lower wavelengths (220 and 228 nm) where the *N*-acetylmannosamine-6-phosphate containing buffer showed significant absorbance and gave a signal at ~0.2 S (not shown). The model (red lines) is fit to the experimental data (yellow lines). The residual r.m.s.d. is reported in **Supplementary Table 3** along with other statistical parameters.

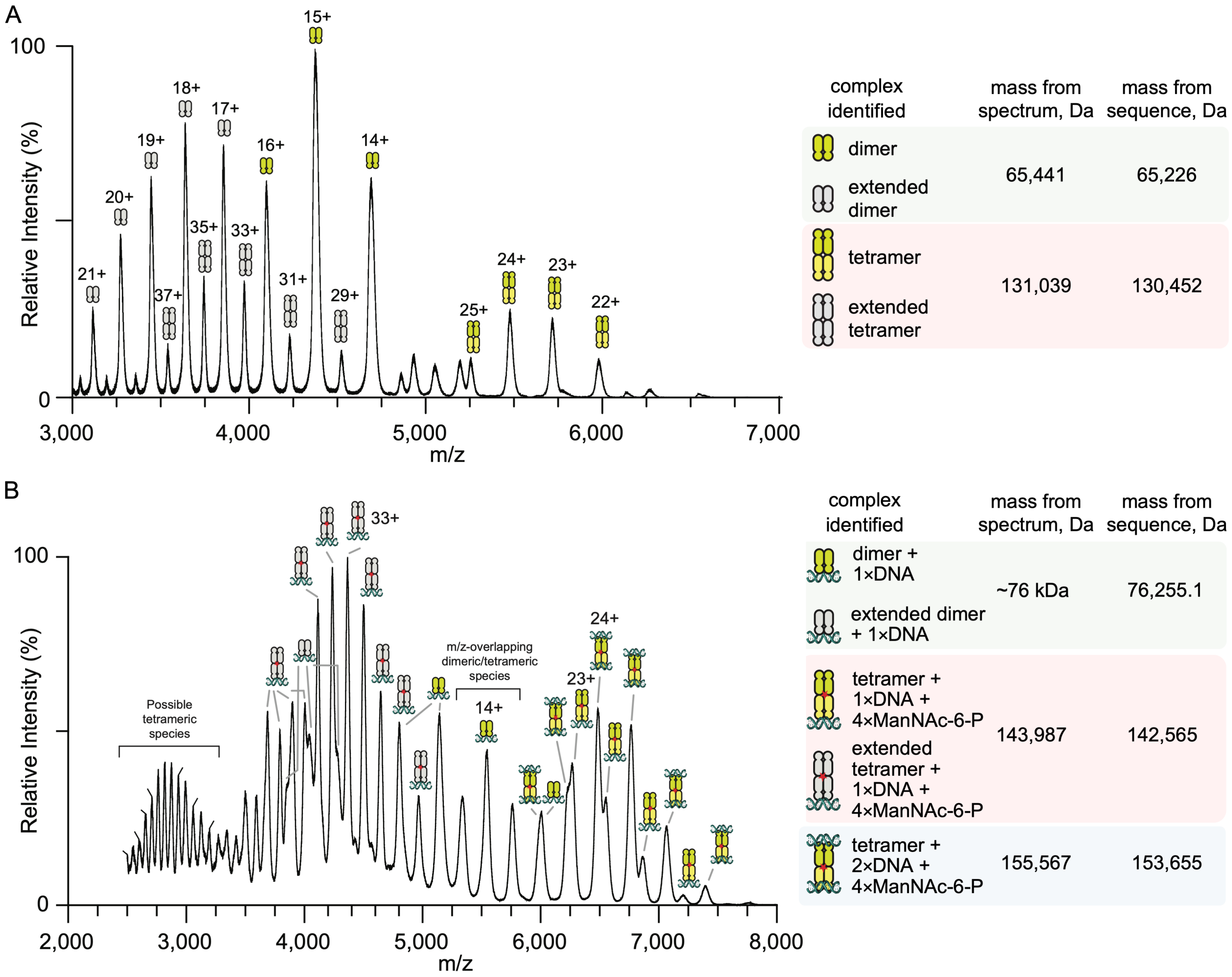


**Supplementary Figure 6 | Native mass spectrometry of *Sp*NanR.** (**A**) Analysis of charge states of *Sp*NanR protein (0.5 mg.mL$^{-1}$, 15 µM) in SEC buffer identifies both dimeric and tetrameric species. The presence of a separate population of higher charge states is indicative of an extended conformation, suggesting there is some flexibility in the protein (6,7). (**B**) Addition of DNA and *N*-acetylmannosamine-6-phosphate (manNAc-6-P) leads to a complex spectrum with multiple species. *Sp*NanR tetramers with one or two DNA oligos are evident, as are species with an extended conformation, suggesting that *Sp*NanR retains flexibility with DNA and manNAc-6-P. Analysis of charge states of *Sp*NanR protein (1.6 mg.mL$^{-1}$, 50 µM) first incubated with the 18 base pair *nan*E recognition sequence (0.5 mg.mL$^{-1}$, 50 µM, dsDNA $M_W$ = 10,990 Da) and then 10 mM manNAc-6-P ($M_W$ = 301.2 Da) identifies multiple species. The tables to the right show the calculated masses of the different complexes identified in the spectra, along with the masses calculated from the sequence.

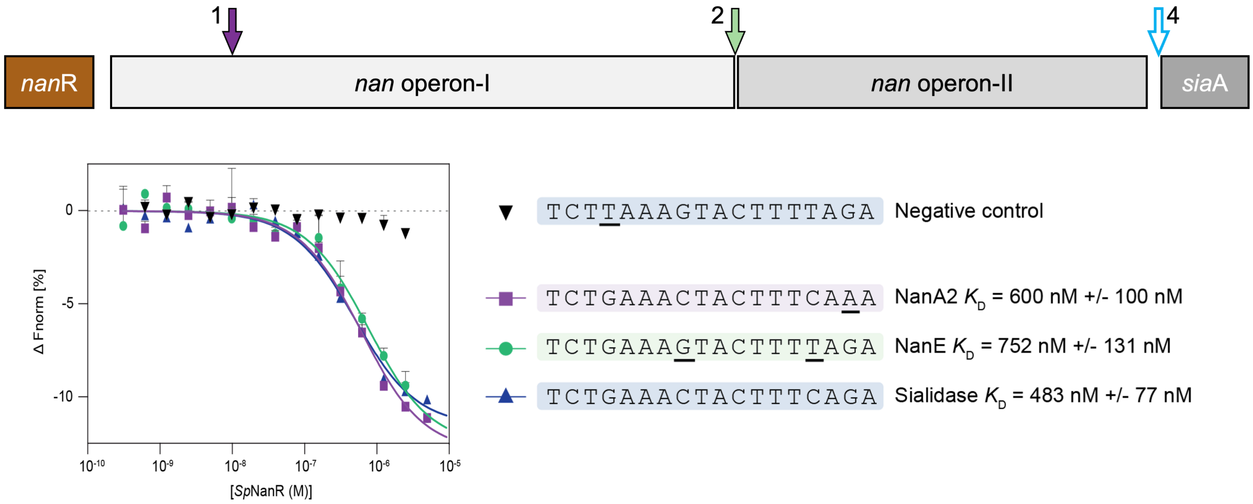


**Supplementary Figure 7 | *Sp*NanR affinity measurement for the three *Sp*NanR recognition sites in *S. pneumoniae* D39.** Location of the *Sp*NanR promoter sites (top) in the *nan* operon of *S. pneumoniae*. Microscale thermophoresis results that measure the binding of *Sp*NanR to the different DNA sequences (bottom). Dissociation constants ($K_D$) are reported. For all sequences in the *nan* operon (except the negative control), *Sp*NanR binds with similar affinity. The negative control is a mutated sequence (underlined) based on structural studies and found to not bind *Sp*NanR, discussed later.

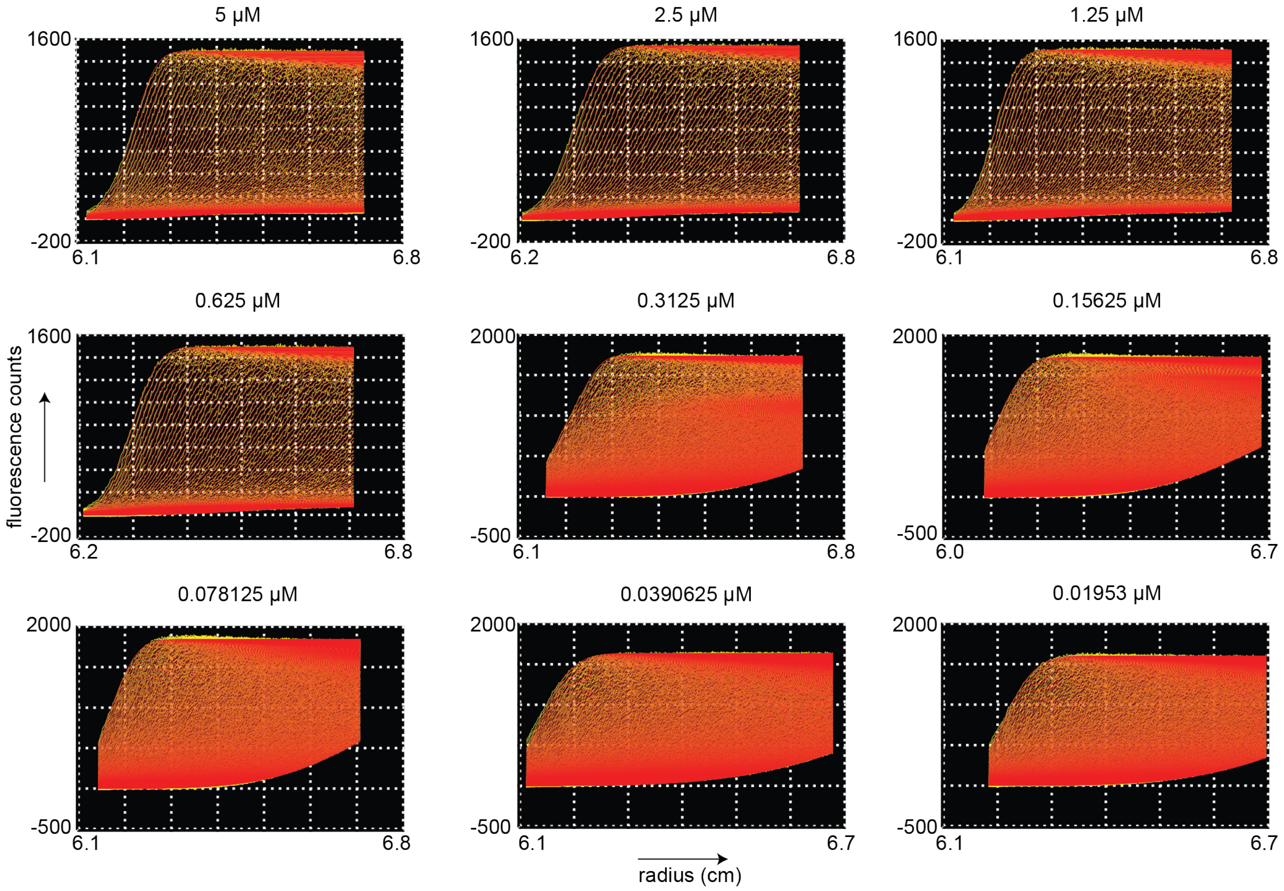


**Supplementary Figure 8 | 2DSA analysis of fluorescent sedimentation velocity data for *Sp*NanR and FAM-labeled *nan*E recognition DNA sequence.** Data and fits for the plots in **Figure 4A**, left. Using UltraScan 4.0 (5), data from each sedimentation experiment was fit to a 2DSA sedimentation model. The model (red) is fit to the experimental data (yellow). The residual r.m.s.d. is reported in **Supplementary Table 5** along with other statistical parameters.

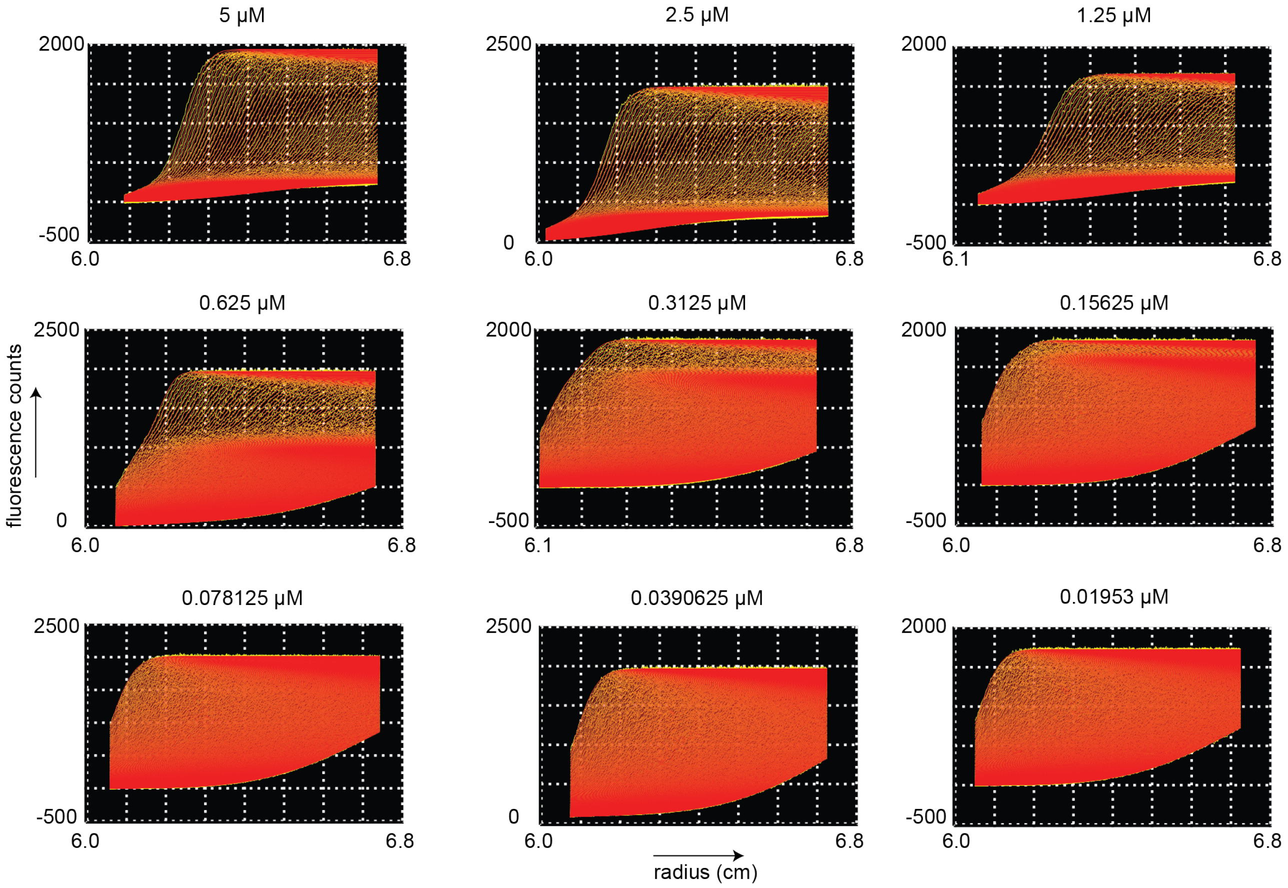


**Supplementary Figure 9 | 2DSA analysis of fluorescent sedimentation velocity data for *Sp*NanR and FAM-labeled *nan*E recognition DNA sequence in the presence of *N*-acetylmannosamine-6-phosphate.** Data and fits for the plots in **Figure 4A**, right. Using UltraScan 4.0 (5), data from each sedimentation experiment was fit to a 2DSA sedimentation model. The model (red) is fit to the experimental data (yellow). The residual r.m.s.d. is reported in **Supplementary Table 5** along with other statistical parameters.

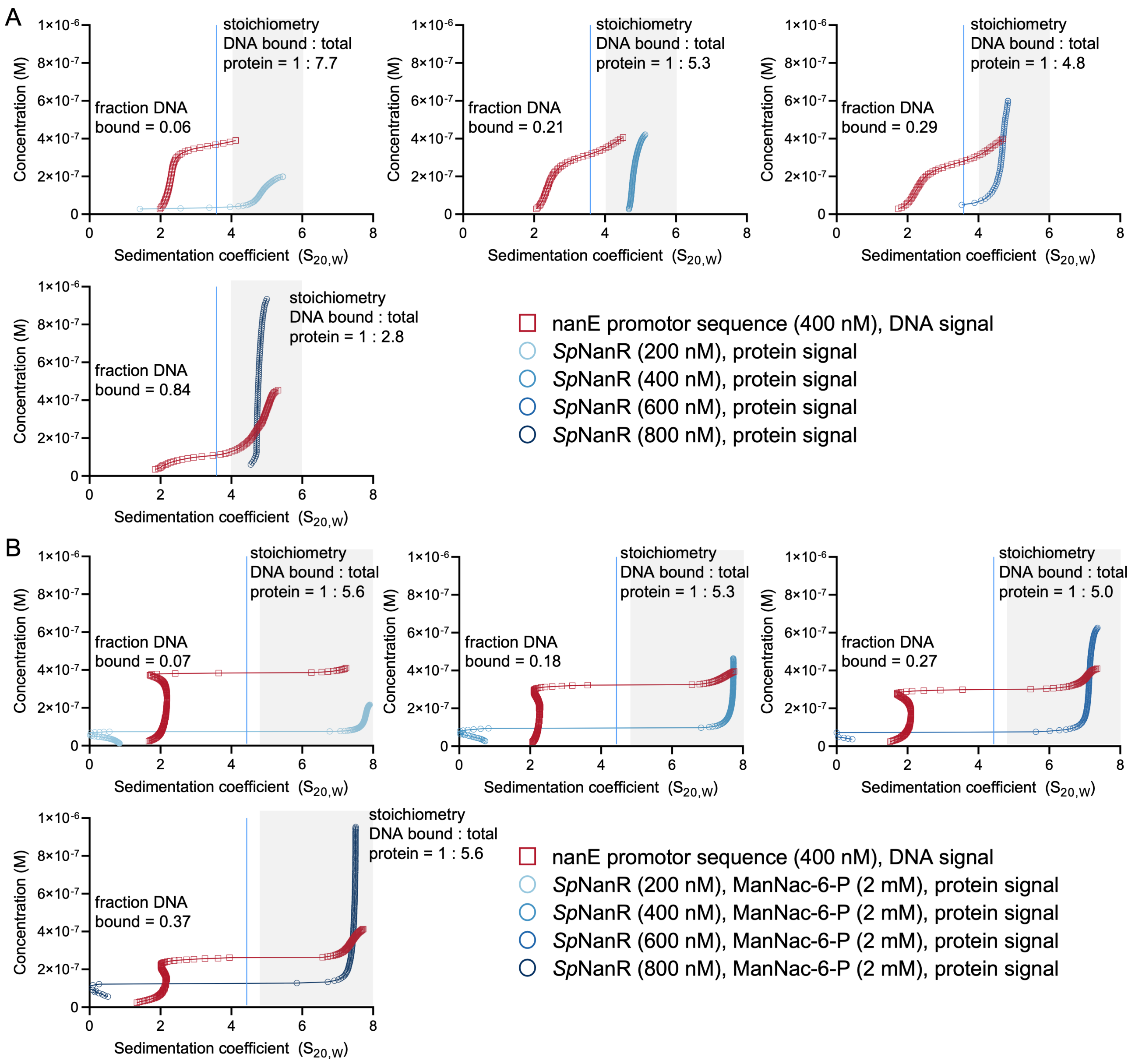


**Supplementary Figure 10 | Multiwavelength sedimentation velocity data verifies that *Sp*NanR binds the *nan*E recognition DNA sequence.** (**A**) van Holde-Weischet plots for the titration of *Sp*NanR (200–800 nM) against a constant concentration of the *nanE* recognition DNA sequence (400 nM). The plots were generated using deconvoluted multiwavelength sedimentation velocity data with UltraScan 4.0 (5), see Methods and Materials. The shaded area shows the co-sedimenting DNA and *Sp*NanR. (**B**) This experiment is as in A but includes 2 mM *N*-acetylmannosamine-6-phosphate. There was significant absorbance by *N*-acetylmannosamine-6-phosphate leading to the van Holde-Weischet plots showing a peak at ~0.1 S in the protein signal.

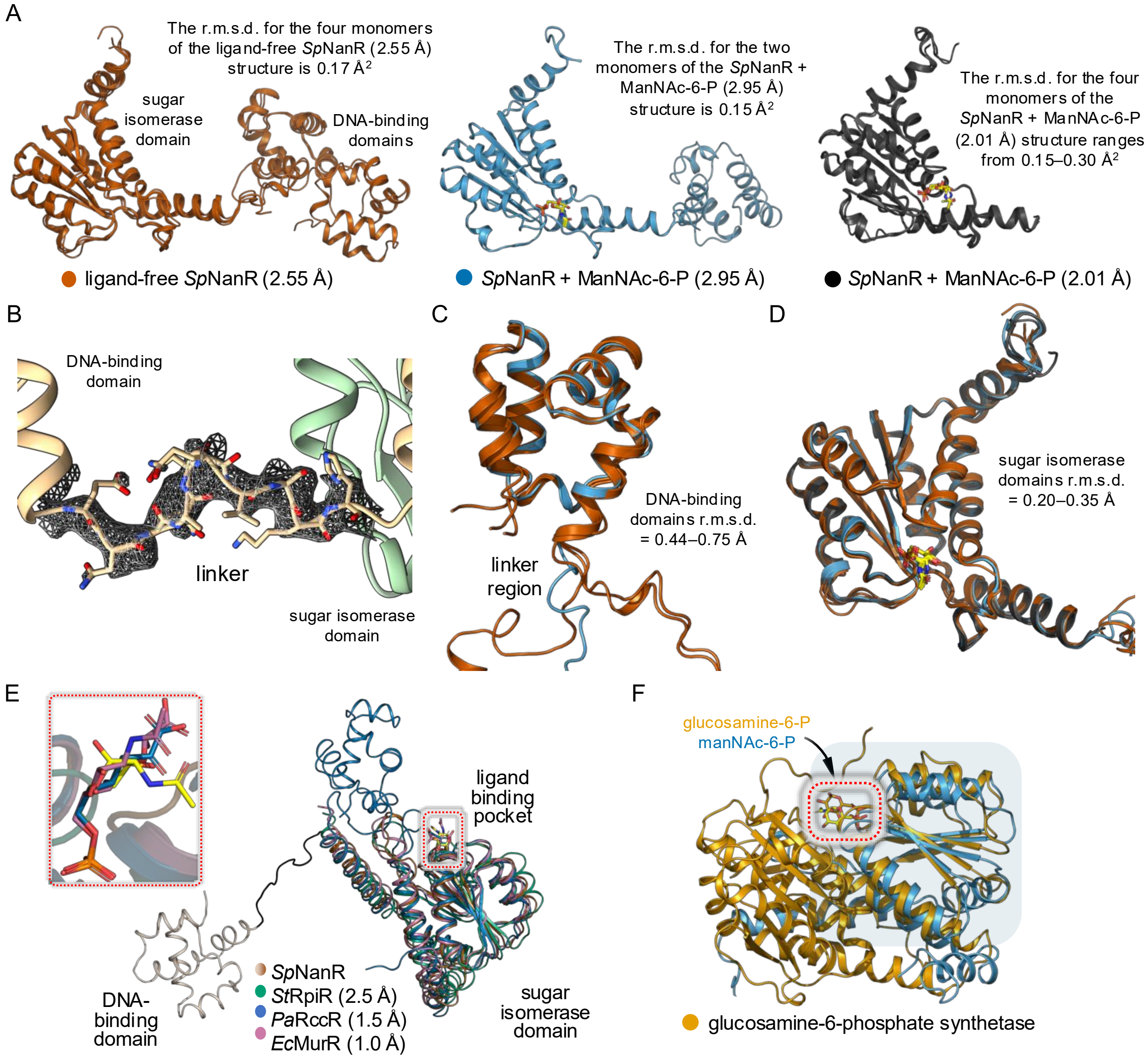


**Supplementary Figure 11 | Domain structure of *Sp*NanR.** (**A**) Overlay of the sugar isomerase domains from the three different structures of *Sp*NanR (labeled below). (**B**) Electron density ($2F_0$-$F_c$ contoured to 1.0) for the linker sequence in the apo *Sp*NanR structure. (**C**) Overlay of DNA binding domains and linker sequences from the apo and *N*-acetylmannosamine-6-phosphate bound *Sp*NanR show no significant changes in the structure of this domain. (**D**) Sugar isomerase domains from the three reported structures in A were aligned to demonstrate the strict similarity between them all (r.m.s.d. = 0.20–0.35 Å$^2$). (**E**) Aligned sugar isomerase domains illustrate the close similarity between the three transcriptional regulators homologues [*Sp*NanR monomer bound with *N*-acetylmannosamine-6-phosphate (brown), *St*RpiR (PDB ID: 3SHO, teal, r.m.s.d. = 2.5 Å), *Pa*RccR (PDB ID: 8K3B, blue, r.m.s.d = 1.5 Å), and *Ec*MurR (PDB ID: 7EN6, salmon, r.m.s.d. = 1.0 Å)]. Inset, the phosphate moieties of the bound ligands overlay very well. (**F**) The sugar isomerase domain of *Sp*NanR bound to manNAc-6-P was aligned to glucosamine-6-phosphate synthetase (PDB ID: 1MOR), thought to be an evolutionary precursor for the sugar isomerase domain of the RpiR family, and the domains align very well (r.m.s.d. Cα atoms = 2.6 Å$^2$).

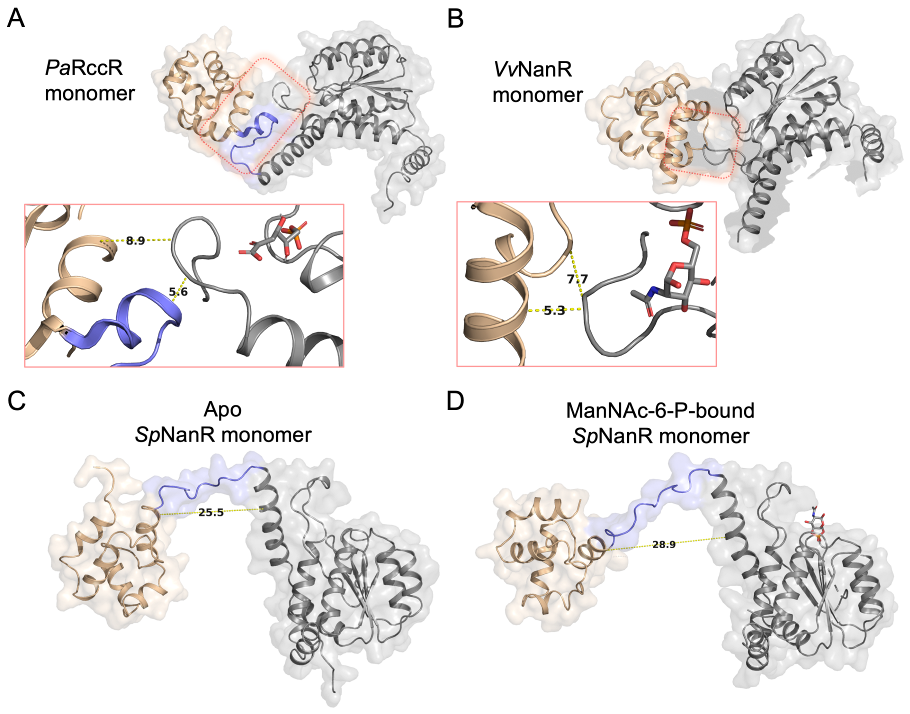


**Supplementary Figure 12 | Domain assembly of *Sp*NanR and RpiR-family homologues.** The monomeric structures of other RpiR-family transcriptional regulators compared with *Sp*NanR. Highlighted with color are the DNA-binding domain (beige) and linker sequence (purple) for each monomer to illustrate their position in relation to the sugar isomerase domain. (**A**) The monomer from the *Pa*RccR crystal structure (PDB ID: 8JU9) is depicted, linker sequence highlighted in purple which forms contacts with a flexible motif within the sugar isomerase domain. (**B**) The monomer corresponding to the *Vv*NanR crystal structure (PDB ID: 4IVN) is depicted; the linker sequence was not modeled due to a lack of density, but the DNA-binding domain exhibits multiple contacts with the sugar isomerase domain. (**C**) The apo *Sp*NanR monomer is depicted, linker sequence highlighted in beige. (**D**) The *N*-acetylmannosamine-6-phosphate bound *Sp*NanR monomer is depicted, linker sequence highlighted in beige.

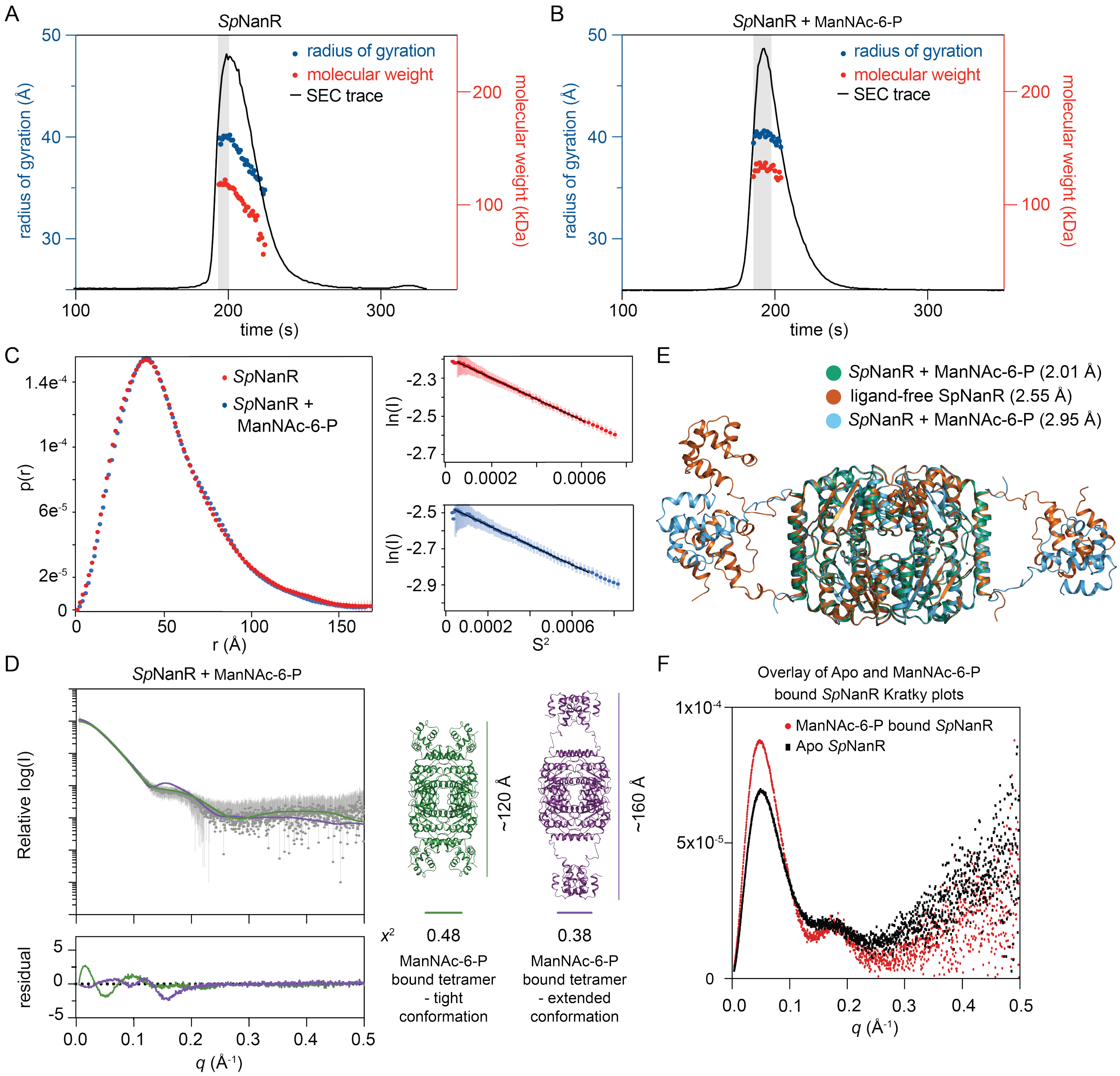


**Supplementary Figure 13 |** Small angle X-ray scattering **of *Sp*NanR.** (**A**, **B**) In-line size exclusion chromatography (SEC) small angle X-ray scattering (SAXS) demonstrates that without manNAc-6-P the molecular weight and radius of gyration decrease, consistent with the dimer-tetramer self-association. With manNAc-6-P (2 mM) the molecular weight and radius of gyration is flat with a mass of ~130 kDa. Data used for subsequent analysis is shaded gray. (**C**) Pairwise distance distributions demonstrate that the maximum distance with or without manNAc-6-P is ~165 Å. Guinier plots show the data is of high quality (right-hand side). (**D**) SAXS profiles for *Sp*NanR and manNAc-6-P (2 mM). The scattering data is best fit (purple line, $\chi^2 = 0.38$) to the theoretical scatter of the extended tetramer over the tight tetramer conformation (green line, $\chi^2 = 0.48$). Residuals for the fits, generated by CRYSOL are shown below the SAXS profiles. Models of each *Sp*NanR conformation are shown. (**E**) Overlay of each crystal structure show the sugar isomerase domains align well (r.m.s.d. = 0.30–0.46 (across 654–670 atoms), suggesting ManNAc-6-P does not alter the tetrameric conformation. Though, the DNA-binding domains adopt an extended and variable conformation. (**F**) Kratky plot for the ligand-free and manNAc-6-P bound *Sp*NanR suggests the protein is multidomain and flexible, consistent notion that the DNA binding domains are flexible around the tetrameric sugar isomerase domain.

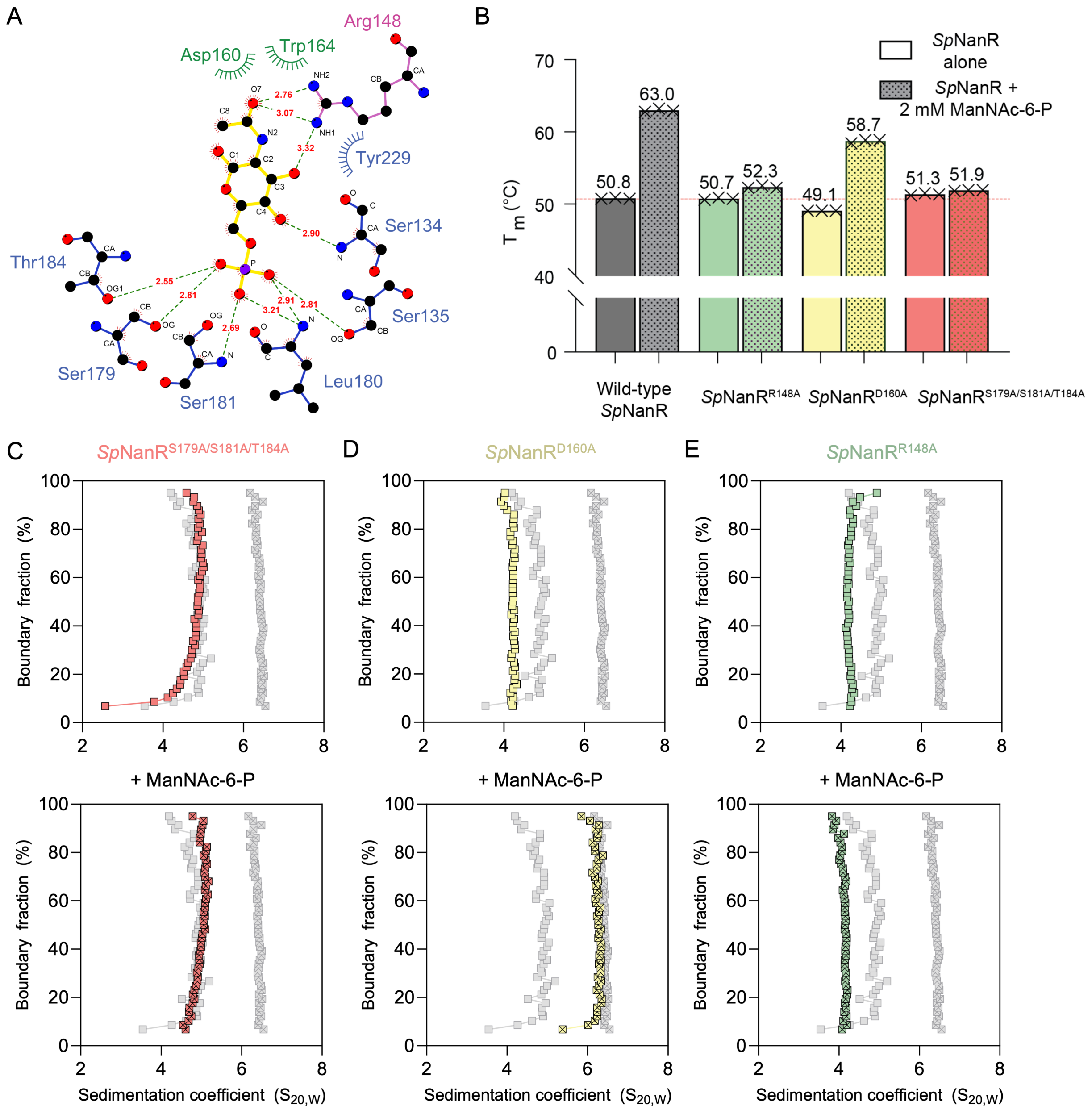


**Supplementary Figure 14 | Interactions between *Sp*NanR and *N*-acetylmannosamine-6-phosphate.** (**A**) LigPlot showing how *N*-acetylmannosamine-6-phosphate interacts with *Sp*NanR. Here, the *N*-acetylmannosamine-6-phosphate molecule, bound to monomer C in PDB ID: 8TX9, was chosen for visualization. In the three other *N*-acetylmannosamine-6-phosphate molecules in 8TX9 there are small differences in contacts. **Supplementary Table 6** reports the average interacting distances for the four *N*-acetylmannosamine-6-phosphate molecules. The dotted green lines indicated hydrogen bond interactions, while atoms with red lines indicate van der Waals interactions. (**B**) Differential scanning fluorimetry results of the thermal stability of each purified mutant protein that was tested without (left bar) and with (right bar) how *N*-acetylmannosamine-6-phosphate (manNAc-6-P). The gray bars represent wild-type *Sp*NanR thermal stability for reference. (**C-E**) Sedimentation velocity experiments assess the dimer-tetramer self-association of the mutant proteins (15 μM) without (top) and with 2 mM *N*-acetylmannosamine-6-phosphate (manNAc-6-P, bottom). The gray curves represent the sedimentation of wild-type *Sp*NanR in the same buffer and protein concentration (15 μM).

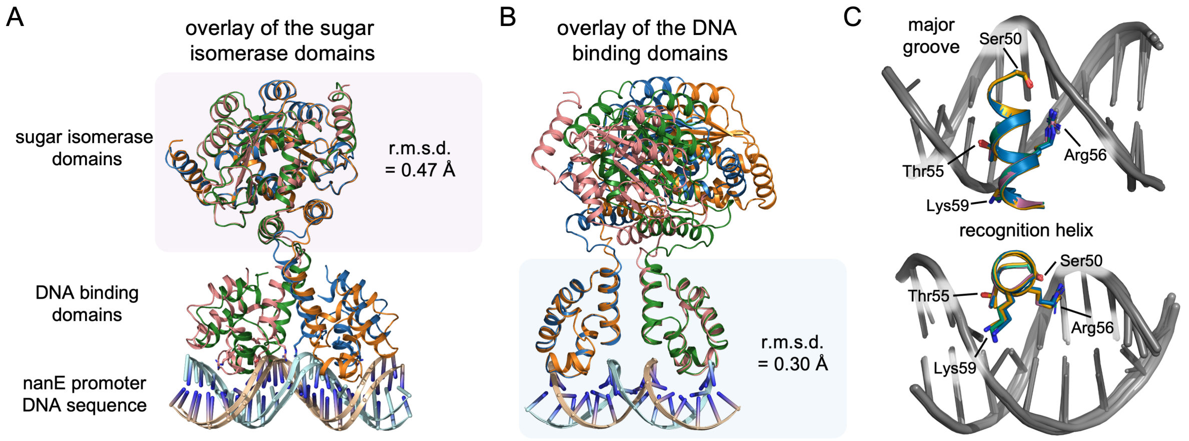


**Supplementary Figure 15 | Overlays of the two dimers that form the asymmetric unit in the *Sp*NanR + *nan*E recognition DNA complex crystal structure.** (**A**) The two sugar isomerase domains align very well (r.m.s.d. = 0.47 Å), whereas the entire dimer is much less (r.m.s.d. = 2.19). (**B**) When the DNA-binding domains are aligned, they align well (r.m.s.d. = 0.30 Å) and demonstrate very similar binding conformations with the *nanE* recognition DNA sequence, whereas now the sugar isomerase domains do not overlay well. (**C**) The interaction between the major DNA groove and the recognition helix.

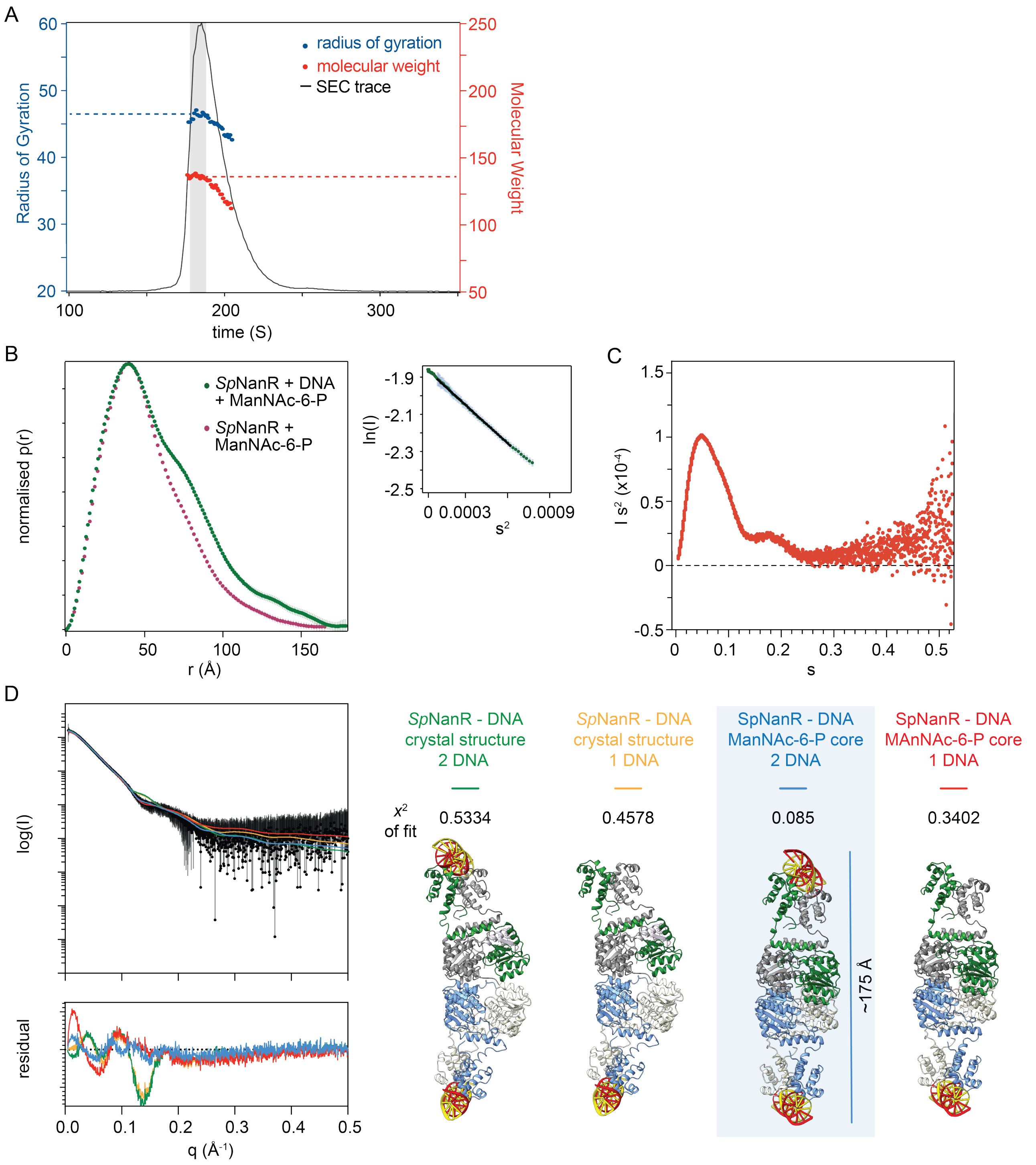


**Supplementary Figure 16 | Small angle X-ray scattering analysis of the *Sp*NanR + *nan*E recognition sequence + manNAc-6-P complex.** (**A**) In-line SEC-SAXS demonstrates that with manNAc-6-P and the *nan*E recognition sequence the molecular weight and radius of gyration decrease, consistent with the dimer tetramer self-association. Data used for subsequent analysis is shaded gray. (**B**) Pairwise distribution (*P*(*r*)) alignments for *Sp*NanR + manNAc-6-P, with (green) and without (pink) the *nan*E recognition sequence. Guinier plots show the data is of high quality (right-hand side). (**C**) Kratky plot for the *Sp*NanR + *nan*E recognition sequence + manNAc-6-P complex scattering data suggests the protein is multidomain and flexible, consistent notion that the DNA binding domains when complexed with DNA are flexible. (**D**) Theoretical SAXS scattering curves for the four PDB models are aligned to experimental SAXS data or *Sp*NanR + *nan*E recognition sequence + manNAc-6-P. Error estimates for the alignments can be seen below the models and the coloring scheme matches the model text. Models based on the crystal structure of

*Sp*NanR + *nan*E recognition sequence were designed to compare the theoretical scattering of a *Sp*NanR tetramer bound to one DNA (orange and red), and two DNAs (green and blue). In the first two models (green and orange), the asymmetric core tetramer of sugar isomerase domains is that of the native crystal DNA bound structure. In the second two models (blue and red), the core tetramer of sugar isomerase domains is that of the symmetric and homogenous manNAc-6-P bound *Sp*NanR tetramer. The chi-square ($X^2$) values for the CRYOSOL alignments are reported for each structure. The residuals demonstrate a very good fit between the *Sp*NanR structure in complex with manNAc-6-P and 2 DNA molecules, and the Chi-square fit value of 0.085 verifies this.

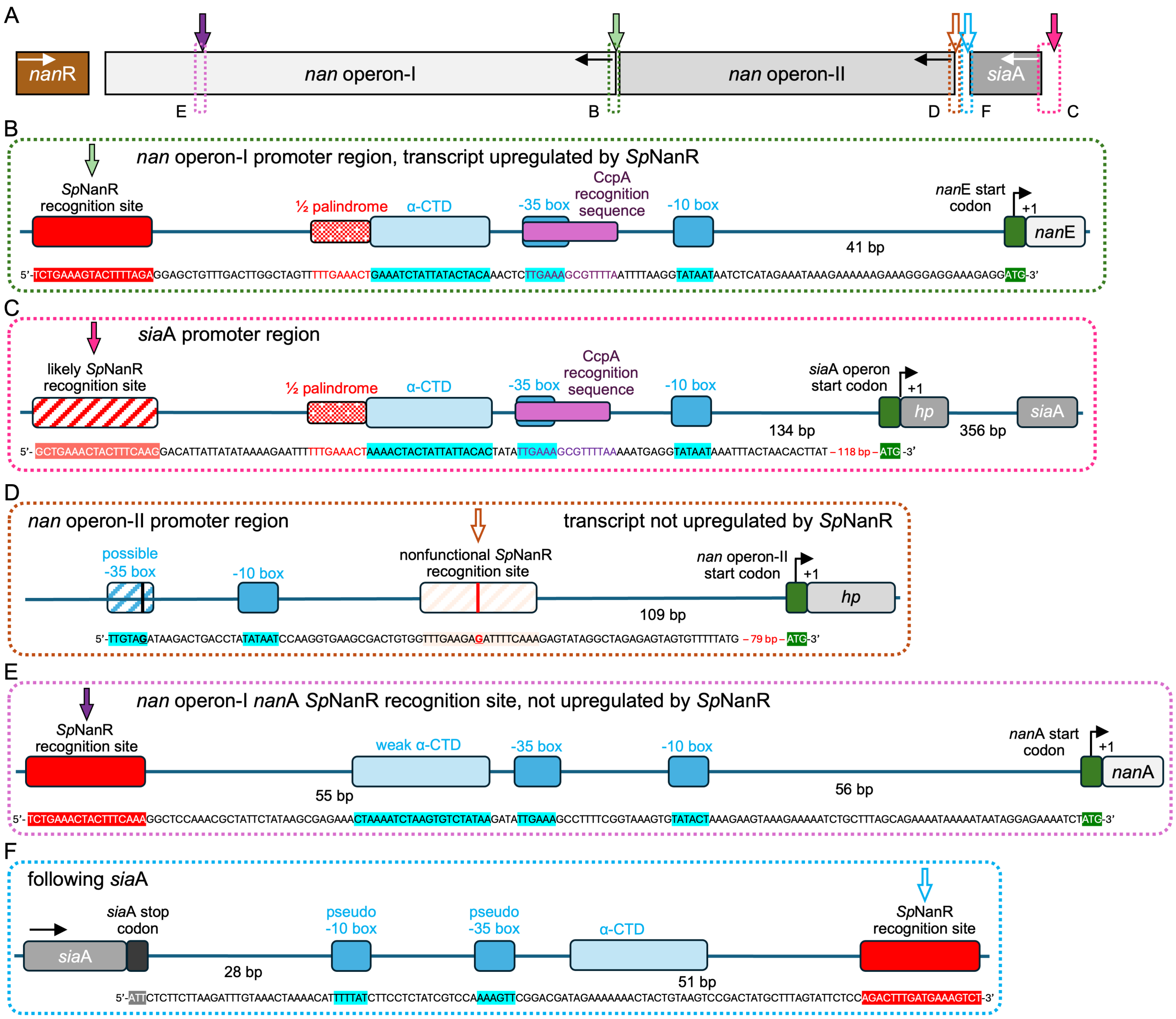


**Supplementary Figure 17 | Promoter regions and *Sp*NanR binding sites across the three *S. pneumoniae* D39 *nan* operons.** (**A**) Schematic of the three *nan* operons for reference. (**B**) Promoter region for *nan* operon-I showing the *Sp*NanR recognition sequence (solid red fill) and its juxtaposition to other regulatory and transcription elements; CcpA recognition sequence (purple), α-CTD binding site (light blue), -35 and -10 boxes (blue), and the translation start site (green). (**C**) Close inspection of the *sia*A operon identifies a possible functional *Sp*NanR recognition sequence and transcription elements (α-CTD binding site (UP), -35 and -10 boxes) ~200 base pairs upstream of the start codon. The first gene in *sia*A operon is a hypothetical protein, annotated *spd_1505*. (**D**) Promoter region for *nan* operon-II showing a potential *Sp*NanR recognition sequence following the -10 box. This sequence has a mutation (highlighted by the red line) that we demonstrate abolishes *Sp*NanR binding in the *nan*E recognition sequence (**Figure 6D**, open red box), so this site likely does not bind *Sp*NanR. The promoter region also has a possible transcription -35 box element. Mutations that lead to low-to-zero affinity are underlined. The first gene in *nan* operon-II is a hypothetical protein, annotated *spd_1503*. (**E**) Promoter region within *nan* operon-I prior to the *nan*A gene. (**F**) *Sp*NanR recognition sequence following the *sia*A operon, including pseudo transcriptional elements (blue).

### Supplementary References


1. Zhu, Y., Mou, X., Song, Y., Zhang, Q., Sun, B., Liu, H., Tang, H. and Bao, R. (2024) Molecular mechanism of the one-component regulator RccR on bacterial metabolism and virulence. *Nucleic Acids Res*, **52**, 3433-3449.
2. Hwang, J., Kim, B.S., Jang, S.Y., Lim, J.G., You, D.J., Jung, H.S., Oh, T.K., Lee, J.O., Choi, S.H. and Kim, M.H. (2013) Structural insights into the regulation of sialic acid catabolism by the Vibrio vulnificus transcriptional repressor NanR. *Proc Natl Acad Sci U S A*, **110**, E2829-2837.
3. Zhang, Y., Chen, W., Wu, D., Liu, Y., Wu, Z., Li, J., Zhang, S.Y. and Ji, Q. (2022) Molecular basis for cell-wall recycling regulation by transcriptional repressor MurR in Escherichia coli. *Nucleic Acids Res*, **50**, 5948-5960.
4. Demeler, B. and Gorbet, G.E. (2016) In Uchiyama, S., Arisaka, F., Stafford, W. F. and Laue, T. (eds.), *Analytical Ultracentrifugation: Instrumentation, Software, and Applications*. Springer Japan, Tokyo, pp. 119-143.
5. Demeler, B. (2024) Methods for the Design and Analysis of Analytical Ultracentrifugation Experiments. *Curr Protoc*, **4**, e974.
6. Natalello, A., Santambrogio, C. and Grandori, R. (2017) Are Charge-State Distributions a Reliable Tool Describing Molecular Ensembles of Intrinsically Disordered Proteins by Native MS? *J Am Soc Mass Spectrom*, **28**, 21-28.
7. Testa, L., Brocca, S., Santambrogio, C., D'Urzo, A., Habchi, J., Longhi, S., Uversky, V.N. and Grandori, R. (2013) Extracting structural information from charge-state distributions of intrinsically disordered proteins by non-denaturing electrospray-ionization mass spectrometry. *Intrinsically Disord Proteins*, **1**, e25068.
